# Evolution of cellular diversity in primary motor cortex of human, marmoset monkey, and mouse

**DOI:** 10.1101/2020.03.31.016972

**Authors:** Trygve E. Bakken, Nikolas L. Jorstad, Qiwen Hu, Blue B. Lake, Wei Tian, Brian E. Kalmbach, Megan Crow, Rebecca D. Hodge, Fenna M. Krienen, Staci A. Sorensen, Jeroen Eggermont, Zizhen Yao, Brian D. Aevermann, Andrew I. Aldridge, Anna Bartlett, Darren Bertagnolli, Tamara Casper, Rosa G. Castanon, Kirsten Crichton, Tanya L. Daigle, Rachel Dalley, Nick Dee, Nikolai Dembrow, Dinh Diep, Song-Lin Ding, Weixiu Dong, Rongxin Fang, Stephan Fischer, Melissa Goldman, Jeff Goldy, Lucas T. Graybuck, Brian R. Herb, Xiaomeng Hou, Jayaram Kancherla, Matthew Kroll, Kanan Lathia, Baldur van Lew, Yang Eric Li, Christine S. Liu, Hanqing Liu, Jacinta D. Lucero, Anup Mahurkar, Delissa McMillen, Jeremy A. Miller, Marmar Moussa, Joseph R. Nery, Philip R. Nicovich, Joshua Orvis, Julia K. Osteen, Scott Owen, Carter R. Palmer, Thanh Pham, Nongluk Plongthongkum, Olivier Poirion, Nora M. Reed, Christine Rimorin, Angeline Rivkin, William J. Romanow, Adriana E. Sedeño-Cortés, Kimberly Siletti, Saroja Somasundaram, Josef Sulc, Michael Tieu, Amy Torkelson, Herman Tung, Xinxin Wang, Fangming Xie, Anna Marie Yanny, Renee Zhang, Seth A. Ament, M. Margarita Behrens, Hector Corrada Bravo, Jerold Chun, Alexander Dobin, Jesse Gillis, Ronna Hertzano, Patrick R. Hof, Thomas Höllt, Gregory D. Horwitz, C. Dirk Keene, Peter V. Kharchenko, Andrew L. Ko, Boudewijn P. Lelieveldt, Chongyuan Luo, Eran A. Mukamel, Sebastian Preissl, Aviv Regev, Bing Ren, Richard H. Scheuermann, Kimberly Smith, William J. Spain, Owen R. White, Christof Koch, Michael Hawrylycz, Bosiljka Tasic, Evan Z. Macosko, Steven A. McCarroll, Jonathan T. Ting, Hongkui Zeng, Kun Zhang, Guoping Feng, Joseph R. Ecker, Sten Linnarsson, Ed S. Lein

## Abstract

The primary motor cortex (M1) is essential for voluntary fine motor control and is functionally conserved across mammals. Using high-throughput transcriptomic and epigenomic profiling of over 450,000 single nuclei in human, marmoset monkey, and mouse, we demonstrate a broadly conserved cellular makeup of this region, whose similarity mirrors evolutionary distance and is consistent between the transcriptome and epigenome. The core conserved molecular identity of neuronal and non-neuronal types allowed the generation of a cross-species consensus cell type classification and inference of conserved cell type properties across species. Despite overall conservation, many species specializations were apparent, including differences in cell type proportions, gene expression, DNA methylation, and chromatin state. Few cell type marker genes were conserved across species, providing a short list of candidate genes and regulatory mechanisms responsible for conserved features of homologous cell types, such as the GABAergic chandelier cells. This consensus transcriptomic classification allowed the Patch-seq identification of layer 5 (L5) corticospinal Betz cells in non-human primate and human and characterization of their highly specialized physiology and anatomy. These findings highlight the robust molecular underpinnings of cell type diversity in M1 across mammals and point to the genes and regulatory pathways responsible for the functional identity of cell types and their species-specific adaptations.

## Introduction

Single-cell transcriptomic and epigenomic methods provide a powerful lens on understanding the cellular makeup of highly complex brain tissues based on distinct patterns of gene expression and underlying regulatory mechanisms ^1–7^. Applied to mouse and human neocortex, single-cell or single-nucleus transcriptomic analysis has yielded a complex but finite classification of cell types with approximately 100 discriminable neuronal and non-neuronal types in any given neocortical region ^1, 2, 6, 8^. Similar analyses using epigenomic methods have shown that many cortical cell types can be distinguished on the basis of regions of open chromatin or DNA methylation ^5, 9, 10^. Furthermore, several recent studies have shown that transcriptomically-defined cell types can be aligned across species ^2, 11–13^, indicating that these methods provide a path to quantitatively study evolutionary conservation and divergence at the level of cell types. However, application of these methods has been highly fragmented to date. Human and mouse comparisons have been performed in different cortical regions, using single-cell (with biases in cell proportions) versus single-nucleus (with biases in transcript makeup) analysis, and most single-cell transcriptomic and epigenomic studies have been performed independently.

The primary motor cortex (MOp in mouse, M1 in human and non-human primates, all referred to as M1 herein) provides an ideal cortical region to address questions about cellular evolution in rodents and primates by integrating these approaches. Unlike the primary visual cortex (V1), which is highly specialized in primates, or frontal and temporal association areas, whose homologues in rodents remain poorly defined, M1 is essential for fine motor control and is functionally conserved across placental mammals. M1 is an agranular cortex, lacking a defined L4, although neurons with L4-like properties have been described ^14^. L5 of carnivore and primate M1 contains exceptionally large “giganto-cellular” corticospinal neurons (Betz cells in primates ^15, 16^ that contribute to the pyramidal tract and are highly specialized for their unusually large size with distinctive “taproot”-style dendrites ^17, 18^.

Extracellular recordings from macaque corticospinal neurons reveal distinctive action potential properties supportive of a high conduction velocity and similar, unique properties have been reported during intracellular recordings from giganto-cellular neurons in cats^19–21^. Additionally, some primate Betz cells directly synapse onto alpha motor neurons, whereas in cats and rodents these neurons synapse instead onto spinal interneurons ^22, 23^. These observations suggest that Betz cells possess specialized intrinsic mechanisms to support rapid communication, some of which are primate specific.

Conservation of cellular features across species is strong evidence for evolutionary constraints on important cellular function. To explore evolutionary conservation and divergence of the M1 cellular makeup and its underlying molecular and gene regulatory mechanisms, we combined saturation coverage single-nucleus transcriptome analysis, DNA methylation, and combined open chromatin and transcriptome analysis of mouse, marmoset, and human M1 and transcriptomic profiling of macaque M1 L5. We describe a robust classification of neuronal and non-neuronal cell types in each species that is highly consistent between the transcriptome and epigenome. Cell type alignment accuracy and similarity varied as a function of evolutionary distance, with human more similar to non-human primate than to mouse. We derived a consensus mammalian classification with globally similar cellular diversity, varying proportions, and species specializations in gene expression between conserved cell classes. Few genes had conserved cell type-specific expression across species and likely contribute to other conserved cellular properties, such as the unique morphology of chandelier GABAergic neurons. Conversely, these data also allow a targeted search for genes responsible for species specializations such as the distinctive anatomy, physiology and axonal projections of Betz cells, large corticospinal neurons in primates that are responsible for voluntary fine motor control. Together these findings highlight the strength of a comparative approach to understand cortical cellular diversity and identify conserved and specialized gene and gene regulatory mechanisms underlying cellular identity and function.

We made all primary and analyzed data publicly available. Raw sequence data are available for download from the Neuroscience Multi-omics Archive (nemoarchive.org) and the Brain Cell Data Center (biccn.org/data). Visualization and analysis tools are available at NeMO Analytics (nemoanalytics.org) and Cytosplore Viewer (viewer.cytosplore.org). These tools allow users to compare cross-species datasets and consensus clusters via genome and cell browsers and calculate differential expression within and among species. A semantic representation of the cell types defined through these studies is available in the provisional Cell Ontology (https://bioportal.bioontology.org/ontologies/PCL; Supplementary Table 1).

## Results

### Multi-omic cell type taxonomies

To characterize the molecular diversity of M1 neurons and non-neuronal cells, we applied multiple single-nucleus transcriptomic (plate-based SMART-seq v4, SSv4, and droplet-based Chromium v3, Cv3, RNA-sequencing) and epigenomic (single-nucleus methylcytosine sequencing 2, snmC-seq2; single-nucleus chromatin accessibility and mRNA expression sequencing, SNARE-seq2) assays on isolated M1 samples from human (Extended Data Fig. 1a), marmoset, and mouse brain. Cellular diversity was also profiled selectively in M1 L5 from macaque monkeys using Cv3 (Fig. 1b) to allow Patch-seq mapping in physiology experiments. M1 was identified in each species based on its stereotyped location in the caudal portion of frontal cortex and histological features such as the presence of exceptionally large pyramidal neurons in L5 of M1, classically known as Betz cells in human, other primates, and carnivores (Fig. 1a; ^17^). Single nuclei were dissociated, sorted, and transcripts were quantified with Cv3 for deep sampling in all four species, and additionally using SSv4 in human and mouse for full-length transcript information. For human and a subset of mouse nuclei, individual layers of M1 were profiled independently using SSv4. Whole-genome DNA methylation, and open chromatin combined with transcriptome measurements, were quantified in single nuclei from a subset of species (Fig. 1b). Mouse datasets are also reported in a companion paper ^6^. Median neuronal gene detection was higher in human using SSv4 (7296 genes) than Cv3 (5657), partially due to 20-fold greater read depth, and detection was lower in marmoset (4211) and mouse (5046) using Cv3 (Extended Data Fig. 1b-i).

**Figure 1.**
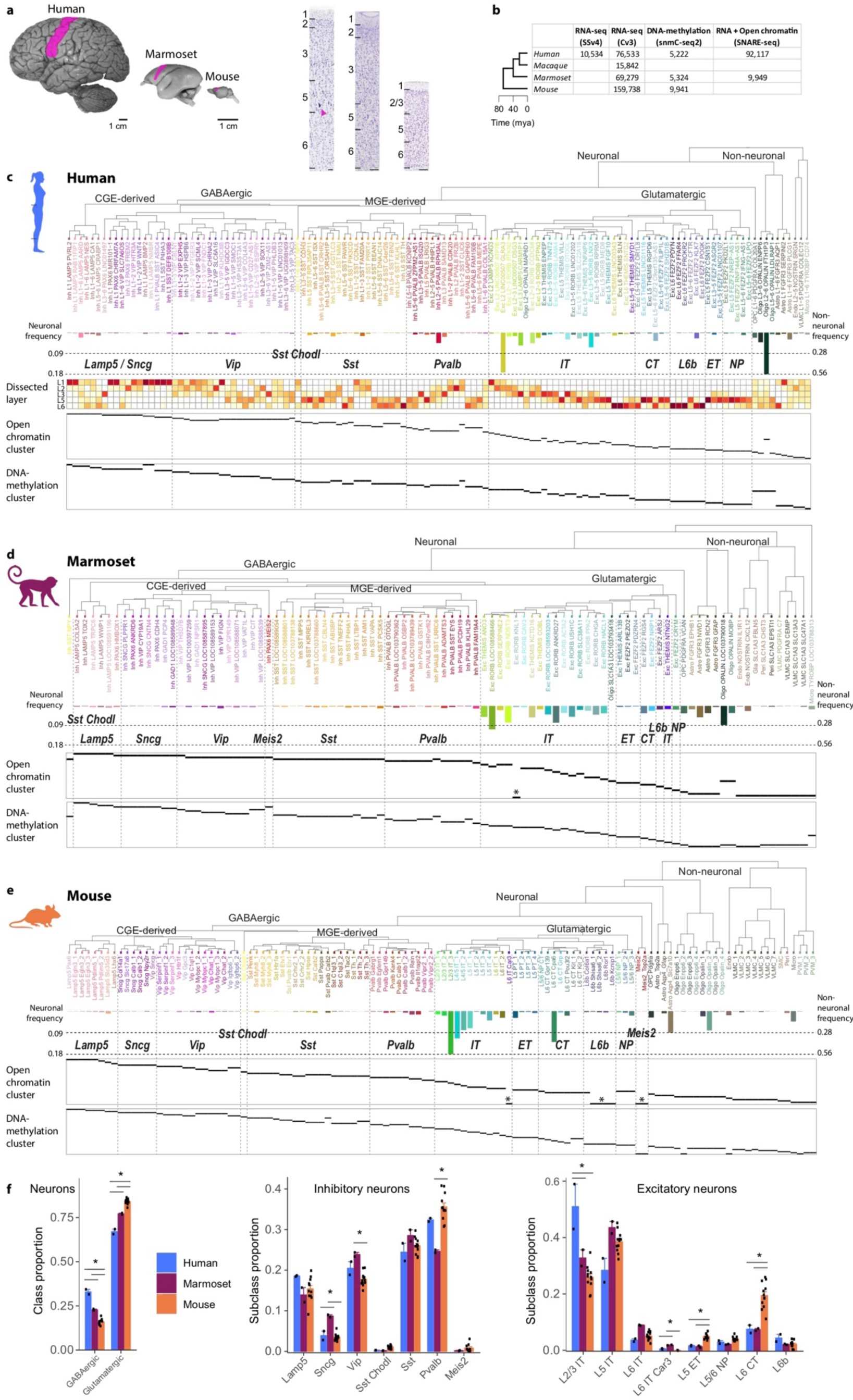
Molecular taxonomy of cell types in M1 of human, marmoset, and mouse. **a**, M1 highlighted in lateral views of neocortex across species. Nissl-stained sections of M1 annotated with layers and showing the relative expansion of cortical thickness, particularly L2 and L3 in primates, and large pyramidal neurons or ‘Betz’ cells in human L5 (arrowhead). Scale bars, 100 µm. b, Phylogeny of species and number of nuclei included in analysis for each molecular assay. All assays used nuclei isolated from the same donors for human and marmoset. SSv4, SMART-Seq v4; Cv3, Chromium v3; mya, millions of years ago. c-e, Dendrograms of cell types defined by RNA-seq (Cv3) for human (c), marmoset (d), and mouse (e) and annotated with cluster frequency and dissected layer (human only). Epigenomic clusters (in rows) aligned to RNA-seq clusters as indicated by horizontal black bars. Asterisks denote RNA clusters that lack corresponding epigenomic clusters. f, Relative proportions of cells in several classes and subclasses were significantly different between species based on an ANOVA followed by Tukey’s HSD tests (asterisk, adjusted P < 0.05).

For each species, a diverse set of neuronal and non-neuronal cell type clusters were defined based on unsupervised clustering of snRNA-seq datasets (cluster metadata in Supplementary Table 2). Human SSv4 and Cv3 data were integrated based on shared co-expression using Seurat ^24^, and 127 clusters were identified that included nuclei from both RNA-seq platforms (Extended Data Fig. j-l). Marmoset clusters (94) were determined based on independent clustering of Cv3 data using a similar analysis pipeline. Mouse clusters (116) were defined in a companion paper ^6^ using seven integrated transcriptomics datasets. These differences in the number of clusters are likely due to a combination of statistical methodological differences as well as sampling and data quality differences rather than true biological differences in cell diversity. For example, more non-neuronal nuclei were sampled in mouse (58,098) and marmoset (21,189) compared to human (4,005), resulting in greater non-neuronal resolution in those species. t-SNE visualizations of transcriptomic similarities across nuclei revealed well-separated clusters in all species and mixing among donors, with some donor-specific technical effects in marmoset (Extended Data Fig. 1m,n).

Post-clustering, cell types were organized into taxonomies based on transcriptomic similarities and given a standardized nomenclature (Supplementary Table 3). As described previously for a different cortical region ^2^, taxonomies were broadly conserved across species and reflected different developmental origins of major non-neuronal and neuronal classes (e.g. GABAergic neurons from ganglionic eminences (GEs) versus glutamatergic neurons from the cortical plate) and subclasses (e.g. GABAergic CGE-derived *Lamp5/Sncg* and *Vip* versus MGE-derived *Pvalb* and *Sst*), allowing identification and naming of these subclasses across species. Consequently, cardinal cell subclass properties can be inferred, such as intratelencephalic (IT) projection patterns. Greater species variation was seen at the highest level of resolution (cell types) that are named based on transcription data in each species including the layer (if available), major class, subclass marker gene, and most specific marker gene (e.g. L3 Exc *RORB OTOGL* in human; additional markers in Supplementary Tables 4-6). GABAergic types were uniformly rare (< 4.5% of neurons), whereas more variable frequencies were found for glutamatergic types (0.01 to 18.4% of neurons) and non-neuronal types (0.15% to 56.2% of non-neuronal cells).

Laminar dissections in human M1 further allowed the estimation of laminar distributions of cell types based on the proportions of nuclei dissected from each layer (Fig. 1c). As expected and previously reported in middle temporal gyrus (MTG) of human neocortex ^2^, glutamatergic neuron types were specific to layers. A subset of CGE-derived *Lamp5/Sncg* GABAergic neurons were restricted to L1, and MGE-derived GABAergic types (*Sst* and *Pvalb*) displayed laminar patterning, with transcriptomically similar types showing proximal laminar distributions, whereas *Vip* GABAergic neuron types displayed the least laminar specificity. Three astrocyte subtypes had frequencies and layer distributions that correlated with known morphologically-defined astrocyte types ^25^, including a common type in all layers (protoplasmic), a rare type in L1 (interlaminar) ^26^, and a rare type in L6 (fibrous).

Single-nucleus sampling provides a relatively unbiased survey of cellular diversity ^2, 27^ and enables comparison of cell subclass proportions across species. For each donor, we estimated the proportion of GABAergic and glutamatergic cells among all neurons and compared the proportions across species. Consistent with previously reported differences in GABAergic neuron frequencies in primate versus rodent somato-motor cortex based on histological measurements (reviewed in ^28^), we found twice as many GABAergic neurons in human (33%) compared to mouse M1 (16%) an intermediate proportion (23%) in marmoset (Fig. 1f). Despite these differences, the relative proportions of GABAergic neuron subclasses were similar. Exceptions to this included an increased proportion of *Vip* and *Sncg* cells and decreased proportion of *Pvalb* cells in marmoset. Among glutamatergic neurons, there were significantly more L2 and L3 IT neurons in human than marmoset and mouse (Fig. 1f), consistent with the dramatic expansion of supragranular cortical layers in human (Fig. 1a) ^29^. The L5 extratelencephalic-projecting (ET) types (also known as pyramidal tract, PT, or subcerebral types), including corticospinal neurons and Betz cells in primate M1, comprised a significantly smaller proportion of glutamatergic neurons in primates than mouse. This species difference was also reported in MTG ^2^, possibly reflecting the spatial dilution of these cells with the expansion of neocortex in primates. Similarly, the L6 cortico-thalamic (CT) neuron populations were less than half as frequent in primates compared to mouse, whereas the L6 *Car3* type was rare in all species and relatively more abundant in marmoset.

Individual nuclei were isolated from M1 of the same donors for each species and molecular profiles were derived for DNA methylation (snmC-seq2) and open chromatin combined with mRNA (SNARE-seq2). Independent unsupervised clustering of epigenomic data also resulted in diverse clusters (see below, Figs. 4 and 5) that were mapped back to RNA clusters based on shared (directly measured or inferred) marker gene expression. Cell epigenomes were highly correlated with transcriptomes, and all epigenomic clusters mapped to one or more transcriptomic clusters. The epigenome data generally had lower cell type resolution (Fig. 1c-e), although this may be due to sampling fewer cells or sparse genomic coverage. Interestingly, snmC-seq2 and SNARE-seq2 resolved different granularities of cell types. For example, more GABAergic *Vip* neuron types were identified in human M1 based on DNA-methylation than open chromatin, despite profiling only 5% as many nuclei with snmC-seq2 (Fig. 1c).

### Consensus cellular M1 taxonomy across species

A consensus cell type classification identifies conserved molecular makeup and allows direct cross-species comparisons. The snRNA-seq Cv3 datasets were integrated using Seurat ^24^ that aligns nuclei across species based on shared co-expression of a subset of orthologous genes with variable expression. We repeated this analysis for three cell classes: GABAergic neurons (Fig. 2), glutamatergic neurons (Extended Data Fig. 3) and non-neuronal cells (Extended Data Fig. 4). As represented in a reduced dimension UMAP space (Fig.2a), GABAergic neuronal nuclei were well-mixed across species. Eight well-defined populations formed distinct islands populated by cells from all three species, including CGE-derived (*Lamp5*, *Sncg*, *Vip*) and MGE-derived (*Pvalb*, *Sst*, *Sst Chodl*) subclasses, and *Lamp5 Lhx6* and chandelier cell (ChC) types. To identify conserved molecular expression for each subclass across species, we first identified genes that were enriched in each subclass (“markers”) compared to all GABAergic neurons in each species (ROC test; AUC > 0.7). Then, we looked for overlap among these genes across species. Each subclass had a core set of conserved markers (Fig. 2b, markers listed in Supplementary Table 7), and many subclass markers were species-specific. The contrast between a minority of conserved and majority of species-specific marker genes enriched in subclasses is particularly clear in the heatmap in Figure 2c (Supplementary Table 8). As expected based on their closer evolutionary distance, human and marmoset shared more subclass markers with each other than with mouse (Fig. 2b).

**Figure 2.**
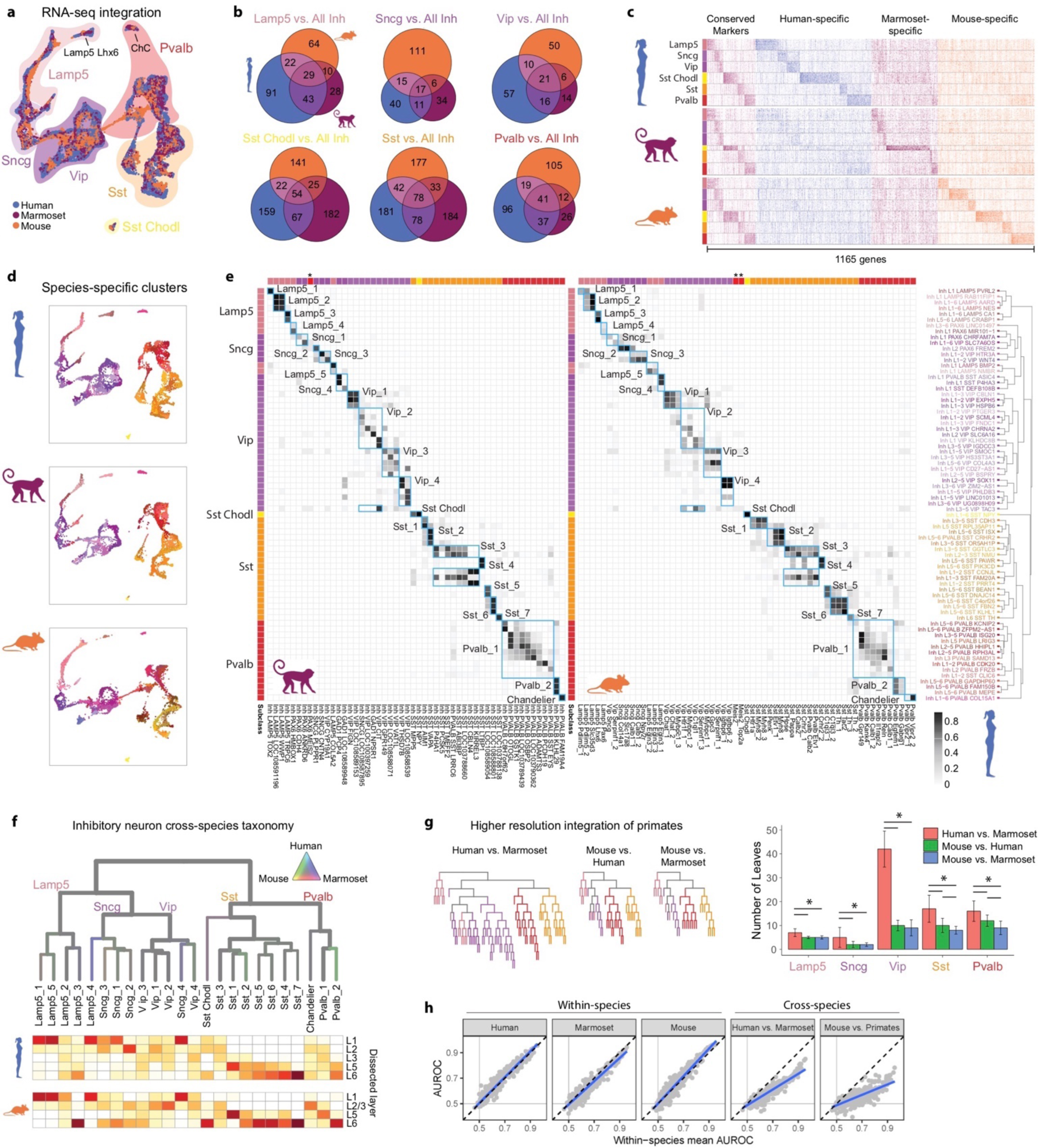
Evolution of GABAergic neuron types across species. **a**, UMAP projection of integrated snRNA-seq data from human, marmoset, and mouse GABAergic neurons. Filled outlines indicate cell subclasses. b, Venn diagrams indicating the number of shared DEGs across species by subclass. DEGs were determined by ROC tests of each subclass versus all other GABAergic subclasses within a species. c, Heatmap of all DEGs from b ordered by subclass and species enrichment. Heatmap shows gene expression scaled by column for up to 50 randomly sampled nuclei from each subclass for each species. d, UMAP projection from a, separated by species, and colored by within-species clusters. e, Cluster overlap heatmap showing the proportion of nuclei in each pair of species clusters that are mixed in the cross-species integrated space. Cross-species consensus clusters are indicated by labeled blue boxes. Human clusters (rows) are ordered by the dendrogram reproduced from Figure 1c. Marmoset (left columns) and mouse (right columns) clusters are ordered to align with human clusters. Color bars at top and left indicate subclasses of within-species clusters. Asterisks indicate marmoset and mouse Meis2 subclasses, which were not present in human. f, Dendrogram of GABAergic neuron consensus clusters with edges colored by species mixture (grey, well mixed). Below: Estimated spatial distributions of clusters based on layer dissections in human (top) and mouse (bottom). g, Dendrograms of pairwise species integrations, colored by subclass. Bar plots quantifying well-mixed leaf nodes. Significant differences (adjusted P < 0.05, Tukey’s HSD test) between species are indicated for each subclass. h, Scatter plots of MetaNeighbor analysis showing the performance (AUROC) of gene-sets to classify GABAergic neurons within and between species. Blue lines, linear regression fits; black lines, mean within species performance; grey lines, performance equivalent to chance.

**Figure 3.**
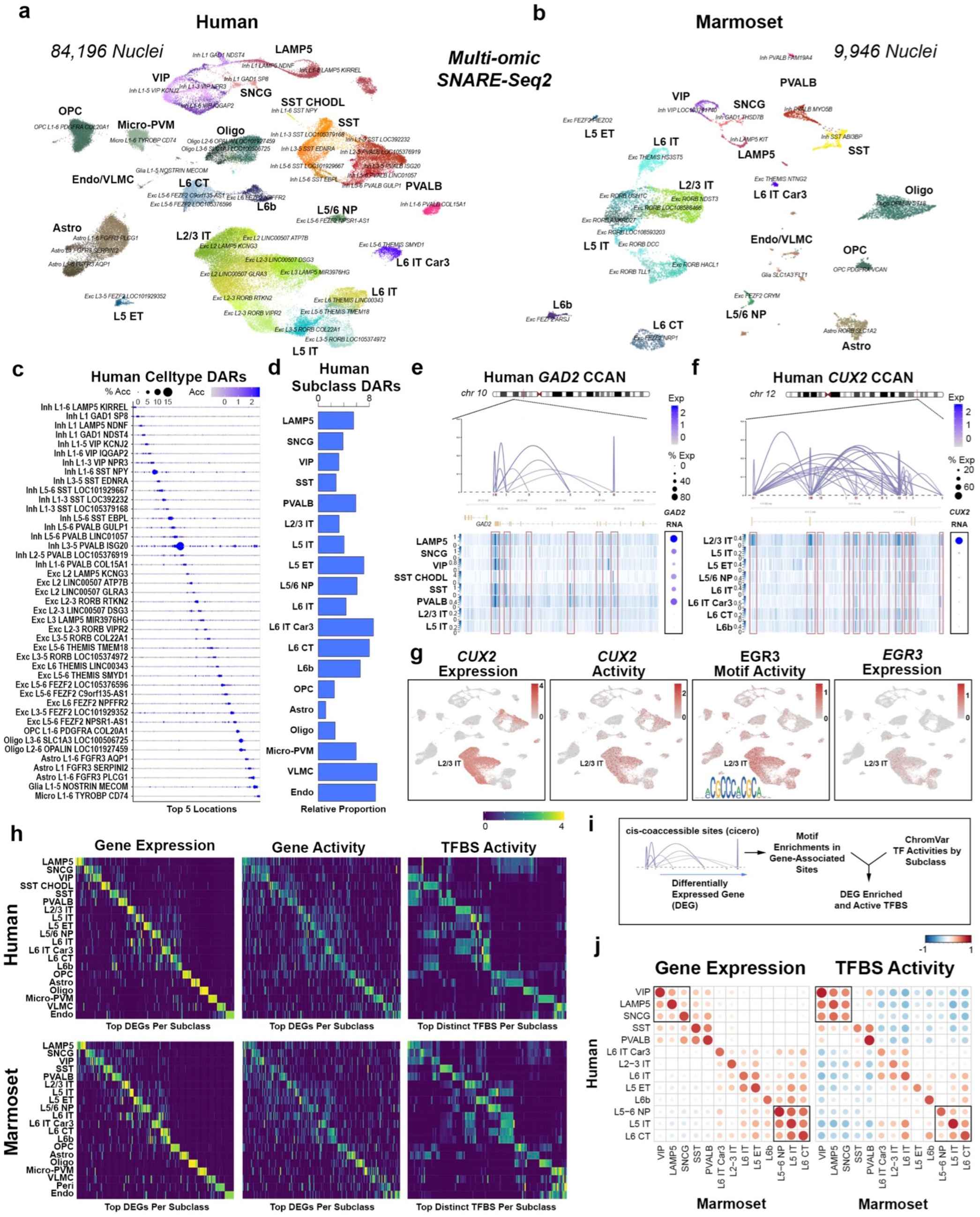
Dual-omic expression and chromatin accessibility reveals regulatory processes defining M1 cell types. **a-b** UMAP visualizations of human (a) and marmoset (b) M1 SNARE-Seq2 data (2 individuals per species) indicating both subclass and accessibility-level cluster identities. c, Dot plot showing proportion and scaled average accessibility of differentially accessible regions (DARs) identified between human AC clusters (adjusted P < 0.001, log-fold change > 1, top 5 distinct sites per cluster). d, Proportion of total human or marmoset DARs identified between subclasses (adjusted P < 0.001, log-fold change > 1) after normalization to cluster sizes. e-f, Connection plots for cis-co-accessible network (CCAN) sites associated with the human *GAD2* (e) and *CUX2* (f) genes. Corresponding AC read pile-up tracts for GABAergic and select glutamatergic subclasses are shown. Right panels are dot plots showing the percentage of expressing nuclei and average gene expression values (log scale) for *GAD2* or *CUX2* within each of the clusters indicated. g, UMAP plots as in Figure 5a (human) showing (scaled from low—gray to high—red) *CUX2* gene expression (RNA) and activity level predicted from AC data. UMAP plots for activity level of the EGR3-binding motif, identified using chromVAR and found to be enriched within *CUX2* co-accessible sites, and the corresponding expression (RNA) of the *EGR3* gene are shown. h, Heatmaps for human (top) and marmoset (bottom) showing TFBS enrichments, according to the scheme outlined in (i), within genes differentially expressed between subclasses and having at least two cis-co-accessible sites. Left panels show scaled average (log scale) gene expression values (RNA) for the top DEGs (adjusted P < 0.05, log-fold change > 1, top 10 distinct sites per cluster visualized), middle panels show the corresponding scaled average cicero gene activity scores and the right panels show scaled values for the corresponding top distinct chromVAR TFBS activities (adjusted P < 0.05, log-fold change > 0.5, top 10 distinct sites per cluster visualized). j, Correlation plots comparing scaled average gene expression profiles (left panel) or chromVAR TFBS activity scores (right panel) between human and marmoset matched subclasses.

**Figure 4.**
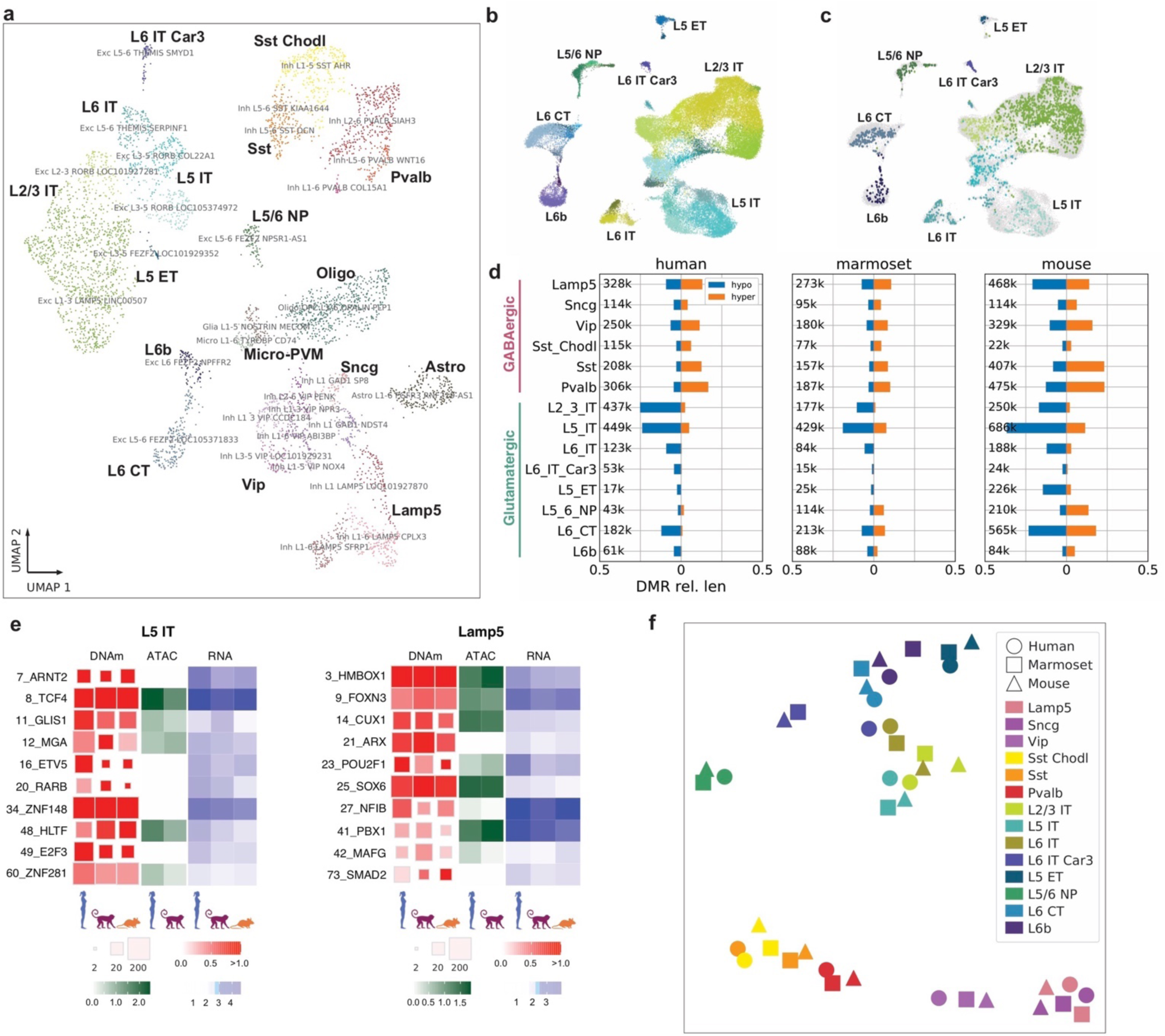
DNA methylation differences across clusters and species. **a**, UMAP visualization of human M1 DNAm-seq (snmC-seq2) data indicating both subclass and DNAm cluster identities. b,c, UMAP visualization of integration between DNAm-seq and RNA-seq of human glutamatergic neurons colored by cell subclass for all nuclei (b) or only nuclei profiled with DNAm-seq (c). d, Barplots of the relative lengths of hypo- and hyper-methylated DMRs among cell subclasses across three species normalized by cytosine coverage genome-wide (Methods). Total number of DMRs for each subclass are listed (k, thousands). e, Distinct TF motif enrichment for L5 IT and *Lamp5* subclasses across species. f, t-SNE visualization of subclass TF motif enrichment that is conserved across species.

**Figure 5.**
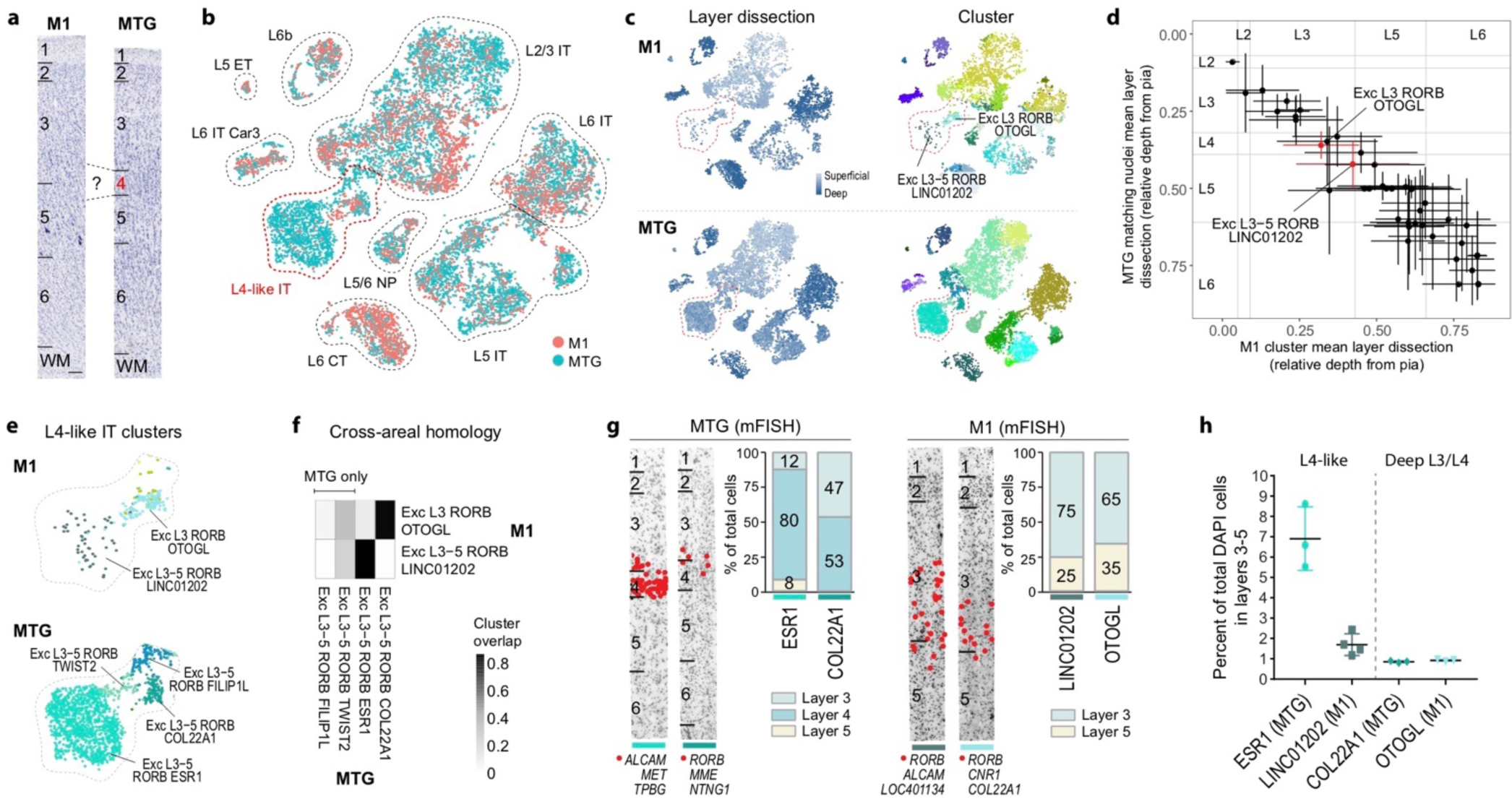
L4-like neurons identified in M1 based on cross-areal cell type homology. **a**, t-SNE projection of glutamatergic neuronal nuclei from M1 and MTG based on similarity of integrated expression levels. Nuclei are intermixed within all cell subclasses. b, Nuclei annotated based on the relative depth of the dissected layer and within-area cluster. A subset of clusters from superficial layers are highlighted. c, Proportion of nuclei in each cluster that overlap between areas. MTG clusters *COL22A1* and *ESR1* map almost one-to-one with M1 clusters *OTOGL* and *LINC01202*, respectively. d, Estimated relative depth from pia of M1 glutamatergic clusters and closest matching MTG neurons based on similarity of integrated expression. Mean (points) and standard deviation (bars) of the dissected layer are shown for each cluster and approximate layer boundaries are indicated for M1 and MTG. e, Magnified view of L4-like clusters in M1 and MTG. Note that MTG clusters *FILIP1L* and *TWIST2* have little overlap with any M1 clusters. f, Overlap of M1 and MTG clusters in integrated space identifies two one-to-one cell type homologies and two MTG-specific clusters. g, ISH labeling of MTG and M1 clusters quantifies differences in layer distributions for homologous types between cortical areas. Cells (red dots) in each cluster were labeled using the markers listed below each representative inverted image of a DAPI-stained cortical column. h, ISH estimated frequencies of homologous clusters shows M1 has a 4-fold sparser L4-like type and similarly rare deep L3 type.

Cell types remained distinct within species and aligned across species in the integrated transcriptomic space (Fig. 2d). To establish a consensus taxonomy of cross-species clusters, we used unsupervised clustering to split the integrated space into more than 500 small clusters (‘metacells’) and built a dendrogram and quantified branch stability by subsampling metacells and reclustering (Extended Data Fig. 2a). Metacells were merged with nearest neighbors until all branches were stable and included nuclei from the three species (see Methods). We used cluster overlap heatmaps to visualize the alignment of cell types across species based on membership in merged metacells (Fig. 2e). 24 GABAergic consensus clusters displayed consistent overlap of clusters among the three species and are highlighted as blue boxes in the heatmaps (Fig. 2e).

We next constructed a consensus taxonomy by pruning the metacell dendrogram (Extended Data Fig. 2a), and demonstrated that all types were well mixed across species (Fig. 2f, grey branches). The robustness of consensus types was bolstered by a conserved set of marker genes (Extended Data Fig. 2d) and high classification accuracy of subclasses (Extended Data Fig. 2e, data in Supplementary Table 9) and types (Extended Data Fig. 2f, data in Supplementary Table 9) compared to nearest neighbors within and among species using a MetaNeighbor analysis ^30^. Distinct consensus types (ChC, *Sst Chodl*) were the most robust (mean AUROC = 0.99 within-species and 0.88 cross-species), while *Sncg* and *Sst* types could not be as reliably differentiated from closely related types (mean AUROC = 0.84 within-species and 0.50 cross-species). Most consensus GABAergic types were enriched in the same layers in human and mouse (Fig. 2f), although there were also notable species differences. For example, ChCs were enriched in L2/3 in mouse and distributed across all layers in human as was seen in temporal cortex (MTG) based on RNA ISH ^2^. *Sst Chodl* was restricted to L6 in mouse and also found in L1 and L2 in human, consistent with previous observations of sparse expression of *SST* in L1 in human not mouse cortex ^31^.

More consensus clusters could be resolved by pairwise alignment of human and marmoset than primates and mouse, particularly *Vip* subtypes (Fig. 2g, Extended Data Fig. 2b). Higher resolution integration of cell types was also apparent in cluster overlap plots between human and marmoset clusters (Fig. 2e, Extended Data Fig. 2c). We quantified the expression conservation of functionally annotated sets of genes by testing the ability of gene sets to discriminate GABAergic consensus types. This analysis was framed as a supervised learning task, both within- and between-species ^30^. Within-species, gene sets related to neuronal connectivity and signaling were most informative for cell type identity (Extended Data Fig. 2g), as reported in human and mouse cortex ^2, 32^. All gene sets had remarkably similar consensus type classification performance across species (r > 0.95; Fig. 2h), pointing to strong evolutionary constraints on the cell type specificity of gene expression central to neuronal function. Gene set classification performance was systematically reduced when training and testing between primates (44% reduction) and between primates and mouse (65% reduction; Fig. 2h). Therefore, many cell type marker genes were expressed in different consensus types between species. Future comparative work can compare reductions in classification performance to evolutionary distances between species to estimate rates of expression change across phylogenies.

Cross-species consensus types were defined for glutamatergic neurons using an identical approach as for GABAergic neurons (Extended Data Fig. 3). In general, glutamatergic subclasses aligned well across species and had a core set of conserved markers as well as many species-specific markers (Extended Data Fig. 3a-c, genes listed in Supplementary Tables 10-11). 13 consensus types were defined across species. Glutamatergic types had fewer conserved markers than GABAergic types (Extended Data Fig. 3d-f,j), although subclasses and types were similarly robust (mean within-species AUROC = 0.86 for GABAergic types and 0.85 for glutamatergic types) based on classification performance (Extended Data Fig. 3k,l and Supplementary Table 9). Human and marmoset had consistently more conserved marker genes than primates and mouse (Extended Data Fig. 3i) and could be aligned at somewhat higher resolution (Extended Data Fig. 3g,h) for L5/6 NP and L5 IT subclasses.

Integration of non-neuronal cells was performed similarly to neurons (Extended Data Fig. 4a). Consensus clusters (blue boxes in Extended Data Fig. 4c) that shared many marker genes were identified across species (Extended Data Fig. 4d), and there was also evidence for the evolutionary divergence of gene expression in consensus types. For example, the Astro_1 type had 560 DEGs (Wilcox test; FDR < 0.01, log-fold change > 2) between human and mouse and only 221 DEGs between human and marmoset (Extended Data Fig. 4e). The human cortex contains several morphologically distinct astrocyte types ^33^: interlaminar (ILA) in L1, protoplasmic in all layers, varicose projection in deep layers, and fibrous in white matter (WM). We previously reported two transcriptomic clusters in human MTG that corresponded to protoplasmic astrocytes and ILAs ^2^, and we validated these types in M1 (Extended Data Fig. 4g,h). We identified a third type, Astro L1-6 *FGFR3 AQP1*, that expresses *APQ4* and *TNC* and corresponds to fibrous astrocytes in WM (Extended Data Fig. 4g, left ISH). A putative varicose projection astrocyte did not express human astrocyte markers (Extended Data Fig. 4g, middle and right ISH), and this rare type may not have been sampled or is not transcriptomically distinct.

Species comparison of non-neuronal cell types was more challenging than for neurons due to variable sampling across species and more immature non-neuronal cells in mouse. 5-to 15-fold lower sampling of non-neuronal cells in human impacted detection of rare types. For example, pericytes, smooth muscle cells (SMCs), and some subtypes of vascular and leptomeningeal cells (VLMCs) were present in marmoset and mouse and not human datasets (Extended Data Fig. 4c, right plot, blue arrows), although these cells are clearly present in human cortex (for example, see ^34^). A maturation lineage between oligodendrocyte precursor cells (OPCs) and oligodendrocytes based on reported marker genes ^35^ that was present in mouse and not primates (Extended Data Fig. 4b) likely represents the younger age of mouse tissues used. Mitotic astrocytes (Astro_*Top2a*) were also only present in mouse (Extended Data Fig. 4a,c) and represented 0.1% of non-neuronal cells. Primates had a unique oligodendrocyte population (Oligo *SLC1A3 LOC103793418* in marmoset and Oligo L2-6 *OPALIN MAP6D1* in human) that was not a distinct cluster in mouse (Extended Data Fig. 4c, left plot, blue arrow). Surprisingly this oligodendrocyte clustered with glutamatergic types (Fig. 1c,d) and was associated with neuronal transcripts such as *NPTX1*, *OLFM3*, and *GRIA1* (Extended Data Fig. 4i). This was not an artifact, as FISH for markers of this type (*SOX10*, *ST18*) co-localized with neuronal markers in the nuclei of cells that were sparsely distributed across all layers of human and marmoset M1 (Extended Data Fig. 4j). This may represent an oligodendrocyte type that expresses neuronal genes or could represent phagocytosis of parts of neurons and accompanying transcripts that are sequestered in phagolysosomes adjacent to nuclei.

To assess differential isoform usage between human and mouse, we used SSv4 data with full transcript coverage and estimated isoform abundance in cell subclasses. Remarkably, 25% of moderately expressed (> 10 transcripts per million) isoforms showed a large change (>9-fold) in usage between species, and isoform switching was 30-60% more common in non-neuronal than neuronal cells (Extended Data Fig. 2h,i, Supplementary Table 12). For example, β2-Chimaerin (*CHN2*), a gene shown to mediate axonal pruning in the hippocampus ^36^, was highly expressed in human and mouse L5/6 NP neurons. In mouse, the short isoform was almost exclusively expressed, while in human, longer isoforms were also expressed (Extended Data Fig. 2j).

### Open chromatin profiling reveals distinct cell type gene regulation

To directly match accessible chromatin profiles to RNA-defined cell populations, we used SNARE-Seq ^37^, now modified for highly multiplexed combinatorial barcoding (SNARE-Seq2) ^38^. We generated 84,178 and 9,946 dual-omic single-nucleus RNA and accessible chromatin (AC) datasets from human (n = 2) and marmoset (n = 2) M1, respectively (Extended Data Fig. 5a-b, Supplementary Table 13). On average, 2,242 genes (5,764 unique transcripts) were detected per nucleus for human and 3,858 genes (12,400 unique transcripts) per nucleus for marmoset, due to more than 4-fold greater sequencing depth for marmoset (average 17,576 reads per nucleus for human and 77,816 reads per nucleus for marmoset).

To define consensus clusters, SNARE-seq2 single-nucleus RNA expression data were mapped to human and marmoset transcriptomic clusters (Fig. 1c,d) based on correlated expression of cell type marker genes. SNARE-seq2 transcriptomes were also independently clustered, with both approaches giving consistent results (Extended Data Fig. 5c-f). Consensus clusters were more highly resolved in transcriptomic compared to AC data (Extended Data Fig. 5g), and so an integrative approach was used to achieve best matched AC-level cluster annotations (Extended Data Fig. 5h-k). AC peak calling at multiple levels of cellular identity (for RNA consensus clusters, resolved AC clusters, subclasses and classes) yielded a combined total of 273,103 (human) and 134,769 (marmoset) accessible sites, with an average of 1527 or 1322 unique accessible peak fragment counts per nucleus, respectively. Gene activity estimates based on cis-regulatory interactions predicted from co-accessible promoter and distal peak regions using Cicero ^39^ were highly correlated with RNA expression values. This highlights the ability of SNARE-Seq2 to meaningfully characterize AC at RNA-defined cellular resolution that cannot be achieved using only AC data (Extended Data Fig. 6a-b). The AC-level clusters (Fig. 3a,b) that showed similar coverage across individual samples (Extended Data Fig. 6c-f) revealed regions of open chromatin that are extremely cell type specific (Fig. 3c). These regulatory regions were relatively more abundant in glutamatergic compared to GABAergic neuron subpopulations (Fig. 3c-d, Supplementary Table 14).

**Figure 6.**
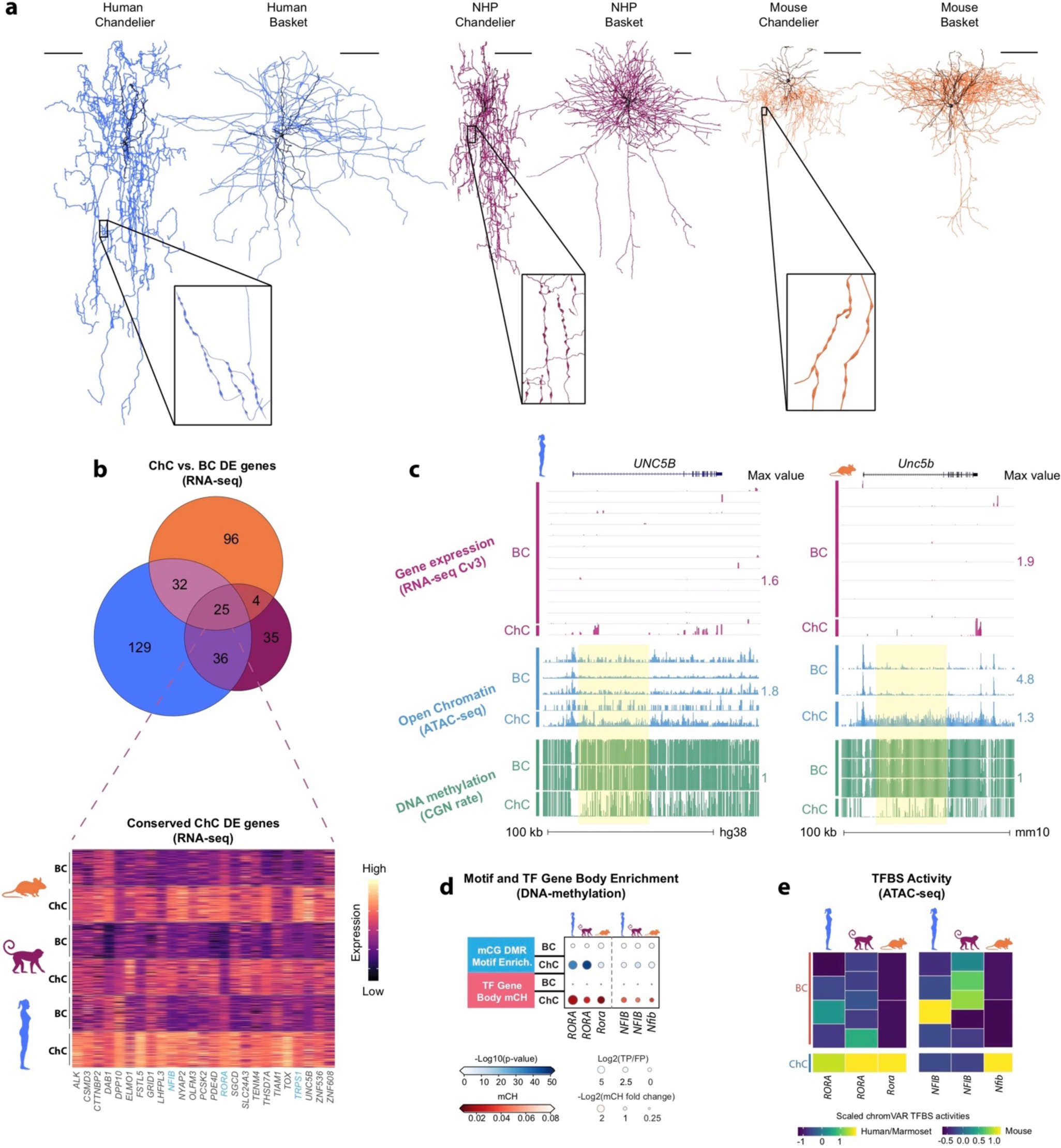
Chandelier neurons have a core set of conserved molecular features. **a**, Representative ultrastructure reconstructions of a ChC and BC from human (left), macaque (middle), and mouse (right). Insets show higher magnification of ChC axon cartridges. Macaque reconstructions were from source data available in Neuromorpho ^63, 64^. Mouse cells also appear in ^65^. b, Venn diagram indicating the number of shared ChC-enriched genes across species (top). DEGs were determined by a ROC test of ChCs against BCs within a species. Heatmap showing scaled expression of the 25 conserved DEGs in 100 randomly selected ChC and BC nuclei for each species (bottom); transcription factors are colored in blue. c, Genome browser tracks showing *UNC5B* locus in human (left) and mouse (right) ChCs and BCs. Tracks show aligned transcripts, regions of accessible chromatin, CGN methylation rate, and CHN methylation rate. Yellow highlights mark examples of ChC-enriched regions of accessible chromatin with hypo-methylated CGN. d, Heatmaps of TF gene body hypo-methylation (mCH) state (bottom half, red) and genome-wide enrichment of TF motif across mCG DMRs in ChCs and BCs (top half, blue). e, Scaled TFBS activities identified from SNARE-seq2 for human and marmoset according to the scheme in Figure 4i and from mouse snATAC-seq data, using genes enriched in ChC versus BC (Supplementary Table 19). Rows correspond to BC and ChC clusters identified in snATAC-seq and SNARE-seq2 datasets.

To better understand the interplay of gene regulation and expression, we compared transcript counts and open chromatin measured in the same nuclei. For example, the GABAergic neuron marker *GAD2* and the L2/3 glutamatergic neuron marker *CUX2* showed cell-type specific chromatin profiles for co-accessible sites that were consistent with their corresponding transcript abundances (Fig. 3e-g).

Transcription factor binding site (TFBS) activities were calculated using chromVAR ^40^, permitting discovery of differentially active TFBSs between cell types. To investigate the regulatory factors that may contribute to marker gene expression, we evaluated active TFBSs for their enrichment within marker gene co-accessible sites. This permitted direct cell type mapping of gene expression and activity levels with the expression and activity of associated regulatory factors (Fig. 3g). Using this strategy, we identified TFBS activities associated with subclass (Fig. 3h-i) and AC-cluster level differentially expressed genes (DEGs) in human and marmoset (Supplementary Table 15). DEG transcript levels and AC-inferred gene activity scores showed high correspondence (Fig. 3h). While most subclasses also showed distinct TFBS activities, correspondence between human and marmoset was higher for glutamatergic rather than GABAergic neurons (Fig. 3h,j). For GABAergic neuron subclasses, gene expression profiles were more conserved than TFBS activities, consistent with fewer differences between GABAergic subpopulations based on AC sites (Fig. 3a,b). This observation is also consistent with fewer distinct TFBS activities among some inhibitory neuron subclasses (*Lamp5*, *Sncg*) in human compared to marmoset (Fig. 3h), despite these cell types having a similar number of AC peak counts (Extended Data Fig. 6d-f). Interestingly, glutamatergic neurons in L5 and L6 showed higher correspondence between primates based on TFBS activities compared to average expression, suggesting that gene regulatory processes are more highly conserved in these subclasses than target gene expression.

### Methylomic profiling reveals conserved gene regulation

We used snmC-seq2 ^41^ to profile the DNA methylome from individual cells in M1. Single-nuclei were labeled with an anti-NeuN antibody and isolated by fluorescence-activated cell sorting (FACS), and neurons were enriched (90% NeuN+ nuclei) to increase detection of rare types. Using snmC-seq2, we generated single-nucleus methylcytosine datasets from M1 of human (n = 2 donors, 6,095 nuclei), marmoset (n = 2, 6,090), and mouse (9,876) (Liu et al. companion paper) (Supplementary Table 16). On average, 5.5 ± 2.7% (mean ± s.d.) of human, 5.6 ± 2.9% of marmoset and 6.2 ± 2.6% of mouse genomes were covered by stringently filtered reads per cell, with 3.4 × 10^4^ (56%), 1.8 × 10^4^ (62%) and 4.5 × 10^4^ (81%) genes detected per cell in the three species, respectively. Based on the DNA methylome profiles in both CpG sites (CG methylation or mCG) and non-CpG sites (CH methylation or mCH), we clustered nuclei (Methods) to group cell populations into 31 cell types in human, 36 cell types in marmoset, and 42 cell types in mouse (Fig. 4a and Extended Data Fig. 7a,b). For each species, cell type clusters could be robustly discriminated using a supervised classification model and had distinct marker genes based on DNA methylation signatures for neurons (mCH) or non-neuronal cells (mCG) (Methods). Differentially methylated regions (DMR) were determined for each cell type versus all other cell types and yielded 9.8 × 10^5^ DMRs in human, 1.0 × 10^6^ in marmoset, and 1.8 × 10^6^ in mouse.

**Figure 7.**
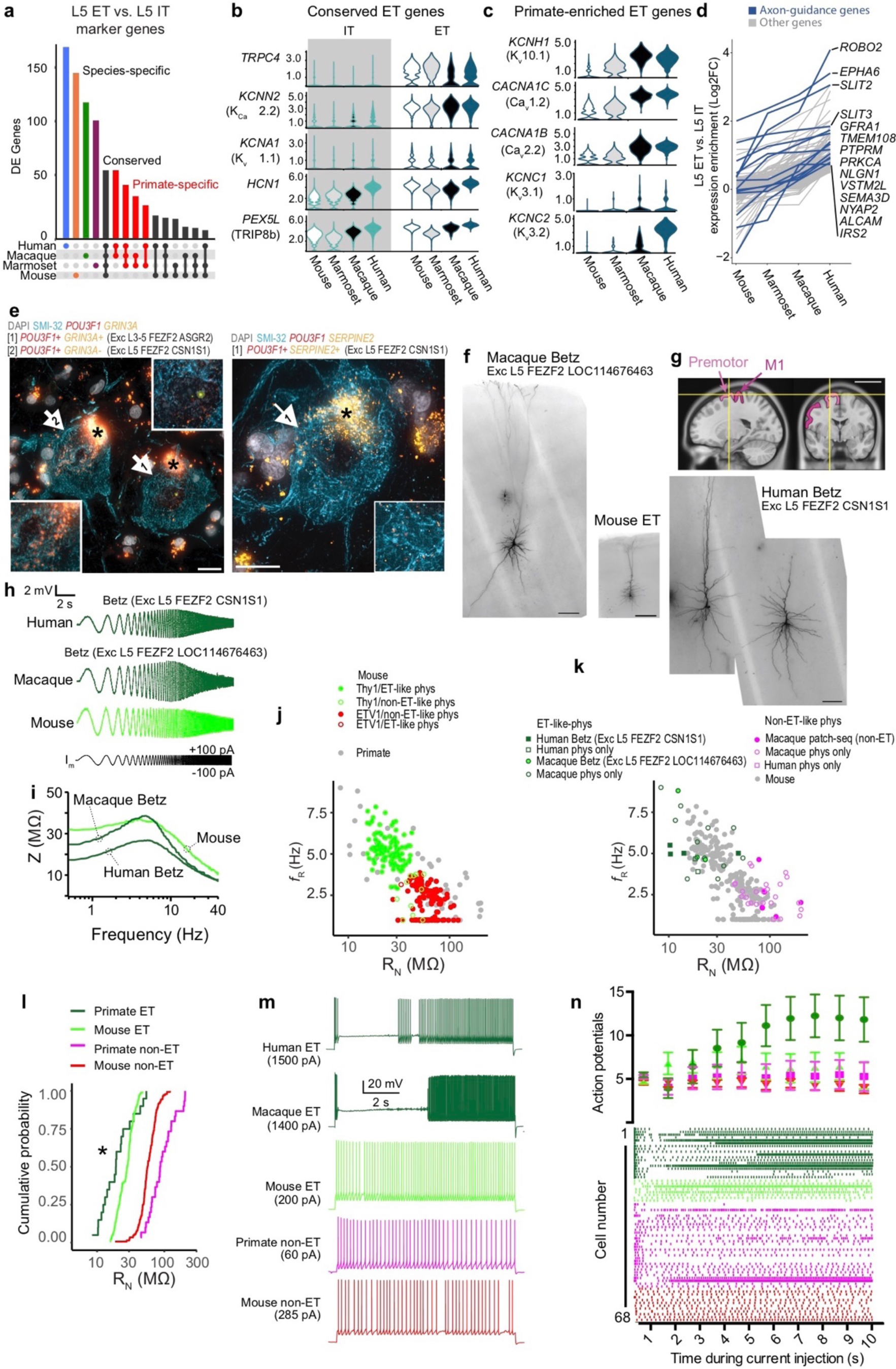
Betz cells have specialized molecular and physiological properties. **a**, Upset plot showing conserved and divergent L5 ET glutamatergic neuron marker genes. DEGs were determined by performing a ROC test between L5 ET and L5 IT within each species. **b, c**, Violin plots of ion channel-related gene expression for genes that are enriched in (**b**) ET versus IT neurons and in (**c**) primate versus mouse ET neurons. Protein names are in parentheses. **d**, Line graph of 131 genes with expression enrichment in L5 ET versus IT neurons in human (>0.5 log_2_ fold-change) that decreases with evolutionary distance from human. **e**, Two example photomicrographs of ISH labeled, SMI-32 IF stained Betz cells in L5 human M1. Cells corresponding to two L5 ET clusters are labeled based on two sets of marker genes. Insets show higher magnification of ISH in corresponding cells. Asterisks mark lipofuscin; scale bar, 20 µm. **f, g**, Exemplar biocytin fills obtained from patch-seq experiments in human, macaque and mouse brain slices. The example human and macaque neurons mapped to a Betz cell transcriptomic cell type. Scale bars, 200 µm. **g**, MRI image in sagittal and coronal planes and approximate location of excised premotor cortex tissue (yellow lines) and adjacent M1. **h**, Voltage responses to a chirp stimulus for the neurons shown in **f** and **g** (left human neuron). **i**, Corresponding ZAP profiles. All neurons were clustered into putative ET and non-IT neurons based upon their resonant frequency and input resistance. **j**, For mouse L5 neurons (*Thy1*-YFP line H, n=117; *Etv1*-EGFP line, n=123; unlabeled, n=21) 99.2 % of neurons in the *Etv1*-EGFP line possessed non-ET-like physiology, whereas, 91.4% of neurons in the *Thy1*-YFP line H had ET-like physiology. **k**, For primate L5 neurons (human, n=8, macaque n=42), all transcriptomically-defined Betz cells (human, n=4, macaque n=3) had ET-like physiology (human n=6, macaque, n=14), whereas all transcriptomically-defined non-ET neurons (human n=2, macaque n=3) had non-ET like physiology (human n=2, macaque n=28). **l**, Cumulative probability distribution of L5 neuron input resistance for primate versus mouse. * p = 0.0064, Kolmogorov-Smirnov test between mouse and primate ET neurons. **m**, Example voltage responses to 10s step current injections for monkey, mouse and human ET and non-ET neurons. The amplitude of the current injection was adjusted to produce ∼5 spikes during the first second. **n**, Raster plot (below) and average firing rate (above) during 1 s epochs during the 10s DC current injection. Primate ET neurons (pooled data from human and macaque) displayed a distinctive decrease followed by a pronounced increase in firing rate over the course of the current injection. Notably, a similar biphasic-firing pattern is observed in macaque corticospinal neurons *in vivo* during prolonged motor movements ^66^, suggesting that the firing pattern of these neurons during behavior is intimately tied to their intrinsic membrane properties.

We determined a consensus molecular classification of cell types in each species by integrating single-nucleus methylomic data with the Cv3 transcriptomic data described above using measurements of gene body differential methylation (CH-DMG) to approximate expression levels. Nuclei from the two data modalities mixed well as visualized in ensemble UMAPs (Fig. 4b,c). Methylation clusters have one-to-one, one-to-many, or many-to-many mapping relation to transcriptomic clusters (Fig. 1c-e and Extended Data Fig. 7d-f). DMRs were quantified for each subclass versus all other subclasses (Fig. 4d), and glutamatergic neurons had more hypo-methylated DMRs compared to GABAergic neurons. Methylome tracks at subclass level can be found at http://neomorph.salk.edu/aj2/pages/cross-species-M1/. To identify enriched transcription factor binding sites (TFBS) in each species and subclass, we performed motif enrichment analysis with hypo-methylated DMRs from one subclass against other DMRs of the same species, and identified 102 ± 57 (mean ± s.d.) TFBS in each subclass (Extended Data Fig. 8 and Supplementary Table 17). We repeated the enrichment analysis using TFBS motif clusters ^42^ and found similarly distinct subclass signatures (Supplementary Table 18). Although subclasses had unique marker genes (Fig. 2c, genes listed in Supplementary Table 8) and CH-DMG across species, they had remarkably conserved TFBS motif enrichment (Fig. 4e,f and Extended Data Fig. 8). For example, *TCF4* is robustly expressed in L5 IT neurons across species and shows significant TFBS enrichment in hypo-methylated DMRs and AC sites. DMRs and AC sites provide independent epigenomic information (Extended Data Fig. 7f,g) and can identify different TFBS enrichment, such as for *ZNF148* in L5 IT neurons. These results are consistent with previous observations of conserved TF network architectures in neural cell types between human and mouse (Stergachis et al. 2014). Conserved sets of TFs have the potential to determine conserved and divergent expression in consensus types based on shared or altered genomic locations of TFBS motifs across species.

### Layer 4-like neurons in human M1

M1 lacks a L4 defined by a thin band of densely packed “granular” neurons that is present in other cortical areas, such as MTG (Fig. 5a), although prior studies have identified neurons with L4-like synaptic properties in mice ^14^ and expression of *RORB*, a L4 marker, in non-human primate M1 ^43^. To address the potential existence of L4-like neurons in human M1 from a transcriptomic perspective, we integrated snRNA-seq data from M1 and the granular MTG, where we previously described multiple L4 glutamatergic neuron types ^2^. This alignment revealed a broadly conserved cellular architecture between M1 and MTG (Fig. 5b,c, Extended Data Fig. 9) including M1 neuron types Exc L3 *RORB OTOGL* (here, *OTOGL*) and Exc L3-5 *RORB LINC01202* (here, *LINC01202*) that map closely to MTG neurons in deep L3 to L4 (Fig. 5c, red outlines). Interestingly, four MTG L2/3 IT types (*LTK*, *GLP2R*, *FREM3*, and *CARM1P1*) whose distinct physiology and morphology are reported in a companion paper ^44^ had less clear homology in M1 than other types (Extended Data Fig. 9a-c), pointing to more variability across cortical areas of superficial as compared to deep glutamatergic neurons. To compare laminar positioning in M1 and MTG, the relative cortical depth from pia for each neuron was estimated based on the layer dissection and average layer thickness ^45^. Transcriptomically similar cell types were found at similar cortical depths in M1 and MTG, and the *OTOGL* and *LINC01202* types were located in deep L3 and superficial L5 in M1 (Fig. 5d).

MTG contains three main transcriptomically-defined L4 glutamatergic neuron types (*FILIP1L*, *TWIST2* and *ESR1*) and a deep L3 type (*COL22A1*) that is found on the border of L3 and L4 (Fig. 5e-g). The M1 types *OTOGL* and *LINC01202* matched one-to-one with MTG *COL22A1* and *ESR1*, whereas there were no matches for the other two MTG L4 types (Fig. 5f). Based on snRNA-seq proportions, the L4-like *OTOGL* type was much sparser in M1 than the *ESR1* type in MTG (Fig. 5e). Multiplex fluorescent in situ hybridization (mFISH) with probes to cell type marker genes confirmed these findings. The MTG *ESR1* type was highly enriched in L4, ^2^, and the homologous M1 *LINC01202* type was sparser and more widely distributed across L3 and L5 (Fig. 5g). The MTG *COL22A1* type was tightly restricted to the L3/4 border ^2^, and the M1 *OTOGL* type was similarly found at the L3/5 border. Quantification of labeled cells as a fraction of DAPI+ cells in L3-5 showed similar frequencies of M1 *OTOGL* and MTG *COL22A1* types and 4-fold sparser M1 *LINC01202* type versus MTG *ESR1* type (Fig. 5h). These data indicate a conservation of deep L3 glutamatergic types and proportions across human cortical areas, but with reduced diversity and sparsification of L4-like neurons to a single (ESR1) type in M1, distributed more broadly where L4 would be if tightly aggregated.

### Chandelier cells share a core molecular identity across species

Conserved transcriptomic and epigenomic features of consensus types likely contribute to cell function and generate hypotheses about the gene regulatory mechanisms underlying cell type identity. Focused analysis of *Pvalb*-expressing GABAergic neurons illustrates the power of these data to predict such gene-function relationships. Cortical *Pvalb*-expressing neurons comprise two major types — basket cells (BCs) and ChCs — that have fast-spiking electrical properties and distinctive cellular morphologies. BCs selectively synapse onto the perisomatic region of glutamatergic pyramidal neurons. ChCs, also called axo-axonic cells ^46^, selectively innervate the axon initial segment (AIS) of pyramidal cells and have unique synaptic specializations called axon cartridges. These cartridges run perpendicular to their post-synaptic target axon, giving a characteristic morphological appearance of candlesticks on a chandelier. This highly conserved feature is shown with biocytin-filled cells from mouse, rhesus macaque, and human (Fig. 6a). To reveal evolutionarily conserved transcriptomic hallmarks of ChCs, we identified DEGs in ChCs versus BCs in each species using an ROC test. 357 DEGs were identified in at least one species, and marmoset ChCs shared more DEGs with human (61 genes) than mouse (29; Fig. 6b, Supplementary Table 19). Remarkably, only 25 DEGs were conserved across all three species. One conserved gene, *UNC5B* (Fig. 6c), is a netrin receptor involved in axon guidance and may help target ChC to pyramidal neuron AIS. Three transcription factors (*RORA*, *TRPS1*, and *NFIB*) were conserved markers and may contribute to gene regulatory networks that determine the unique attributes of ChCs.

To determine if ChCs had enriched epigenomic signatures for *RORA* and *NFIB* (*TRPS1* lacked motif data), we compared DMRs between ChCs and BCs. In all species, *RORA* and *NFIB* had significant CH-DMGs in ChCs not BCs (Fig. 6d), consistent with differential expression. To discern if these TFs may preferentially bind to DNA in ChCs, we tested for TF motif enrichment in hypo-methylated (mCG) DMRs and AC sites genome-wide. We found that the RORA motif was significantly enriched in DMRs in primates (Fig. 6d) and in AC sites of ChCs in all species (Fig. 6e, Supplementary Table 14). The NFIB binding motif was only significantly enriched in AC sites of mouse ChCs, possibly because enrichment was transient during development or NFIB specificity is due to expression alone. Three independent genomic assays converge to implicate *RORA* as a ChC-specific TF among *Pvalb*-expression neurons. Notably, 60 of 357 DEGs contained a ROR-motif in DMRs and AC regions in at least one species, further implicating *RORA* in defining ChC identity.

### Primate Betz cell specialization

In mouse cortex, L5 glutamatergic neurons have distinct long-range projection targets (ET versus IT) and transcriptomes ^1^. L5 ET and IT neuron subclasses clearly align between human and mouse using snRNA-seq in M1 (Extended Data Fig. 3) and in temporal ^2^ and fronto-insular cortex ^12^. Betz cells in L5 of primate M1 connect to spinal motor-neurons via the pyramidal tracts and are predicted to be L5 ET neurons. The species aligned transcriptomic types allow for the identification of genes whose expression may contribute to conserved ET versus IT features and primate-specific physiology, anatomy, and connectivity. Furthermore, Patch-seq methods that jointly measure the transcriptome, physiological properties and morphology of cells, allow the direct identification and characterization of L5 ET and IT neurons across mouse, non-human primate, and human. As primate physiology experiments are largely restricted to macaque, we also profiled L5 of macaque M1 with snRNA-seq (Cv3) to allow accurate Patch-seq mapping.

L5 ET neurons had many DEGs compared to L5 IT neurons in all 4 species. Approximately 50 DEGs were conserved across all species and similarity to human varied as a function of evolutionary distance (Fig. 7a, Supplementary Table 20). Several genes encoding ion channel subunits were enriched in ET versus IT neurons in all species, potentially mediating conserved ET physiological properties (Fig. 7b). A number of additional potassium and calcium channels were primate-enriched (Fig. 7c), potentially underlying primate-specific ET or Betz cell physiology. Interestingly, many of these primate-specific ET-enriched genes showed gradually increasing ET specificity in species more closely related to human.

To explore this idea of gradual evolutionary change further, we identified genes with increasing L5 ET versus IT specificity as a function of evolutionary distance from human (Fig. 7d, Supplementary Table 21). Interestingly, this gene set was highly enriched for genes associated with axon guidance including members of the Robo, Slit and Ephrin gene families. These genes are potential candidates for regulating the cortico-motoneuronal connections associated with increasingly dexterous fine motor control across these species ^23^.

To investigate if transcriptomically defined L5 ET types contain anatomically-defined Betz cells, FISH for L5 ET neurons was combined with immunolabeling against SMI-32, a protein enriched in Betz cells and other long-range projecting neurons in macaque ^47–49^ (Fig. 7e). Cells consistent with the size and shape of Betz cells were identified in two L5 ET clusters (Exc L5 *FEZF2 ASGR2* and Exc L5 *FEZF2 CSN1S1*). Similar to previous reports on von Economo neurons in the insular cortex ^12^, ET clusters in M1 also included neurons with non-Betz morphologies.

To facilitate cross-species comparisons of Betz cells and mouse ET neurons we made patch clamp recordings from L5 neurons in acute and cultured slice preparations of mouse and macaque M1. For a subset of recordings, Patch-seq analysis was applied for transcriptomic cell type identification (Extended Data Fig. 10h). To permit visualization of cells in heavily myelinated macaque M1, we used AAV viruses to drive fluorophore expression in glutamatergic neurons in macaque slice culture (Extended Data Fig. 10g). As shown in Figure 7f, Patch-seq neurons mapping to the macaque Betz/ET cluster (Exc L5 *FEZF2 LOC114676463*) had large somata (diameter > 65 μm) and long “tap root” basal dendrites, canonical hallmarks of Betz cell morphology ^17, 50^. A unique opportunity to record from neurosurgical tissue excised from human premotor cortex (near the confluence of the precentral and superior frontal gyri) during an epilepsy treatment surgery using the same methods as for macaque yielded multiple neurons that mapped transcriptomically to one of the Betz-containing cell types and had canonical Betz cell morphology (Fig. 7g). Macaque and human ET neurons were grouped for physiological analysis because intrinsic properties were not significantly different, and many corticospinal axons originate from premotor cortex^23^.

Shared transcriptomic profiles of mouse, primate, and human L5 ET neurons predicted conservation of some physiological properties of rodent and primate neurons. Transcriptomically-defined ET neurons across species expressed high levels of genes encoding an HCN channel-subunit and a regulatory protein (*HCN1* and *PEX5L;* Fig. 7b). We hypothesized that HCN-dependent membrane properties, which are used to distinguish rodent ET from IT neurons ^51^, would similarly separate cell types in primates. Some primate L5 neurons possessed distinctive HCN-related properties such as a lower input resistance (R_N_) and a peak resonance (*f*_R_) in voltage response around 3-9 Hz (Fig. 7h,i), similar to rodent ET neurons. To determine whether HCN-related physiology is a conserved feature of L5 neurons, we grouped all neurons into physiologically defined ET and non-ET neurons based on their R_N_ and *f*_R_. We asked whether these physiologically-defined neurons corresponded to genetically-defined ET/Betz or non-ET neurons using Patch-seq and cell-type specific mouse lines. For mouse M1, the ET-specific *Thy1*-YFP^21, 52^ and IT specific *Etv1*-EGFP^53^ mouse lines preferentially labeled physiologically defined ET and non-ET neurons, respectively (Fig. 7j). For primates, transcriptomically-defined Betz cells were physiologically defined ET neurons, whereas transcriptomically defined non-ET neurons were physiologically defined non-ET neurons (Fig. 7k). Thus, there was broad correspondence between physiologically-defined and genetically-defined ET neurons in both mouse and primate M1.

There were notable differences in physiology between mouse and primate ET neurons, however. A greater fraction of primate ET neurons exhibited an exceptionally low R_N_ compared to mouse (Fig. 7l). Additional differences in action potential properties across cell types and species may be explained in part by differences in the expression of ion channel-related genes (Fig. 7c, Extended Data Fig. 10).

Most strikingly, primate Betz/ET neurons displayed a distinctive biphasic-firing pattern during long spike trains. The firing rate of both primate and mouse non-ET neurons decreased to a steady state within the first second of a 10 second depolarizing current injection, whereas the firing rate of mouse ET neurons increased moderately over the same time period (Fig. 7m,n; Extended Data Fig. 10m,n). The acceleration in rodent ET neurons has been attributed to the expression of Kv1-containing voltage-gated K^+^ channels that are encoded by genes like the conserved ET gene *KCNA1*. In macaque and human ET/Betz neurons, a distinctive biphasic pattern was characterized by an early cessation of firing followed by a sustained and dramatic increase in firing later in the current injection. Thus, while ET neurons in both primate and rodent M1 displayed spike frequency acceleration, the temporal dynamics and magnitude of this acceleration appears to be a unique feature of primate ET/Betz neurons. These data emphasize how transcriptomic data from this specialized neuron type can be linked to shared and unique physiological properties across species.

## Discussion

Comparative analysis is a powerful strategy to understand brain structure and function. Species conservation is strong evidence for functional relevance under evolutionary constraints that can help identify critical molecular and regulatory mechanisms ^54, 55^. Conversely, divergence indicates adaptive specialization, which may be essential to understand the mechanistic underpinnings of human brain function and susceptibility to human-specific diseases. In the current study, we applied a comparative approach to understand conserved and species-specific features of M1 at the level of cell types using single-nucleus RNA-seq (Cv3 and SSv4), open chromatin (SNARE-seq2 and ATAC-seq) and DNA-methylation (snmC-seq2) technologies. Integrated analysis of over 450,000 nuclei in human, non-human primates (marmoset, a New World monkey, and to a lesser degree macaque, an Old World monkey that is evolutionarily more closely related to humans), and mouse (see also companion paper^6^) yielded a high-resolution, multimodal classification of cell types in each species, and a coarser consensus classification conserved between rodent and primate lineages. Robust species conservation strongly argues for the functional relevance of this consensus cellular architecture. Species specializations are also apparent, both in the additional granularity in cell types within species and differences between conserved cell types. A comparative evolutionary approach provides an anchor point to define the cellular architecture of any tissue and to discover species-specific adaptations.

A key result of the current study is the identification of a consensus classification of cell types across species that allows the comparison of relative similarities in human compared to common mammalian model organisms in biomedical research. Prior studies have demonstrated that high resolution cellular taxonomies can be generated in mouse, non-human primate and human cortex, and that there is generally good concordance across species ^2, 11^. However, inconsistencies in the methods and sampling depths used made strong conclusions difficult, compounded by the analysis of different cortical regions in different species. The current study overcame these challenges by focusing on M1, a functionally and anatomically conserved cortical region across mammals, and comparing a variety of methods on similarly isolated tissues (and the same specimens from human and marmoset). Several important points emerged from these integrated analyses. First, with deeper sampling and the same methodology (snRNA-seq with Cv3), a similar cellular complexity on the order of 100 cell types was seen in all three species. The highest resolution molecular classification was seen with RNA-seq compared to epigenomic methods, and among RNA-seq methods with those that allow the most cells to be analyzed. Strikingly, the molecular classifications were well aligned across all methods tested, albeit at different levels of resolution as a function of the information content of the assay and the number of cells profiled. All methods were consistent at the level of subclasses as defined above, both across methods and species; significantly better alignment was achieved among species based on transcriptomics, and with epigenomic methods in some subclasses. Mismatches in cellular sampling affect the ability to compare across species; for example, higher non-neuronal sampling in mouse and marmoset increased detection of rare cell types compared to human. One important comparison was between plate-based (SSv4) and droplet-based (Cv3) RNA-seq of human nuclei, where we compared results between approximately 10,000 SSv4 and 100,000 Cv3 nuclei. On average, SSv4 detected 30% more genes per nucleus and enabled comparisons of isoform usage between cell types, albeit with 20-fold greater sequencing depth. However, SSv4 cost 10 times as much as Cv3 and did not allow detection of additional cell types.

The snmC-seq2 clustering aligned closely with the transcriptomic classification, although with significantly lower resolution in rarer subclasses. Hypo-methylated sites correlated with gene expression and specific transcription factor binding motifs were enriched in cell type specific sites. Multi-omic SNARE-seq2 measured RNA profiles of nuclei that allowed high confidence assignment to transcriptomic clusters. Examining accessible chromatin (AC) regions within the same nuclei led to strong correlations between cell subclass or type gene expression and active regulatory regions of open chromatin. Using this strategy, gene regulatory activities could be identified within RNA-defined cell populations (including RNA consensus clusters) that could not be resolved from AC data alone (Extended Data Fig. 6a, Supplementary Table 15). By joint consideration of these epigenomic modalities, glutamatergic neurons were found to have more hypo-methylated DMRs and differentially accessible chromatin, consistent with having larger somata and expressing more genes. Within-species, cell types have many more unique AC sites than uniquely expressed marker genes. At the same time, there is striking conservation across species of subclass TFBS motif enrichment within AC and hypo-methylated DMRs. Most subclasses have distinct motifs, although L2/3 and L6 IT and *Lamp5* and *Sncg* subclasses share many motifs and are more clearly distinguished based on gene expression. Taken together, these results show a robust cell type classification that is consistent at the level of subclasses both across transcriptomic and chromatin measures and across species, with additional cell type-level granularity identified with transcriptomics.

Alignment across species allowed a comparison of relative similarities and differences between species. A common (and expected) theme was that more closely related species are more similar to one another. This was true at the level of gene expression and epigenome patterning across cell types, and in the precision with which transcriptomically-defined cell types could be aligned across species. For example, human and marmoset GABAergic types could be aligned at higher resolution than human and mouse. Human was more similar to macaque than to marmoset. This indicates that cell type similarity increases as a function of evolutionary distance to our closest common ancestors with mouse (∼70 mya), marmoset (∼40 mya), and macaque (∼25 mya). Interestingly, many gene expression differences may change gradually over evolution. This is apparent in the graded changes in expression levels of genes enriched in L5 ET versus L5 IT neurons and in the reduced performance of cell type classification based on marker gene expression that is correlated with evolutionary distance between species.

Several prominent species differences in cell type proportions were observed. First, the ratio of glutamatergic excitatory projection neurons compared to GABAergic inhibitory interneurons was 2:1 in human compared to 3:1 in marmoset and 5:1 in mouse and leads to a profound shift in the overall excitation-inhibition balance of the cortex. A similar species difference has been described based on histological measures (reviewed in ^28^), indicating that snRNA-seq gives a reasonably accurate measurement of cell type proportions. Surprisingly, the relative proportions of GABAergic subclasses and types were similar across species. These results suggest a developmental shift in the size of the GABAergic progenitor pool in the ganglionic eminences or an extended period of neurogenesis and migration. A decreased proportion of the subcortically targeting L5 ET neurons in human was also seen, as previously shown in temporal ^2^ and frontoinsular ^12^ cortex. This shift likely reflects the evolutionary increase in cortical neurons relative to their subcortical targets ^56^ and was less prominent in M1, suggesting regional variation in the proportion of L5 ET neurons. Finally, a large increase in the proportion of L2 and L3 IT neurons was seen in human compared to mouse and marmoset. This increase parallels the disproportionate expansion of human cortical area and supragranular layers that contain neurons projecting to other parts of the cortex, presumably to facilitate greater corticocortical communication. Interestingly, L2 and L3 IT neurons appear to be particularly highly variable across cortical areas and species, and also are more diverse and specialized in human compared to mouse (see companion paper ^44^).

A striking and somewhat paradoxical observation is the high degree of species specialization of consensus types. The majority of DEGs between cell types were consistently species-specific. This result suggests that the conserved cellular features of a cell type are largely due to a minority of DEGs with conserved expression patterns. The current study demonstrates this point for one of the most distinctive brain cell types, the cortical *Pvalb*-expressing GABAergic ChC. ChCs in mouse, non-human primate, and human have 100-150 genes with highly enriched expression compared to other *Pvalb*-expressing interneurons (BCs); however, only 25 of these ChC-enriched genes are shared across species. This small overlapping gene set includes several transcription factors and a member of the netrin family (*UNC5B*) that could be responsible for AIS targeting. Binding sites for these TFs are enriched in ChC cluster regions of open chromatin and in hypo-methylated regions around ChC-enriched genes. While these associations between genes and cellular phenotypes for conserved and divergent features remain to be tested, a comparative strategy can identify these core conserved genes and make strong predictions about the TF code for cell types and the genes responsible for their evolutionarily constrained functions.

M1 is an agranular cortex lacking a L4, although a recent study demonstrated that there are neurons with L4-like properties in mouse ^14^. Here we confirm and extend this finding in human M1. We find a L4-like neuron type in M1 that aligns to a L4 type in human MTG and is scattered between the deep part of L3 and the superficial part of L5 where L4 would be if aggregated into a layer. However, MTG contained several additional L4 types not found in M1, and with a much higher frequency. The human M1 L4-like type is part of the L5 IT_1 consensus cluster that includes several IT types in all species, including two L4-like types in mouse (L4/5 IT_1 and L4/5 IT_2) that also express the canonical L4 marker *Rorb* (see companion paper ^6^). Therefore, it appears that M1 has L4-like cells from a transcriptomic perspective, but only a subset of the types compared to granular cortical areas, at much lower density, and scattered rather than aggregated into a tight layer.

The most distinctive cellular hallmark of M1 in primates and cats is the enormous Betz cell, which contributes to direct corticospinal connections to spinal motoneurons in primates that participate in fine motor control ^15, 16, 57–59^. Intracellular recordings from cats have shown highly distinctive characteristics including HCN channel-related membrane properties, spike frequency acceleration, and extremely fast maintained firing rates ^19, 20^. However, they have never been recorded in primates using patch clamp physiology due to the high degree of myelination in M1 that prevents their visualization, and the inability to obtain motor cortex tissue from neurosurgical procedures which are careful to be function-sparing. A goal of the current project was to identify the transcriptomic cluster corresponding to Betz cells and use this to understand gene expression that may underlie their distinctive properties and species specializations. We have recently taken a similar approach to study von Economo neurons in the fronto-insular cortex, showing they are found within a transcriptomic class consisting of ET neurons ^12^. Betz cells are classical ET neurons that, together with the axons of smaller corticospinal neurons, make up part of the pyramidal tract from the cortex to the spinal cord ^16, 60^. We show that neurons with Betz cell morphology label with markers for the M1 ET clusters. Like von Economo neurons, there does not appear to be an exclusively Betz transcriptomic type. Rather, M1 ET clusters are not exclusive for neurons with Betz morphology, and we find more than one ET cluster contains neurons with Betz morphology.

Although comparative transcriptomic alignments provide strong evidence for functional similarity, the distinctions between corticospinal neurons across species or even between L5 ET and IT neuron types in primates or humans has not been demonstrated physiologically. We recently developed a suite of methodologies for studying specific neuron types in human and non-human tissues, including triple modality Patch-seq to combine physiology, morphology and transcriptome analysis, acute and cultured slice physiology in adult human neurosurgical resections and macaque brain, and AAV-based neuronal labeling to allow targeting of neurons in highly myelinated tissues (companion paper ^44^; ^61^. Specifically, these tools allow the targeting of L5 neurons in mouse and non-human primate and the assignment of neurons to their transcriptomic types using Patch-seq, which we facilitated by generating and aligning a L5 transcriptomic classification in macaque where such analyses could be performed. We show here that several of the characteristic features of L5 ET versus IT neurons are conserved, and can be reliably resolved from one another in mouse and non-human primate. Furthermore, macaque neurons with Betz-like morphologies mapped to the Betz-containing clusters. However, as predicted by differences in ion channel-related gene expression, not all physiological features were conserved between macaque and mouse ET neurons. Betz/ET neurons had the distinctive pauses, bursting and spike-frequency acceleration described previously in cats but not seen in rodents ^19, 20^. Finally, we had access to an extremely rare human neurosurgical case where a region of premotor cortex was resected. Similar to macaque M1, this premotor region contained large neurons with characteristic Betz-like morphology that mapped transcriptomically to the Betz-containing clusters. Together these results highlight the predictive power of transcriptomic mapping and cross-species inference of cell types for L5 pyramidal neurons including the Betz cells. Furthermore, these data are consistent with observations that Betz cells may not in fact be completely restricted to M1 but distribute across other proximal motor-related areas that contribute to the pyramidal tract ^62^. Finally, a number of ion channels that may contribute to conserved ET versus IT features as well as species specializations of Betz cell function were identified that provide candidate genes to explore gene-function relationships. For example, axon guidance-associated genes are enriched in Betz-containing ET neuron types in primates, possibly explaining why Betz cells in primates directly contact spinal motor neurons rather than spinal interneurons as in rodents. Thus, as the comparative approach is helpful in identifying core conserved molecular programs, it may be equally valuable to understand what is different in human or can be well modeled in closer non-human primate relatives. This is particularly relevant in the context of Betz cells and other ET neuron types that are selectively vulnerable in amyotrophic lateral sclerosis, some forms of frontotemporal dementia, and other neurodegenerative conditions.

## Methods

### Ethical compliance

Postmortem adult human brain tissue was collected after obtaining permission from decedent next-of-kin. Postmortem tissue collection was performed in accordance with the provisions of the United States Uniform Anatomical Gift Act of 2006 described in the California Health and Safety Code section 7150 (effective 1/1/2008) and other applicable state and federal laws and regulations. The Western Institutional Review Board reviewed tissue collection processes and determined that they did not constitute human subjects research requiring institutional review board (IRB) review.

### Postmortem human tissue specimens

Male and female donors 18–68 years of age with no known history of neuropsychiatric or neurological conditions (‘control’ cases) were considered for inclusion in the study (Extended Data Table 1). Routine serological screening for infectious disease (HIV, Hepatitis B, and Hepatitis C) was conducted using donor blood samples and only donors negative for all three tests were considered for inclusion in the study. Only specimens with RNA integrity (RIN) values ≥7.0 were considered for inclusion in the study. Postmortem brain specimens were processed as previously described ^2^. Briefly, coronal brain slabs were cut at 1cm intervals and frozen for storage at −80°C until the time of further use. Putative hand and trunk-lower limb regions of the primary motor cortex were identified, removed from slabs of interest, and subdivided into smaller blocks. One block from each donor was processed for cryosectioning and fluorescent Nissl staining (Neurotrace 500/525, ThermoFisher Scientific). Stained sections were screened for histological hallmarks of primary motor cortex. After verifying that regions of interest contained M1, blocks were processed for nucleus isolation as described below.

### Human RNA-sequencing, QC and clustering

#### SMART-seq v4 nucleus isolation and sorting

Vibratome sections were stained with fluorescent Nissl permitting microdissection of individual cortical layers (<u>dx.doi.org/10.17504/protocols.io.7aehibe</u>). Nucleus isolation was performed as previously described (<u>dx.doi.org/10.17504/protocols.io.ztqf6mw</u>). NeuN staining was carried out using mouse anti-NeuN conjugated to PE (FCMAB317PE, EMD Millipore) at a dilution of 1:500. Control samples were incubated with mouse IgG1k-PE Isotype control (BD Biosciences, 555749). DAPI (4’,6-diamidino-2-phenylindole dihydrochloride, ThermoFisher Scientific, D1306) was applied to nuclei samples at a concentration of 0.1µg/ml. Single-nucleus sorting was carried out on either a BD FACSAria II SORP or BD FACSAria Fusion instrument (BD Biosciences) using a 130 µm nozzle. A standard gating strategy based on DAPI and NeuN staining was applied to all samples as previously described ^2^. Doublet discrimination gates were used to exclude nuclei aggregates.

#### SMART-seq v4 RNA-sequencing

The SMART-Seq v4 Ultra Low Input RNA Kit for Sequencing (Takara #634894) was used per the manufacturer’s instructions. Standard controls were processed with each batch of experimental samples as previously described. After reverse transcription, cDNA was amplified with 21 PCR cycles. The NexteraXT DNA Library Preparation (Illumina FC-131-1096) kit with NexteraXT Index Kit V2 Sets A-D (FC-131-2001, 2002, 2003, or 2004) was used for sequencing library preparation. Libraries were sequenced on an Illumina HiSeq 2500 instrument using Illumina High Output V4 chemistry.

#### SMART-seq v4 gene expression quantification

Raw read (fastq) files were aligned to the GRCh38 human genome sequence (Genome Reference Consortium, 2011) with the RefSeq transcriptome version GRCh38.p2 (current as of 4/13/2015) and updated by removing duplicate Entrez gene entries from the gtf reference file for STAR processing. For alignment, Illumina sequencing adapters were clipped from the reads using the fastqMCF program. After clipping, the paired-end reads were mapped using Spliced Transcripts Alignment to a Reference (STAR) using default settings. Reads that did not map to the genome were then aligned to synthetic construct (i.e. ERCC) sequences and the *E. coli* genome (version ASM584v2). Quantification was performed using summerizeOverlaps from the R package GenomicAlignments. Expression levels were calculated as counts per million (CPM) of exonic plus intronic reads.

10x Chromium RNA-sequencing. Nucleus isolation for 10x Chromium RNA-sequencing was conducted as described (dx.doi.org/10.17504/protocols.io.y6rfzd6). After sorting, single-nucleus suspensions were frozen in a solution of 1X PBS, 1% BSA, 10% DMSO, and 0.5% RNAsin Plus RNase inhibitor (Promega, N2611) and stored at −80°C. At the time of use, frozen nuclei were thawed at 37°C and processed for loading on the 10x Chromium instrument as described (dx.doi.org/10.17504/protocols.io.nx3dfqn). Samples were processed using the 10x Chromium Single-Cell 3’ Reagent Kit v3. 10x chip loading and sample processing was done according to the manufacturer’s protocol. Gene expression was quantified using the default 10x Cell Ranger v3 pipeline except substituting the curated genome annotation used for SMART-seq v4 quantification. Introns were annotated as “mRNA”, and intronic reads were included in expression quantification.

Quality control of RNA-seq data. Nuclei were included for analysis if they passed all QC criteria.

SMART-seq v4 criteria:

> 30% cDNA longer than 400 base pairs
> 500,000 reads aligned to exonic or intronic sequence
> 40% of total reads aligned
> 50% unique reads
> 0.7 TA nucleotide ratio
Cv3 criteria:
> 500 (non-neuronal nuclei) or > 1000 (neuronal nuclei) genes detected
< 0.3 doublet score

#### Clustering RNA-seq data

Nuclei passing QC criteria were grouped into transcriptomic cell types using a previously reported iterative clustering procedure (Tasic et al. 2018; Hodge, Bakken et al., 2019). Briefly, intronic and exonic read counts were summed, and log_2_-transformed expression was centered and scaled across nuclei. X- and Y-chromosomes and mitochondrial genes were excluded to avoid nuclei clustering based on sex or nuclei quality. DEGs were selected, principal components analysis (PCA) reduced dimensionality, and a nearest neighbor graph was built using up to 20 principal components. Clusters were identified with Louvain community detection (or Ward’s hierarchical clustering if N < 3000 nuclei), and pairs of clusters were merged if either cluster lacked marker genes. Clustering was applied iteratively to each subcluster until clusters could not be further split.

Cluster robustness was assessed by repeating iterative clustering 100 times for random subsets of 80% of nuclei. A co-clustering matrix was generated that represented the proportion of clustering iterations that each pair of nuclei were assigned to the same cluster. We defined consensus clusters by iteratively splitting the co-clustering matrix as described (Tasic et al. 2018; Hodge, Bakken et al., 2019). The clustering pipeline is implemented in the R package “scrattch.hicat”, and the clustering method is provided by the “run_consensus_clust” function (https://github.com/AllenInstitute/scrattch.hicat).

Clusters were curated based on QC criteria or cell class marker expression (*GAD1*, *SLC17A7*, *SNAP25*). Clusters were identified as donor-specific if they included fewer nuclei sampled from donors than expected by chance. To confirm exclusion, clusters automatically flagged as outliers or donor-specific were manually inspected for expression of broad cell class marker genes, mitochondrial genes related to quality, and known activity-dependent genes.

### Marmoset sample processing and nuclei isolation

Marmoset experiments were approved by and in accordance with Massachusetts Institute of Technology IACUC protocol number 051705020. Two adult marmosets (2.3 and 3.1 years old; one male, one female; Extended Data Table 2) were deeply sedated by intramuscular injection of ketamine (20-40 mg/kg) or alfaxalone (5-10 mg/kg), followed by intravenous injection of sodium pentobarbital (10–30 mg/kg). When pedal withdrawal reflex was eliminated and/or respiratory rate was diminished, animals were transcardially perfused with ice-cold sucrose-HEPES buffer. Whole brains were rapidly extracted into fresh buffer on ice. Sixteen 2-mm coronal blocking cuts were rapidly made using a custom-designed marmoset brain matrix. Coronal slabs were snap-frozen in liquid nitrogen and stored at −80°C until use.

As with human samples, M1 was isolated from thawed slabs using fluorescent Nissl staining (Neurotrace 500/525, ThermoFisher Scientific). Stained sections were screened for histological hallmarks of primary motor cortex. Nuclei were isolated from the dissected regions following this protocol: https://www.protocols.io/view/extraction-of-nuclei-from-brain-tissue-2srged6 and processed using the 10x Chromium Single-Cell 3’ Reagent Kit v3. 10x chip loading and sample processing was done according to the manufacturer’s protocol.

### Marmoset RNA-sequencing, QC and clustering

#### RNA-sequencing

Libraries were sequenced on NovaSeq S2 instruments (Illumina). Raw sequencing reads were aligned to calJac3. Mitochondrial sequence was added into the published reference assembly. Human sequences of RNR1 and RNR2 (mitochondrial) and RNA5S (ribosomal), were aligned using gmap to the marmoset genome and added to the calJac3 annotation. Reads that mapped to exons or introns of each assembly were assigned to annotated genes. Libraries were sequenced to a median read depth of 5.95 reads per unique molecular index (UMI). The alignment pipeline can be found at https://github.com/broadinstitute/Drop-seq.

Cell filtering. Cell barcodes were filtered to distinguish true nuclei barcodes from empty beads and PCR artifacts by assessing proportions of ribosomal and mitochondrial reads, ratio of intronic/exonic reads (> 50% of intronic reads), library size (> 1000 UMIs) and sequencing efficiency (true cell barcodes have higher reads/UMI). The resulting digital gene expression matrix (DGE) from each library was carried forward for clustering.

#### Clustering

Clustering analysis proceeded as in Krienen et al (2019, bioRxiv). Briefly, independent component analysis. (ICA, using the fastICA package in R) was performed jointly on all marmoset DGEs after normalization and variable gene selection as in (Saunders et al 2018, Cell). The first-round clustering resulted in 15 clusters corresponding to major cell classes (neurons, glia, endothelial). Each cluster was curated as in (Saunders et al 2018, Cell) to remove doublets and outliers. Independent components (ICs) were partitioned into those reflecting artifactual signals (e.g. those for which cell loading indicated replicate or batch effects). Remaining ICs were used to determine clustering (Louvain community detection algorithm igraph package in R); for each cluster nearest neighbor and resolution parameters were set to optimize 1:1 mapping between each IC and a cluster.

#### Mouse snRNA-seq data generation and analysis

Single-nuclei were isolated from mouse primary motor cortex, gene expression was quantified using Cv3 and SSv4 RNA-sequencing, and transcriptomic cell types and dendrogram were defined as described in a companion paper ^6^.

#### Integrating and clustering human Cv3 and SSv4 snRNA-seq datasets

To establish a set of human consensus cell types, we performed a separate integration of snRNA-seq technologies on the major cell classes (Glutamatergic, GABAergic, and Non-neuronal). Broadly, this integration is comprised of 6 steps: (1) subsetting the major cell class from each technology (e.g. Cv3 GABAergic and SSv4 GABAergic); (2) finding marker genes for all clusters within each technology; (3) integrating both datasets with Seurat’s standard workflow using marker genes to guide integration (Seurat 3.1.1); (4) overclustering the data to a greater number of clusters than were originally identified within a given individual dataset; (5) finding marker genes for all integrated clusters; and (6) merging similar integrated clusters together based on marker genes until all merging criteria were sufficed, resulting in the final human consensus taxonomy.

More specifically, each expression matrix was log_2_(CPM + 1) transformed then placed into a Seurat object with accompanying metadata. Variable genes were determined by downsampling each expression matrix to a maximum of 300 nuclei per scrattch.hicat-defined cluster (from a previous step; see scrattch.hicat clustering) and running select_markers (scrattch.io 0.1.0) with n set to 20, to generate a list of up to 20 marker genes per cluster. The union of the Cv3 and SSv4 gene lists were then used as input for anchor finding, dimensionality reduction, and Louvain clustering of the full expression matrices. We used 100 dimensions for steps in the workflow, and 100 random starts during clustering. Louvain clustering was performed to overcluster the dataset to identify more integrated clusters than the number of scrattch.hicat-defined clusters. For example, GABAergic neurons had 79 and 37 scrattch.hicat-defined clusters, 225 overclustered integrated clusters, and 72 final human consensus clusters after merging for Cv3 and SSv4 datasets, respectively. To merge the overclustered integrated clusters, up to 20 marker genes were found for each cluster to establish the neighborhoods of the integrated dataset. Clusters were then merged with their nearest neighbor if there were not a minimum of ten Cv3 and two SSv4 nuclei in a cluster, and a minimum of 4 DEGs that distinguished the query cluster from the nearest neighbor (note: these were the same parameters used to perform the initial scrattch.hicat clustering of each dataset).

#### Integrating and clustering MTG and M1 SSv4 snRNA-seq datasets

To compare cell types between our M1 human cell type taxonomy and our previously described human MTG taxonomy ^2^, we used Seurat’s standard integration workflow to perform a supervised integration of the M1 and MTG SSv4 datasets. Intronic and exonic reads were summed into a single expression matrix for each dataset, CPM normalized, and placed into a Seurat object with accompanying metadata. All nuclei from each major cell class were integrated and clustered separately. Up to 100 marker genes for each cluster within each dataset were identified, and the union of these two gene lists was used as input to guide alignment of the two datasets during integration, dimensionality reduction, and clustering steps. We used 100 dimensions for all steps in the workflow.

#### Integrating Cv3 snRNA-seq datasets across species

To identify homologous cell types across species, we used Seurat’s SCTransform workflow to perform a separate supervised integration on each cell class across species. Raw expression matrices were reduced to only include genes with one-to-one orthologs defined in the three species (14,870 genes; downloaded from NCBI Homologene in November, 2019) and placed into Seurat objects with accompanying metadata. To avoid having one species dominate the integrated space and to account for potential differences in each species’ clustering resolution, we downsampled the number of nuclei to have similar numbers across species at the subclass level (e.g. *Lamp5*, *Sst*, L2/3 IT, L6b, etc.). The species with the largest number of clusters under a given subclass was allowed a maximum of 200 nuclei per cluster. The remaining species then split this theoretical maximum (200 nuclei times the max number of clusters under subclass) evenly across their clusters. For example, the L2/3 IT subclass had 8, 4, and 3 clusters for human, marmoset, and mouse, respectively. All species were allowed a maximum of 1600 L2/3 IT nuclei total; or a maximum of 200 human, 400 marmoset, and 533 mouse nuclei per cluster. To integrate across species, all Seurat objects were merged and normalized using Seurat’s SCTransform function. To better guide the alignment of cell types from each species, we found up to 100 marker genes for each cluster within a given species. We used the union of these gene lists as input for integration and dimensionality reduction steps, with 30 dimensions used for integration and 100 for dimensionality reduction and clustering. Clustering the human-marmoset-mouse integrated space provided an additional quality control mechanism, revealing numerous small, species-specific integrated clusters that contained only low-quality nuclei (low UMIs and genes detected). We excluded 4836 nuclei from the marmoset dataset that constituted low-quality integrated neuronal clusters.

To identify which clusters in our three species taxonomy aligned with macaque clusters from our L5 dissected Cv3 dataset, we performed an identical integration workflow on Glutamatergic neurons as was used for the three species integration. Macaque clusters were assigned subclass labels based on their corresponding alignment with subclasses from the other species. The annotated L5 dissected macaque Cv3 dataset was then used as a reference for mapping macaque patch-seq nuclei (see section below).

### Estimation of cell type homology

To identify homologous groups from different species, we applied a tree-based method (https://github.com/AllenInstitute/BICCN_M1_Evo and package: https://github.com/huqiwen0313/speciesTree). In brief, the approach consists of 4 steps: 1) metacell clustering, 2) hierarchical reconstruction of a metacell tree, 3) measurements of species mixing and stability of splits and 4) dynamic pruning of the hierarchical tree.

Firstly, to reduce noise in single-cell datasets and to remove species-specific batch effects, we clustered cells into small highly similar groups based on the integrated matrix generated by Seurat, as described in the previous section. These cells were further aggregated into metacells and the expression values of the metacells were calculated by averaging the gene expression of individual cells that belong to each metacell. Correlation was calculated based on the metacell gene expression matrix to infer the similarity of each metacell cluster. Then hierarchical clustering was performed based on the metacell gene expression matrix using Ward’s method. For each node or corresponding branch in the hierarchical tree, we calculated 3 measurements, and the hierarchical tree was visualized based on these measurements: 1) Cluster size visualized as the thickness of branches in the tree; 2) Species mixing calculated based on entropy of the normalized cell distribution and visualized as the color of each node and branch; 3) Stability of each node. The entropy of cells was calculated as: *H = − ∑_i_ p_i_logp_i_*, where p_i_ is the probability of cells from one species appearing among all the cells in a node. We assessed the node stability by evaluating the agreement between the original hierarchical tree and a result on a subsampled dataset calculated based on the optimal subtree in the subsampled hierarchical trees derived from subsampling 95% of cells in the original dataset. The entire subsampling process was repeated 100 times and the mean stability score for every node in the original tree was calculated. Finally, we recursively searched each node in the tree. If the heuristic criteria (see below) were not met for any node below the upper node, the entire subtree below the upper node was pruned and all the cells belonging to this subtree were merged into one homologous group.

To identify robust homologous groups, we applied criteria in two steps to dynamically search the cross-species tree. Firstly, for each node in the tree, we computed the mixing of cells from 3 species based on entropy and set it as a tuning parameter. For each integrated tree, we tuned the entropy parameter to make sure the tree method generated the highest resolution of homologous clusters without losing the ability to identify potential species-specific clusters. Nodes with entropy larger than 2.9 (for inhibitory neurons) or 2.75 (for excitatory neurons) were considered as well-mixed nodes. For example, an entropy of 2.9 corresponded to a mixture of human, marmoset, and mouse equal to (0.43, 0.37, 0.2) or (0.38, 0.30, 0.32). We recursively searched all the nodes in the tree until we found the node nearest the leaves of the tree that was well-mixed among species, and this node was defined as a well-mixed upper node. Secondly, we further checked the within-species cell composition for the subtrees below the well-mixed upper node to determine if further splits were needed. For the cells on the subtrees below the well mixed upper node, we measured the purity of within-species cell composition by calculating the percentage of cells that fall into a specific sub-group in each individual species. If the purity for any species was larger than 0.8, we went one step further below the well mixed upper node so that its children were selected. Any branches below these nodes (or well-mixed upper node if the within-species cell composition criteria was not met) were pruned and cells from these nodes were merged into the same homologous groups, then the final integrated tree was generated.

As a final curation step, the homologous groups generated by the tree method were merged to be consistent with within-species clusters. We defined consensus types by comparing the overlap of within-species clusters between human and marmoset and human and mouse, as previously described^2^. For each pair of human and mouse clusters and human and marmoset clusters, the overlap was defined as the sum of the minimum proportion of nuclei in each cluster that overlapped within each leaf of the pruned tree. This approach identified pairs of clusters that consistently co-clustered within one or more leaves. Cluster overlaps varied from 0 to 1 and were visualized as a heatmap with human M1 clusters in rows and mouse or marmoset M1 clusters in columns. Cell type homologies were identified as one-to-one, one-to-many, or many-to many so that they were consistent in all three species. For example, the Vip_2 consensus type could be resolved into multiple homologous types between human and marmoset but not human and mouse, and the coarser homology was retained. Consensus type names were assigned based on the annotations of member clusters from human and mouse and avoided specific marker gene names due to the variability of marker expression across species.

To quantify cell type alignment between pairs of species, we pruned the hierarchical tree described above based on the stability and mixing of two species. We performed this analysis for human-marmoset, human-mouse, and marmoset-mouse and compared the alignment resolution of each subclass. The pruning criteria were tuned to fit the two-species comparison and to remove bias, and we set the same criteria for all comparisons (entropy cutoff 3.0). Specifically, for each subclass and pairwise species comparison, we calculated the number of leaves in the pruned tree. We repeated this analysis on the 100 subsampled datasets and calculated the mean and standard deviation of the number of leaves in the pruned trees. For each subclass, we tested for significant differences in the average number of leaves across pairs of species using an ANOVA test followed by post-hoc Tukey HSD tests.

### Marker determination for cell type clusters by NS-Forest v2.1

NS-Forest v2.1 was used to determine the minimum set of marker genes whose combined expression identified cells of a given type with maximum classification accuracy (T. Bakken et al. 2017; Aevermann et al. 2018). (https://github.com/JCVenterInstitute/NSForest/releases). Briefly, for each cluster NS-Forest produces a Random Forest (RF) model using a one vs all binary classification approach. The top ranked genes from RF are then filtered by expression level to retain genes that are expressed in at least 50% of the cells within the target cluster. The selected genes are then reranked by Binary Score calculated by first finding median cluster expression values for a given gene and dividing by the target median cluster expression value. Next, one minus this scaled value is calculated resulting in 0 for the target cluster and 1 for clusters that have no expression, while negative scaled values are set to 0.

These values are then summed and normalized by dividing by the total number of clusters. In the ideal case, where all off-target clusters have no expression, the binary score is 1. Finally, for the top 6 binary genes optimal expression level cutoffs are determined and all permutations of genes are evaluated by f-beta score, where the beta is weighted to favor precision. This f-beta score indicates the power of discrimination for a cluster and a given set of marker genes. The gene combination giving the highest f-beta score is selected as the optimal marker gene combination. Marker gene sets for human, mouse and marmoset primary motor cortex are listed in Supplementary Tables 4, 5, and 6, respectively, and were used to construct the semantic cell type definitions provided in Supplementary Table 1.

### Calculating differentially expressed genes (DEGs)

To identify subclass level DEGs that are conserved and divergent across species, we used the integrated Seurat objects from the species integration step. Seurat objects for each major cell class were downsampled to have up to 200 cells per species cell type. Positive DEGs were then found using Seurat’s FindAllMarkers function using the ROC test with default parameters. We compared each subclass within species to all remaining nuclei in that class and used the SCT normalized counts to test for differential expression. For example, human *Sst* nuclei were compared to all other GABAergic human neurons using the ROC test. Venn diagrams were generated using the eulerr package (6.0.0) to visualize the relationship of DEGs across species for a given subclass. Heatmaps of DEGs for all subclasses under a given class were generated by downsampling each subclass to 50 random nuclei per species. SCT normalized counts were then scaled and visualized with Seurat’s DoHeatmap function.

To identify ChC DEGs that are enriched over BCs, we used the integrated Seurat objects from the species integration step. The *Pvalb* subclass was subset and species cell types were then designated as either ChCs or BCs. Positive DEGs were then found using Seurat’s FindAllMarkers function using the ROC test to compare ChCs and BCs for each species. Venn diagrams were generated using the eulerr package (6.0.0) to visualize the relationship of ChC-enriched DEGs across species.

Heatmaps of conserved DEGs were generated by downsampling the dataset to have 100 randomly selected BCs and ChCs from each species. SCT normalized counts were then scaled and visualized with Seurat’s DoHeatmap function.

We used the four species (human, macaque, marmoset, and mouse) integrated Glutamatergic Seurat object from the species integration step for all L5 ET DEG figures. L5 ET and L5 IT subclasses were downsampled to 200 randomly selected nuclei per species. A ROC test was then performed using Seurat’s FindAllMarkers function between the two subclasses for each species to identify L5 ET-specific marker genes. We then used the UpSetR (1.4.0) package to visualize the intersections of the marker genes across all four species as an upset plot. To determine genes that decrease in expression across evolutionary distance in L5 ET neurons, we found the log-fold change between L5 ET and L5 IT for each species across all genes. We then filtered the gene lists to only include genes that had a trend of decreasing log-fold change (human > macaque > marmoset > mouse). Lastly, we excluded any gene that did not have a log-fold change of 0.5 or greater in the human comparison. These 131 genes were then used as input for GO analysis with the PANTHER Classification System ^67^ for the biological process category, with organism set to Homo sapiens. All significant GO terms for this gene list were associated with cell-cell adhesion and axon-guidance, and are colored blue in the line graph of their expression enrichment.

### Estimating differential isoform usage between human and mouse

To assess changes of isoform usage between mouse and human, we used SSv4 data with full transcript coverage and estimated isoform abundance in each cell subclasses. To mitigate low read depth in each cell, we aggregated reads from all cells in each subclass. We estimated the relative isoform usage in each subclass by calculating its genic proportion (P), defined as the ratio (R) of isoform expression to the gene expression, where R = (P_human_ - P_mouse_) / (P_human_ + P_mouse_). For a common set of transcripts for mouse and human, we used UCSC browser TransMapV5 set of human transcripts (hg38 assembly, Gencode v31 annotations) mapped to the mouse genome (mm10 assembly) http://hgdownload.soe.ucsc.edu/gbdb/mm10/transMap/V5/mm10.ensembl.transMapV5.bigPsl. We considered only medium to highly expressed isoforms, which have abundance > 10 TPM (Transcripts per Million) and P > 0.2 in either mouse or human and gene expression > 10 TPM in both mouse and human.

Calculating isoform abundance in each cell subclass:

1. Aggregated reads from each subclass
2. Mapped reads to the mouse or human reference genome with STAR 2.7.3a using default parameters
3. Transformed genomic coordinates into transcriptomic coordinates using STAR parameter: -- quantMode TranscriptomeSAM
4. Quantified isoform and gene expression using RSEM 1.3.3 parameters: --bam --seed 12345 -- paired-end --forward-prob 0.5 --single-cell-prior --calc-ci

Estimating statistical significance:

Calculated the standard deviation of isoform genic proportion (P_human_ and P_mouse_) from the RSEM’s 95% confidence intervals of isoform expression Calculated the P-value using normal distribution for the (P_human_ - P_mouse_) and the summed (mouse + human) variance Bonferroni-adjusted P-values by multiplying nominal P-values by the number of medium to highly expressed isoforms in each subclass

### Species cluster dendrograms

DEGs for a given species were identified using Seurat’s FindAllMarkers function with a Wilcox test and comparing each cluster to every other cluster under the same subclass, with logfc.threshold set to 0.7 and min.pct set to 0.5. The union of up to 100 genes per cluster with the highest avg_logFC were used.The average log_2_ expression of the DEGs were then used as input for the build_dend function from scrattch.hicat to create the dendrograms. This was performed on both human and marmoset datasets. For mouse dendrogram methods, see the companion paper ^6^.

### Multiplex fluorescent in situ hybridization (FISH)

Fresh-frozen human postmortem brain tissues were sectioned at 14-16 μm onto Superfrost Plus glass slides (Fisher Scientific). Sections were dried for 20 minutes at −20°C and then vacuum sealed and stored at −80°C until use. The RNAscope multiplex fluorescent v1 kit was used per the manufacturer’s instructions for fresh-frozen tissue sections (ACD Bio), except that fixation was performed for 60 minutes in 4% paraformaldehyde in 1X PBS at 4°C and protease treatment was shortened to 5minutes. Primary antibodies were applied to tissues after completion of mFISH staining. Primary antibodies used were mouse anti-GFAP (EMD Millipore, MAB360, 1:250 dilution) and mouse anti-Neurofilament H (SMI-32, Biolegend, 801701). Secondary antibodies were goat anti-mouse IgG (H+L) Alexa Fluor conjugates (594, 647). Sections were imaged using a 60X oil immersion lens on a Nikon TiE fluorescence microscope equipped with NIS-Elements Advanced Research imaging software (version 4.20). For all RNAscope mFISH experiments, positive cells were called by manually counting RNA spots for each gene. Cells were called positive for a gene if they contained ≥ 3 RNA spots for that gene. Lipofuscin autofluorescence was distinguished from RNA spot signal based on the larger size of lipofuscin granules and broad fluorescence spectrum of lipofuscin.

### Gene family conservation

To investigate the conservation and divergence of gene family coexpression between primates and mouse, MetaNeighbor analysis ^30^ was performed using gene groups curated by the HUGO Gene Nomenclature Committee (HGNC) at the European Bioinformatics Institute (https://www.genenames.org; downloaded January 2020) and by the Synaptic Gene Ontology (SynGO) ^68^ (downloaded February 2020). HGNC annotations were propagated via the provided group hierarchy to ensure the comprehensiveness of parent annotations. Only groups containing five or more genes were included in the analysis.

After splitting data by class, MetaNeighbor was used to compare data at the cluster level using labels from cross-species integration with Seurat. Cross-species comparisons were performed at two levels of the phylogeny: 1) between the two primate species, marmoset and human; and 2) between mouse and primates. In the first case, the data from the two species were each used as the testing and training set across two folds of cross-validation, reporting the average performance (AUROC) across folds. In the second case, the primate data were used as an aggregate training set, and performance in mouse was reported. Results were compared to average within-species performance.

### Replicability of clusters

MetaNeighbor was used to provide a measure of neuronal subclass and cluster replicability within and across species. For this application, we tested all pairs of species (human-marmoset, marmoset-mouse, human-mouse) as well as testing within each species. After splitting the data by class, highly variable genes were identified using the get_variable_genes function from MetaNeighbor, yielding 928 genes for GABAergic and 763 genes for Glutamatergic neuron classes, respectively. These were used as input for the MetaNeighborUS function, which was run using the fast_version and one_vs_best parameters set to TRUE. Using the one_vs_best parameter means that only the two closest neighboring clusters are tested for their similarity to the training cluster, with results reported as the AUROC for the closest neighbor over the second closest. AUROCs are plotted in heatmaps in Extended Data Figures 2 and 3. Data to reproduce these figures can be found in Supplementary Table 9, and scripts are on GitHub (http://github.com/gillislab/MetaNeighbor).

### Single-cell methylome data (snmC-seq2): Sequencing and quantification

Library preparation and Illumina sequencing. Single nuclei were isolated from human and marmoset M1 tissue as described above for RNA-seq profiling and for mouse as detailed in ^6^. Detailed methods for bisulfite conversion and library preparation were previously described for snmC-seq2^5, 41^. The snmC-seq2 libraries generated from mouse brain tissues were sequenced using an Illumina Novaseq 6000 instrument with S4 flowcells and 150 bp paired-end mode.

### Mapping and feature count pipeline

We implemented a versatile mapping pipeline (http://cemba-data.rtfd.io) for all the single-cell methylome based technologies developed by our group ^5, 41, 69^. The main steps of this pipeline included: 1) demultiplexing FASTQ files into single-cell; 2) reads level QC; 3) mapping; 4) BAM file processing and QC; and 5) final molecular profile generation. The details of the five steps for snmC-seq2 were described previously ^41^. We mapped all the reads from the three corresponding species onto the human hg19 genome, the marmoset ASM275486v1 genome, and the mouse mm10 genome. After mapping, we calculated the methyl-cytosine counts and total cytosine counts for two sets of genome regions in each cell: the non-overlapping chromosome 100-kb bins of each genome, the methylation levels of which were used for clustering analysis, and the gene body regions, the methylation levels of which were used for cluster annotation and integration with RNA expression data.

### snmC-seq2: Quality control and preprocessing

Cell filtering. We filtered the cells based on these main mapping metrics: 1) mCCC rate < 0.03. mCCC rate reliably estimates the upper bound of bisulfite non-conversion rate ^5^; 2) overall mCG rate > 0.5; 3) overall mCH rate < 0.2; 4) total final reads > 500,000; and 5) bismark mapping rate > 0.5. Other metrics such as genome coverage, PCR duplicates rate, and index ratio were also generated and evaluated during filtering. However, after removing outliers with the main metrics 1-5, few additional outliers can be found.

Feature filtering. 100kb genomic bin features were filtered by removing bins with mean total cytosine base calls < 250 or > 3000. Regions overlap with the ENCODE blacklist ^70^ were also excluded from further analysis.

Computation and normalization of the methylation rate. For CG and CH methylation, the computation of methylation rate from the methyl-cytosine and total cytosine matrices contains two steps: 1) prior estimation for the beta-binomial distribution and 2) posterior rate calculation and normalization per cell. Step 1. For each cell we calculated the sample mean, *m*, and variance, *v*, of the raw mc rate *(mc / cov)* for each sequence context (CG, CH). The shape parameters (*α*, *β*) of the beta distribution were then estimated using the method of moments:

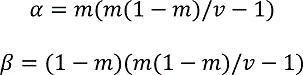

This approach used different priors for different methylation types for each cell and used weaker prior to cells with more information (higher raw variance).

Step 2. We then calculated the posterior: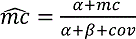., We normalized this rate by the cell’s global mean methylation, *m* = *α*/(*α* + *β*). Thus, all the posterior *mc*˄ with 0 *cov* will be constant 1 after normalization. The resulting normalized *mc* rate matrix contains no NA (not available) value, and features with less *cov* tend to have a mean value close to 1.

Selection of highly variable features. Highly variable methylation features were selected based on a modified approach using the scanpy package *scanpy.pp.highly_variable_genes* function ^71^. In brief, the *scanpy.pp.highly_variable_genes* function normalized the dispersion of a gene by scaling with the mean and standard deviation of the dispersions for genes falling into a given bin for mean expression of genes. In our modified approach, we reasoned that both the mean methylation level and the mean *cov* of a feature (100kb bin or gene) could impact *mc* rate dispersion. We grouped features that fall into a combined bin of mean and *cov*, and then normalized the dispersion within each *mean-cov* group. After dispersion normalization, we selected the top 3000 features based on normalized dispersion for clustering analysis.

Dimension reduction and combination of different mC types. For each selected feature, *mc* rates were scaled to unit variance, and zero mean. PCA was then performed on the scaled *mc* rate matrix. The number of significant PCs was selected by inspecting the variance ratio of each PC using the elbow method. The CH and CG PCs were then concatenated together for further analysis in clustering and manifold learning.

### snmC-seq2: Data analysis

Consensus clustering on concatenated PCs. We used a consensus clustering approach based on multiple Leiden-clustering ^72^ over K-Nearest Neighbor (KNN) graph to account for the randomness of the Leiden clustering algorithms. After selecting dominant PCs from PCA in both mCH and mCG matrix, we concatenated the PCs together to construct a KNN graph using *scanpy.pp.neighbors* with Euclidean distance. Given fixed resolution parameters, we repeated the Leiden clustering 300 times on the KNN graph with different random starts and combined these cluster assignments as a new feature matrix, where each single Leiden result is a feature. We then used the outlier-aware DBSCAN algorithm from the scikit-learn package to perform consensus clustering over the Leiden feature matrix using the hamming distance. Different epsilon parameters of DBSCAN are traversed to generate consensus cluster versions with the number of clusters that range from minimum to the maximum number of clusters observed in the multiple Leiden runs. Each version contained a few outliers that usually fall into three categories: 1) cells located between two clusters that had gradient differences instead of clear borders; 2) cells with a low number of reads that potentially lack information in essential features to determine the specific cluster; and 3) cells with a high number of reads that were potential doublets. The amount of type 1 and 2 outliers depends on the resolution parameter and is discussed in the choice of the resolution parameter section. The type 3 outliers were very rare after cell filtering. The supervised model evaluation then determined the final consensus cluster version.

Supervised model evaluation on the clustering assignment. For each consensus clustering version, we performed a Recursive Feature Elimination with Cross-Validation (RFECV) ^73^ process from the scikit-learn package to evaluate clustering reproducibility. We first removed the outliers from this process, and then we held out 10% of the cells as the final testing dataset. For the remaining 90% of the cells, we used tenfold cross-validation to train a multiclass prediction model using the input PCs as features and *sklearn.metrics.balanced_accuracy_score* ^74^ as an evaluation score. The multiclass prediction model is based on *BalancedRandomForestClassifier* from the imblearn package that accounts for imbalanced classification problems^75^. After training, we used the 10% testing dataset to test the model performance using the *balanced_accuracy_score* score. We kept the best model and corresponding clustering assignments as the final clustering version. Finally, we used this prediction model to predict outliers’ cluster assignments, we rescued the outlier with prediction probability > 0.3, otherwise labeling them as outliers.

Choice of resolution parameter. Choosing the resolution parameter of the Leiden algorithm is critical for determining the final number of clusters. We selected the resolution parameter by three criteria: 1. The portion of outliers < 0.05 in the final consensus clustering version. 2. The ultimate prediction model accuracy > 0.95. 3. The average cell per cluster ≥ 30, which controls the cluster size to reach the minimum coverage required for further epigenome analysis such as DMR calls. All three criteria prevented the over-splitting of the clusters; thus, we selected the maximum resolution parameter under meeting the criteria using a grid search.

Three-level of iterative clustering analysis. We used an iterative approach to cluster the data into three levels of categories with the consensus clustering procedure described above. In the first level termed CellClass, clustering analysis is done on all cells. The resulting clusters are then manually merged into three canonical classes, glutamatergic neurons, GABAergic neurons, and non-neurons, based on marker genes. The same clustering procedure was then conducted within each CellClass to get clusters as the MajorType level. Within each MajorType, we got the final clusters as the SubTypes in the same way.

### Integrating cell clusters identified from snmC-seq2 and from Cv3

We identified gene markers based on gene body mCH hypo-methylation for each level of clustering of snmC-seq2 data using our in-house analysis utilities (https://github.com/lhqing/cemba_data), and identified gene markers for cell class from Cv3 analysis using scanpy ^71^. We then used Scanorama ^76^ to integrate the two modalities.

### Calling CG differentially methylated regions (DMRs)

We identified CG DMRs using methylpy (https://github.com/yupenghe/methylpy) as previously described ^77^. Briefly, we first called CG differentially methylated sites and then merged them into blocks if they both showed similar sample-specific methylation patterns and were within 250bp. Normalized relative lengths of DMRs (Figure 4d) were calculated by summation of lengths of DMRs and 250bp around divided by numbers of cytosine covered in sequencing.

TFBS motif enrichment analysis. For each cell subclass (cluster), we performed TFBS motif enrichment analysis for its hypo-methylated DMRs against the hypo-methylated DMRs from other cell subclasses (clusters) using software AME ^78^. DMRs and 250bp regions around were used in the analysis.

### SNARE-Seq2: Sample preparation

Human and marmoset primary motor cortex nuclei were isolated for SNARE-seq2 according to the following protocol: https://www.protocols.io/view/nuclei-isolation-for-snare-seq2-8tvhwn6 ^7, 79^. Fluorescence-activated nuclei sorting (FANS) was then performed on a FACSAria Fusion (BD Biosciences, Franklin Lakes, NJ) gating out debris from FSC and SSC plots and selecting DAPI^+^ singlets (Extended Data Fig. 5a). Samples were kept on ice until sorting was complete and were used immediately for SNARE-seq2.

### SNARE-Seq2: Library preparation and sequencing

A detailed step-by-step protocol for SNARE-Seq2 has been outlined in a companion paper ^38^. The resulting AC libraries were sequenced on MiSeq (Illumina) (R1: 75 cycles for the 1^st^ end of AC DNA read, R2: 94 cycles for cell barcodes and UMI read, R3: 8 cycles for i5 read, R4: 75 cycles for the 2^nd^ end of AC DNA read) for library validation, then on NovaSeq6000 (Illumina) using 300 cycles reagent kit for data generation. RNA libraries were combined at equimolar ratio and sequenced on MiSeq (Illumina) (Read 1: 70 cycles for the cDNA read, Index 1: 6 cycles for i7 read, Read 2: 94 cycles for cell barcodes and UMI read) for library validation, then on NovaSeq6000 (Illumina) using 200 cycles reagent kit for data generation.

### SNARE-Seq2: Data processing

A detailed step-by-step SNARE-seq2 data processing pipeline has been provided in a companion paper ^38^. For RNA data, this has involved the use of dropEst to extract cell barcodes and STAR (v2.5.2b) to align tagged reads to the genome (GRCh38 version 3.0.0 for human; GCF 000004665.1 Callithrix jacchus-3.2, marmoset). For AC data, this involved snaptools for alignment to the genome (cellranger-atac-GRCh38-1.1.0 for human, GCF 000004665.1 Callithrix jacchus-3.2, marmoset) and to generate snap objects for processing using the R package snapATAC.

### SNARE-Seq2: Data analysis

#### RNA quality filtering

For SNARE-Seq2 data, quality filtering of cell barcodes and clustering analysis were first performed on transcriptomic (RNA) counts and used to inform on subsequent accessible chromatin quality filtering and analysis. Each cell barcode was tagged by an associated library batch ID (for example MOP1, MOP2… etc.), RNA read counts associated with dT and n6 adaptor primers were merged, libraries were combined for each sample within each experiment and empty barcodes removed using the emptyDrops() function of DropletUtils ^80^, mitochondrial transcripts were removed, doublets were identified using the DoubletDetection software ^81^ and removed. All samples were combined across experiments within species and cell barcodes having greater than 200 and less than 7500 genes detected were kept for downstream analyses. To further remove low quality datasets, a gene UMI ratio filter (gene.vs.molecule.cell.filter) was applied using Pagoda2 (https://github.com/hms-dbmi/pagoda2).

#### RNA data clustering

For human SNARE-seq2 RNA data, clustering analysis was first performed using Pagoda2 where counts were normalized to the total number per nucleus and batch variations were corrected by scaling expression of each gene to the dataset-wide average. After variance normalization, the top 6000 over-dispersed genes were used for principal component analysis. Clustering was performed using an approximate k-nearest neighbor graph (k values between 50 – 500) based on the top 75 principal components and cluster identities were determined using the infomap community detection algorithm. Major cell types were identified using a common set of broad cell type marker genes: *GAD1/GAD2* (GABAergic neurons), *SLC17A7/SATB2* (glutamatergic neurons), *PDGFRA* (oligodendrocyte progenitor cells), *AQP4* (astrocytes), *PLP1/MOBP* (oligodendrocytes), *MRC1* (perivascular macrophages), *PTPRC* (T cells), *PDGFRB* (vascular smooth muscle cells), *FLT1* (vascular endothelial cells), *DCN* (vascular fibroblasts), *APBB1IP* (microglia) (Extended Data Fig. 5c). Low quality clusters that showed very low gene/UMI detection rates, low marker gene detection and/or mixed cell type marker profiles were removed. Oligodendrocytes were over-represented (54,080 total), possibly reflecting a deeper subcortical sampling then intended, therefore, to ensure a more balanced distribution of cell types, we capped the number of oligodendrocytes at 5000 total and repeated the PAGODA2 clustering as above. To achieve optimal clustering of the different cell types, different k values were used to identify cluster subpopulations for different cell types (L2/3 glutamatergic neurons, k = 500; all other glutamatergic neurons, astrocytes, oligodendrocytes, OPCs, k = 100; GABAergic neurons, vascular cells, microglia/perivascular macrophages, k = 50). To assess the appropriateness of the chosen k values, clusters were compared against SMARTer clustering of data generated on human M1 through correlation of cluster-averaged scaled gene expression values using the corrplot package (https://github.com/taiyun/corrplot) (Extended Data Fig. 5d). For cluster visualization, uniform manifold approximation and projection (UMAP) dimensional reduction was performed in Seurat (version 3.1.0) using the top 75 principal components identified using Pagoda2. For marmoset, clustering was initially performed using Seurat, where the top 2000 variable features were selected from the mean variance plot using the ‘vst’ method and used for principal component analysis. UMAP embeddings were generated using the top 75 principal components. To harmonize cellular populations across platforms and modalities, snRNA-seq within-species cluster identities were then predicted from both human and marmoset data. We used an iterative nearest centroid classifier algorithm (Methods, ‘Mapping of samples to reference taxonomies’) to generate probability scores for each SNARE-seq2 nuclei mapping to their respective species’ snRNA-seq reference cluster (Cv3 for marmoset and SMART-Seqv4 for human). Comparing the predicted RNA cluster assignment of each nuclei with Pagoda2-identified clusters showed highly consistent cluster membership using Jaccard similarity index (Extended Data Fig. 5e), confirming the robustness of these cell identities discovered using different analysis platforms.

#### AC quality filtering and peak calling

Initial analysis of corresponding SNARE-Seq2 chromatin accessibility data was performed using SnapATAC software (version 2) (https://github.com/r3fang/SnapATAC) (https://doi.org/10.1101/615179). Snap objects were generated by combining individual snap files across libraries within each species. Cell barcodes were included for downstream analyses only if cell barcodes passed RNA quality filtering (above) and showed greater than 1000 read fragments and 500 UMI. Read fragments were then binned to 5000 bp windows of the genome and only cell barcodes showing the fraction of binned reads within promoters greater than 10% (15% for marmoset) and less than 80% were kept for downstream analysis. Peak regions were called independently for RNA cluster, subclass and class groupings using MACS2 software (https://github.com/taoliu/MACS) using the following options “--nomodel --shift 100 --ext 200 --qval 5e-2-B --SPMR”. Peak regions were combined across peak callings and used to generate a single peak count matrix (cell barcodes by chromosomal peak locations) using the “createPmat” function of SnapATAC.

#### AC data clustering

The peak count matrices were filtered to keep only locations from chromosomes 1-22, x or y, and processed using Seurat (version 3.1.0) and Signac (version 0.1.4) software ^24^ (https://satijalab.org). All peaks having at least 100 counts (20 for marmoset) across cells were used for dimensionality reduction using latent semantic indexing (“RunLSI” function) and visualized by UMAP using the first 50 dimensions (40 for marmoset).

#### Calculating gene activity scores

For a gene activity matrix from accessibility data, cis-co-accessible sites and gene activity scores were calculated using Cicero software (v1.2.0) ^39^ (https://cole-trapnell-lab.github.io/cicero-release/). The binary peak matrix was used as input with expression family variable set to “binomialff” to make the aggregated input Cicero CDS object using the AC peak-derived UMAP coordinates and setting 50 cells to aggregate per bin. Co-accessible sites were then identified using the “run_cicero” function using default settings and modules of cis-co-accessible sites identified using the “generate_ccans” function. Co-accessible sites were annotated to a gene if they fell within a region spanning 10,000 bp upstream and downstream of the gene’s transcription start site (TSS). The Cicero gene activity matrix was then calculated using the “build_gene_activity_matrix” function using a co-accessibility cutoff of 0.25 and added to a separate assay of the Seurat object.

#### Integrating RNA/AC data modalities

For reconciliation of differing resolutions achievable from RNA and accessible chromatin (Extended Data Fig. 5f-k), integrative analysis was performed using Seurat. Transfer anchors were identified between the activity and RNA matrices using the “FindTransferAnchors” function. For human, transfer anchors were generated using an intersected list of variable genes identified from Pagoda2 analysis of RNA clusters (top 2000 genes) and marker genes for clusters identified from SSv4 data (2492 genes having β-scores > 0.4), and canonical correlation analysis (CCA) for dimension reduction. For marmoset, transfer anchors were generated using an intersected list of variable genes identified using Seurat (top 2000 genes) and DEGs identified between marmoset consensus clusters (Cv3 snRNA-seq data, P < 0.05, top 100 markers per cluster). Imputed RNA expression values were then calculated using the “TransferData” function from the Cicero gene activity matrix using normalized RNA expression values for reference and LSI for dimension reduction. RNA and imputed expression data were merged, a UMAP co-embedding and shared nearest neighbor (SNN) graph generated using the top 50 principal components (40 for marmoset) and clusters identified (“FindClusters”) using a resolution of 4. Resulting integrated clusters were compared against consensus RNA clusters by calculating jaccard similarity scores using scratch.hicat software. Cell populations identified as T-cells from Pagoda2 analysis (human only) and those representing low quality integrated clusters, showing a mixture of disparate cell types, were removed from these analyses. RNA clusters were assigned to co-embedded clusters based on the highest jaccard similarity score and frequency and then merged to generate the best matched co-embedded clusters, taking in account cell type and subclass to ensure more accurate merging of ambiguous populations. This enabled AC-level clusters that directly matched the RNA-defined populations (Extended Data Fig. 5k). For consensus cluster and subclass level predictions (Extended Data Fig. 5g) the Seurat “TransferData” function was used to transfer RNA consensus cluster or subclass labels to AC data using the pre-computed transfer anchors and LSI dimensionality reduction.

#### Final AC peak and gene activity matrices

A final combined list of peak regions was then generated using MACS2 as detailed above for all cell populations corresponding to RNA consensus (> 100 nuclei), accessibility-level, subclass (> 50 nuclei) and class level barcode groupings. The corresponding peak by cell barcode matrix generated by SnapATAC was used to establish a Seurat object as outlined above, with peak counts, Cicero gene activity scores and RNA expression values for matched cell barcodes contained within different assay slots.

#### Transcription factor motif analyses

Jaspar motifs (JASPAR2020, all vertebrate) were used to generate a motif matrix and motif object that was added to the Seurat object using Signac (“CreateMotifMatrix”, “CreateMotifObject”, “AddMotifObject”) and GC content, region lengths and dinucleotide base frequencies calculated using the “RegionStats” function. Motif enrichments within specific chromosomal sites were calculated using the FindMotifs function. For motif activity scores, chromVAR (https://greenleaflab.github.io/chromVAR) was performed according to default parameters (marmoset) or using Signac “RunChromVAR” function on the peak count matrix (human). The chromVAR deviation score matrix was then added to a separate assay slot of the Seurat object and differential activity of TFBS between different populations were assessed using the “Find[All]Markers” function through logistic regression and using the number of peak counts as a latent variable.

Differentially accessible regions (DARs) between cell populations (Fig. 4b) were identified using the “find_all_diff” function (https://github.com/yanwu2014/chromfunks) and p-values calculated using a hypergeometric test. For visualization, the top DARs (q value < 0.001 and log-fold change > 1) were selected and the top distinct sites visualized by dot plot in Seurat. For motif enrichment analyses, peak counts associated with the clusters selected for comparison (all subclasses, all AC-level clusters, *PVALB*-positive for ChC analyses) were used to identify cis-co-accessible site networks or CCANs using cicero as indicated above. Peak locations were annotated to the nearest gene (10,000 bases upstream and downstream of the TSS) and only genes identified from SNARE-seq2 RNA data as being differentially expressed (Seurat, Wilcoxon Rank Sum test) within the clusters of interest (adjusted P < 0.05, average log-fold change > 0.5) were used. Genes having more than one co-accessible site were assessed for motif enrichments within all overlapping sites using the “FindMotifs” function in Signac (using peaks for all cell barcodes for subclass and AC-level, or only peaks for ChC or L5 ET cells).

Motifs were then trimmed to only those showing significant differential activity (chromVAR) between the clusters of interest (P < 0.05) as assessed using the “FindMarkers” function on the chromVAR assay slot using Seurat and using the number of total peaks as a latent variable. The top distinct genes (subclass, AC-level) or all genes (ChC, Betz) used for motif enrichment analysis were visualized for scaled average RNA expression levels and scaled average cicero gene activities using the ggHeat plotting function (SWNE package, https://github.com/yanwu2014/swne). Top chromVAR TFBS activities were also visualized using ggHeat.

#### Correlation plots

For correlation of RNA expression and associated AC activities for consensus and AC-level clusters (Extended Data Fig. 6a-b), average scaled expression values were generated and pairwise correlations performed for marker genes identified from an intersected list of variable genes identified from Pagoda2 analysis of RNA clusters (top 2000 genes) and marker genes for clusters identified from SSv4 data (2492 genes having β-scores > 0.4). For correlation across species, expression values for genes used to integrate human and marmoset GABAergic and glutamatergic clusters (Cv3 scRNA-seq data), or chromVAR TFBS activity scores for all Jaspar motifs were averaged by subclass, scaled (trimming values to a minimum of 0 and a maximum of 4) for each species separately, then correlated and visualized using corrplot.

#### Plots and figures

All UMAP, feature, dot, and violin plots were generated using Seurat. Connection plots were generated using cicero and peak track gradient heatmaps were generated using Gviz ^82^ from bedGraph files generated during peak calling using SnapATAC. Correlation plots were generated using the corrplot package.

#### Mouse chandelier cell ATAC-Seq: Data acquisition and analysis

Chandelier cells are rare in mouse cortex and were enriched by isolating individual neurons from transgenically-labelled mouse primary visual cortex (VISp). Many of the transgenic mouse lines have previously been characterized by single-cell RNA-seq ^1^. Single-cell suspensions of cortical neurons were generated as described previously ^1^ and subjected to tagmentation (ATAC-seq) ^83, 84^. Mixed libraries, containing 60 to 96 samples were sequenced on an Illumina MiSeq. In total, 4,275 single-cells were collected from 36 driver-reporter combinations in 67 mice. After sequencing, raw FASTQ files were aligned to the GRCm38 (mm10) mouse genome using Bowtie v1.1.0 as previously described ^9^.

Following alignment, duplicate reads were removed using samtools rmdup, which yielded only single copies of uniquely mapped paired reads in BAM format. Quality control filtering was applied to select samples with >10,000 uniquely mapped paired-end fragments, >10% of which were longer than 250 base pairs and with >25% of their fragments overlapping high-depth cortical DNase-seq peaks from ENCODE ^85^. The resulting dataset contained a total of 2,799 samples.

To increase the cell-type resolution of chromatin accessibility profiles beyond that provided by driver lines, a feature-free method for computation of pairwise distances (Jaccard) was used. Using Jaccard distances, principal component analysis (PCA) and t-distributed stochastic neighbor embedding (t-SNE) were performed, followed by Phenograph clustering ^86^. This clustering method grouped cells from class-specific driver lines together, but also segregated them into multiple clusters. Phenograph-defined neighborhoods were assigned to cell subclasses and clusters by comparison of accessibility near transcription start site (TSS ± 20 kb) to median expression values of scRNA-seq clusters at the cell type and at the subclass level from mouse primary visual cortex ^87^. From this analysis, a total of 226 samples were assigned to *Pvalb* and 124 samples to *Pvalb Vipr2* (ChC) clusters. The sequence data for these samples were grouped together and further processed through the Snap-ATAC pipeline.

Mouse scATAC-seq peak counts for *Pvalb* and ChC were used to generate a Seurat object as outlined for human and marmoset SNARE-Seq2 AC data. Cicero cis-co-accessible sites were identified, gene activity scores calculated, and motif enrichment analyses performed as outlined above. Genes used for motif enrichment were ChC markers identified from differential expression analysis between *PVALB-positive* clusters in mouse Cv3 scRNA-seq data (adjusted P < 0.05).

### Patch-seq neuronal physiology, morphology, and transcriptomics

#### Subjects

The human neurosurgical specimen was obtained from a 61-year old female patient that underwent deep tumor resection (glioblastoma) from the frontal lobe at a local hospital (Harborview Medical Center). The patient provided informed consent and experimental procedures were approved by the hospital institute review board before commencing the study. Post-hoc analysis revealed that the neocortical tissue obtained from this patient was from a premotor region near the confluence of the superior frontal gyrus and the precentral gyrus (Fig. 7g). All procedures involving macaques and mice were approved by the Institutional Animal Care and Use Committee at either the University of Washington or the Allen Institute for Brain Science. Macaque M1 tissue was obtained from male (n=4) and female (n=5) animals (mean age= 10 ± 2.21 years) designated for euthanasia via the Washington National Primate Research Center’s Tissue Distribution Program. Mouse M1 tissue was obtained from 4-12 week old male and female mice from the following transgenic lines: *Thy1h*-eyfp (B6.Cg-Tg(*Thy1*-YFP)-HJrs/J: JAX Stock No. 003782), *Etv1*-egfp Tg(*Etv1*-EGFP)BZ192Gsat/Mmucd (etv1) mice maintained with the outbred Charles River Swiss Webster background (Crl:CFW(SW) CR Stock No. 024), and C57BL/6-Tg(*Pvalb*-tdTomato)15Gfng/J: JAX stock No. 027395.

#### Brain slice preparation

Brain slice preparation was similar for *Pvalb*-TdTomato mice, macaque and human samples. Upon resection, human neurosurgical tissue was immediately placed in a chilled and oxygenated solution formulated to prevent excitotoxicity and preserve neural function ^88^. This artificial cerebral spinal fluid (NMDG aCSF) consisted of (in mM): 92 with N-methyl-D-glucamine (NMDG), 2.5 KCl, 1.25 NaH_2_PO_4_, 30 NaHCO_3_, 20 4-(2-hydroxyethyl)-1-piperazineethanesulfonic acid (HEPES), 25 glucose, 2 thiourea, 5 Na-ascorbate, 3 Na-pyruvate, 0.5 CaCl_2_·4H_2_O and 10 MgSO_4_·7H_2_O. The pH of the NMDG aCSF was titrated to pH 7.3–7.4 with concentrated hydrochloric acid and the osmolality was 300-305 mOsmoles/Kg. The solution was pre-chilled to 2-4°C and thoroughly bubbled with carbogen (95% O_2_/5% CO_2_) prior to collection. Macaques were anesthetized with sevoflurane gas during which the entire cerebrum was extracted and placed in the same protective solution described above. After extraction, macaques were euthanized with sodium-pentobarbital. We dissected the trunk/limb area of the primary motor cortex for brain slice preparation. *Pvalb*-TdTomato mice were deeply anesthetized by intraperitoneal administration of Advertin (20mg/kg IP) and were perfused through the heart with NMDG aCSF (bubbled with carbogen).

Brains were sliced at 300-micron thickness on a vibratome using the NMDG protective recovery method and a zirconium ceramic blade ^61, 88^. Mouse brains were sectioned coronally, and human and macaque brains were sectioned such that the angle of slicing was perpendicular to the pial surface. After sections were obtained, slices were transferred to a warmed (32-34° C) initial recovery chamber filled with NMDG aCSF under constant carbogenation. After 12 minutes, slices were transferred to a chamber containing an aCSF solution consisting of (in mM): 92 NaCl, 2.5 KCl, 1.25 NaH_2_PO_4_, 30 NaHCO_3_, 20 HEPES, 25 glucose, 2 thiourea, 5 Na-ascorbate, 3 Na-pyruvate, 2 CaCl_2_·4H_2_O and 2 MgSO_4_·7H_2_O continuously bubbled with 95% O_2_/5% CO_2_. Slices were held in this chamber for use in acute recordings or transferred to a 6-well plate for long-term culture and viral transduction. Cultured slices were placed on membrane inserts and wells were filled with culture medium consisting of 8.4 g/L MEM Eagle medium, 20% heat-inactivated horse serum, 30 mM HEPES, 13 mM D-glucose, 15 mM NaHCO_3_, 1 mM ascorbic acid, 2 mM MgSO_4_·7H_2_O, 1 mM CaCl_2_.4H_2_O, 0.5 mM GlutaMAX-I, and 1 mg/L insulin (Ting et al 2018). The slice culture medium was carefully adjusted to pH 7.2-7.3, osmolality of 300-310 mOsmoles/Kg by addition of pure H_2_O, sterile-filtered and stored at 4°C for up to two weeks.

Culture plates were placed in a humidified 5% CO_2_ incubator at 35°C and the slice culture medium was replaced every 2-3 days until end point analysis. 1-3 hours after brain slices were plated on cell culture inserts, brain slices were infected by direct application of concentrated AAV viral particles over the slice surface (Ting et al 2018).

Thy1 and Etv1 mice were deeply anesthetized by IP administration of ketamine (130 mg/kg) and xylazine (8.8 mg/kg) mix and were perfused through the heart with chilled (2-4°C) sodium-free aCSF consisting of (in mM): 210 Sucrose, 7 D-glucose, 25 NaHCO_3,_ 2.5 KCl, 1.25 NaH_2_PO_4_, 7 MgCl_2_, 0.5 CaCl_2,_1.3 Na-ascorbate, 3 Na-pyruvate bubbled with carbogen (95% O_2_/5% CO_2_). Near coronal slices 300 microns thick were generated using a Leica vibratome (VT1200) in the same sodium-free aCSF and were transferred to warmed (35°C) holding solution (in mM): 125 NaCl, 2.5 KCl, 1.25 NaH_2_PO_4_, 26 NaHCO_3_, 2 CaCl_2_, 2 MgCl_2_, 17 dextrose, and 1.3 sodium pyruvate bubbled with carbogen (95% O_2_/5% CO_2_). After 30 minutes of recovery, the chamber holding slices was allowed to cool to room temperature.

#### Patch clamp electrophysiology

Macaque, human and *Pvalb*-TdTomato mouse brain slices were placed in a submerged, heated (32-34°C) recording chamber that was continually perfused (3-4 mL/min) with aCSF under constant carbogenation and containing (in mM) 1): 119 NaCl, 2.5 KCl, 1.25 NaH_2_PO_4_, 24 NaHCO_3_, 12.5 glucose, 2 CaCl_2_·4H_2_O and 2 MgSO_4_·7H_2_O (pH 7.3-7.4). Slices were viewed with an Olympus BX51WI microscope and infrared differential interference contrast (IR-DIC) optics and a 40x water immersion objective. The infragranular layers of macaque primary motor cortex and human premotor cortex are heavily myelinated, which makes visualization of neurons under IR-DIC virtually impossible. To overcome this challenge, we labeled neurons using various viral constructs in organotypic slice cultures (Extended Data Fig. 10g).

Patch pipettes (2-6 MΩ) were filled with an internal solution containing (in mM): 110.0 K-gluconate, 10.0 HEPES, 0.2 EGTA, 4 KCl, 0.3 Na2-GTP, 10 phosphocreatine disodium salt hydrate, 1 Mg-ATP, 20 µg/ml glycogen, 0.5U/µL RNAse inhibitor (Takara, 2313A) and 0.5% biocytin (Sigma B4261), pH 7.3. Fluorescently labeled neurons from *Thy1* or *Etv1* mice were visualized through a 40x objective using either Dodt contrast with a CCD camera (Hamamatsu) and/or a 2-photon imaging/ uncaging system from Prairie (Bruker) Technologies. Recordings were made in aCSF: (in mM): 125 NaCl, 3.0 KCl, 1.25 NaH_2_PO_4_, 26 NaHCO_3_, 2 CaCl_2_, 1 MgCl2, 17 dextrose, and 1.3 sodium pyruvate bubbled with carbogen (95% O_2_/5% CO_2_) at 32-35°, with synaptic inhibition blocked using 100 µM picrotoxin.

Sylgard-coated patch pipettes (3-6 MΩ) were filled with an internal solution containing (in mM): 135 K-gluconate, 12 KCl, 11 HEPES, 4 MgATP, 0.3 NaGTP, 7 K_2_-phosphocreatine, 4 Na_2_-phophocreatine (pH 7.42 with KOH) with neurobiotin (0.1-0.2%), Alexa 594 (40 µM) and Oregon Green BAPTA 6F (100 µM).

Whole cell somatic recordings were acquired using either a Multiclamp 700B amplifier, or an AxoClamp 2B amplifier (Molecular Devices) and were digitized using an ITC-18 (HEKA). Data acquisition software was either MIES (https://github.com/AllenInstitute/MIES/) or custom software written in Igor Pro. Electrical signals were digitized at 20-50 kHz and filtered at 2-10 kHz. Upon attaining whole-cell current clamp mode, the pipette capacitance was compensated and the bridge was balanced. Access resistance was monitored throughout the recording and was 8-25 MΩ.

### Data analysis

Data were analyzed using custom analysis software written in Igor Pro. All measurements were made at resting membrane potential. Input resistance (R_N_) was measured from a series of 1 s hyperpolarizing steps from −150 pA to +50 pA in +20 pA increments. For neurons with low input resistance (e.g. the Betz cells) this current injection series was scaled by upwards of 4x. Input resistance (R_N_) was calculated from the linear portion of the current−steady state voltage relationship generated in response to these current injections. Resonance (*f*_R_) was determined from the voltage response to a constant amplitude sinusoidal current injection (Chirp stimulus). The chirp stimulus increased in frequency either linearly from 1-20 Hz over 20 s or logarithmically from 0.2-40 Hz over 20s. The amplitude of the Chirp was adjusted in each cell to produce a peak-to-peak voltage deflection of ∼10 mV. The impedance amplitude profile (ZAP) was constructed from the ratio of the fast Fourier transform of the voltage response to the fast Fourier transform of the current injection. ZAPs were produced by averaging at least three presentations of the Chirp and were smoothed using a running median smoothing function. The frequency corresponding to the peak impedance (Z_max_) was defined as the resonant frequency. Spike input/output curves were constructed in response to 1 s step current injections (50 pA-500 pA in 50 pA steps). For a subset of experiments, this current injection series was extended to 3A in 600 pA steps to probe the full dynamic range of low R_N_ neurons. Spike frequency acceleration analysis was performed for current injections producing ∼10 spikes during the 1 s step.

Acceleration ratio was defined as the ratio of the second to the last interspike interval. To examine the dynamics of spike timing over longer periods, we also measured spiking in response to 10 s step current injections in which the amplitude of the current was adjusted to produce ∼5 spikes in the first second. Action potential properties were measured for currents near rheobase. Action potential threshold was defined as the voltage at which the first derivative of the voltage response exceeded 20 V/s. AP width was measured at half the amplitude between threshold and the peak voltage. Fast AHP was defined relative to threshold. We clustered mouse, macaque and human pyramidal neurons into two broad groups based on their R_N_ and *f*_R_ using Ward’s algorithm.

### Biocytin histology

A horseradish peroxidase (HRP) enzyme reaction using diaminobenzidine (DAB) as the chromogen was used to visualize the filled cells after electrophysiological recording, and 4,6-diamidino-2-phenylindole (DAPI) stain was used identify cortical layers as described previously ^89^.

### Microscopy

Mounted sections were imaged as described previously ^89^. Briefly, operators captured images on an upright AxioImager Z2 microscope (Zeiss, Germany) equipped with an Axiocam 506 monochrome camera and 0.63x optivar. Two-dimensional tiled overview images were captured with a 20X objective lens (Zeiss Plan-NEOFLUAR 20X/0.5) in brightfield transmission and fluorescence channels. Tiled image stacks of individual cells were acquired at higher resolution in the transmission channel only for the purpose of automated and manual reconstruction. Light was transmitted using an oil-immersion condenser (1.4 NA). High-resolution stacks were captured with a 63X objective lens (Zeiss Plan-Apochromat 63x/1.4 Oil or Zeiss LD LCI Plan-Apochromat 63x/1.2 Imm Corr) at an interval of 0.28 µm (1.4 NA objective; mouse specimens) or 0.44 µm (1.2 NA objective; human and non-human primate specimens) along the Z axis. Tiled images were stitched in ZEN software and exported as single-plane TIFF files.

### Morphological reconstruction

Reconstructions of the dendrites and the full axon were generated based on a 3D image stack that was run through a Vaa3D-based image processing and reconstruction pipeline as described previously ^89^.

### Viral vector production and transduction

Recombinant AAV vectors were produced by triple-transfection of ITR-containing enhancer plasmids along with AAV helper and rep/cap plasmids using the AAV293 cell line, followed by harvest, purification and concentration of the viral particles. The AAV293 packaging cell line and plasmid supplying the helper function are available from a commercial source (Cell Biolabs). The PHP.eB capsid variant was generated by Dr. Viviana Gradinaru at the California Institute of Technology ^90^ and the DNA plasmid for AAV packaging is available from Addgene (plasmid#103005). Quality control of the packaged AAV was determined by viral titering to determine an adequate concentration was achieved (>5E^12^ viral genomes per mL), and by sequencing the AAV genome to confirm the identity of the viral vector that was packaged. Human and NHP L5 ET neurons including Betz cells were targeted in cultured slices by transducing the slices with viral vectors that either generically label neurons (AAV-hSyn1-tdTomato), or that enrich for L5 ET neurons by expressing reporter transgene under the control of the msCRE4 enhancer ^87^.

### Processing of Patch-seq samples

For a subset of experiments, the nucleus was extracted at the end of the recording and processed for RNA-sequencing. Prior to data collection for these experiments, all surfaces were thoroughly cleaned with RNAse Zap. The contents of the pipette were expelled into a PCR tube containing lysis buffer (Takara, 634894). cDNA libraries were produced using the SMART-Seq v4 Ultra Low Input RNA Kit for Sequencing according to the manufacturer’s instructions. We performed reverse transcription and cDNA amplification for X PCR cycles. Sample proceeded through Nextera NT DNA Library Preparation using Nextera XT Index Kit V2 Set A(FC-131-2001).

### Mapping of samples to reference taxonomies

To identify which cell type a given patch-seq nuclei mapped to, we used our previously described nearest centroid classifier ^1^. Briefly, a centroid classifier was constructed for Glutamatergic reference data (human SSv4 or macaque Cv3) using marker genes for each cluster. Patch-seq nuclei were then mapped to the appropriate species reference 100 times, using 80% of randomly sampled marker genes during each iteration. Probabilities for each nuclei mapping to each cluster were computed over the 100 iterations, resulting in a confidence score ranging from 0 to 100. We identified four human patch-seq nuclei that mapped with > 85% confidence and four macaque nuclei that mapped with > 93% confidence to a cluster in the L5 ET subclass.

### Data availability

Raw sequence data are available for download from the Neuroscience Multi-omics Archive (https://nemoarchive.org/) and the Brain Cell Data Center (https://biccn.org/data). Visualization and analysis tools are available at NeMO Analytics (Individual species: https://nemoanalytics.org//index.html?layout_id=ac9863bf; Integrated species: https://nemoanalytics.org//index.html?layout_id=34603c2b) and Cytosplore Viewer (https://viewer.cytosplore.org/). These tools allow users to compare cross-species datasets and consensus clusters via genome and cell browsers and calculate differential expression within and among species. A semantic representation of the cell types defined through these studies is available in the provisional Cell Ontology (https://bioportal.bioontology.org/ontologies/PCL; Supplementary Table 1).

### Code availability

Code to reproduce figures will be available for download from https://github.com/AllenInstitute/BICCN_M1_Evo.

## Acknowledgements

We thank the Tissue Procurement, Tissue Processing and Facilities teams at the Allen Institute for Brain Science for assistance with the transport and processing of postmortem and neurosurgical brain specimens; the Technology team at the Allen Institute for assistance with data management; M. Vawter, J. Davis and the San Diego Medical Examiner’s Office for assistance with postmortem tissue donations. We thank Ximena Opitz-Araya and Allen Institute for Brain Science Viral Technology team for AAV packaging. We thank Krissy Brouner, Augustin Ruiz, Tom Egdorf, Amanda Gary, Michelle Maxwell, Alice Pom and Jasmine Bomben for biocytin staining. We thank Nadezhda Dotson, Rachel Enstrom, Madie Hupp, Lydia Potekhina, and Shea Ransford for imaging biocytin filled cells. We thank Lindsay Ng, Dijon Hill and Ram Rajanbabu for patching the human and mouse cells in the figure describing chandelier neurons and Sara Kebede, Alice Mukora, Grace Willams for reconstructing them. This work was funded by the Allen Institute for Brain Science and by the U.S. National Institutes of Health grant U01 MH114812-02 to E.S.L. Support for the development of NS-Forest v.2 and the provisional cell ontology was provided by the Chan–Zuckerberg Initiative DAF, an advised fund of the Silicon Valley Community Foundation (2018-182730). G.Q. is supported by NSF CAREER award 1846559. This work was partially supported by an NWO Gravitation project: BRAINSCAPES: A Roadmap from Neurogenetics to Neurobiology (NWO: 024.004.012) and NWO TTW project 3DOMICS (NWO: 17126). This project was supported in part by NIH grants P51OD010425 from the Office of Research Infrastructure Programs (ORIP) and UL1TR000423 from the National Center for Advancing Translational Sciences (NCATS). Its contents are solely the responsibility of the authors and do not necessarily represent the official view of NIH, ORIP, NCATS, the Institute of Translational Health Sciences or the University of Washington National Primate Research Center. This work is supported in part by NIH BRAIN Initiative award RF1MH114126 from the National Institute of Mental Health to E.S.L., J.T.T., and B.P.L., NIH BRAIN Initiative award U19MH121282 to J.R.E., the National Institute on Drug Abuse award R01DA036909 to B.T., National Institute of Neurological Disorders and Stroke award R01NS044163 to W.J.S. and the California Institute for Regenerative Medicine (GC1R-06673-B) and the Chan Zuckerberg Initiative DAF, an advised fund of the Silicon Valley Community Foundation (2018–182730) to R.H.S. J.R.E. is an Investigator of the Howard Hughes Medical Institute. The authors thank the Allen Institute founder, Paul G. Allen, for his vision, encouragement and support.

## Author contributions

RNA data generation: A.M.Y., A.Re., A.T., B.B.L., B.T., C.D.K., C.R., C.R.P., C.S.L., D.B., D.D., D.M., E.S.L., E.Z.M., F.M.K., G.F., H.T., H.Z., J.C., J.Go., J.S., K.C., K.L., K.Si., K.Sm., K.Z., M.G., M.K., M.T., N.Dee, N.M.R., N.P., R.D.H., S.A.M., S.D., S.L., T.C., T.E.B., T.P., W.J.R. mC data generation: A.B., A.I.A., A.Ri., C.L., H.L., J.D.L., J.K.O., J.R.E., J.R.N., M.M.B., R.G.C. ATAC data generation: A.E.S., B.B.L., B.R., B.T., C.R.P., C.S.L., D.D., J.C., J.D.L., J.K.O., K.Z., L.T.G., M.M.B., N.P., S.P., W.J.R., X.H., X.W. Electrophysiology, morphology, and Patch-seq data generation: A.L.K., B.E.K., D.M., E.S.L., G.D.H., J.Go., J.T.T., K.Sm., M.T., N.Dem., P.R.N., R.D., S.A.S., S.O., T.L.D., T.P., W.J.S. Data archive and infrastructure: A.E.S., A.M., B.R.H., H.C.B., J.A.M., J.Go., J.K., J.O., M.M., O.R.W., R.H., S.A.A., S.S., Z.Y. Cytosplore Viewer software: B.P.L., B.V.L., J.E., T.H. Data analysis: A.D., B.B.L., B.D.A., B.E.K., B.P.L., B.V.L., D.D., E.A.M., E.S.L., F.M.K., F.X., H.L., J.E., J.Gi., J.Go., J.R.E., J.T.T., K.Sm., M.C., N.Dem., N.L.J., O.P., P.V.K., Q.H., R.F., R.H.S., R.Z., S.F., S.O., T.E.B., T.H., W.D., W.T., Y.E.L., Z.Y. Data interpretation: A.D., A.Re., B.B.L., B.E.K., B.T., C.K., C.L., E.S.L., F.X., H.L., H.Z., J.Gi., J.Go., J.R.E., J.T.T., M.C., M.H., M.M.B., N.Dem., N.L.J., P.R.H., P.V.K., Q.H., R.D.H., R.H.S., R.Z., S.D., S.O., T.E.B., W.T., Y.E.L., Z.Y. Writing manuscript: A.D., B.B.L., B.E.K., C.K., E.S.L., F.M.K., M.C., N.Dem., N.L.J., P.R.H., Q.H., R.H.S., T.E.B., W.J.S., W.T.

## Materials & Correspondence

Correspondence and requests for materials should be addressed to E.S.L. and T.E.B.

## Supplementary Table legends

**Supplementary Table 1.** Provisional cell ontology (pCL) terms for human, mouse, and marmoset primary motor cortex cell types. Column headers are described as follows: pCL_id is a unique alphanumeric identifier assigned to each provisional cell type. CL_id is the Cell Ontology (CL) identifier for those parent cell type classes already represented in CL. pCL_name and Transcriptome data cluster are labels given according to each species naming convention that combines information about cortical layer enrichment and genes expressed in data cluster transcriptomes. TDC_id is a unique identifier assigned to the transcriptome data cluster. The part_of (uberon_id) and part_of (uberon_name) columns contain unique identifiers and names for tissue anatomic regions from which the experiment specimen was derived, in this case primary motor cortex. The is_a (CL or pCL_id) and is_a (CL or pCL_name) columns contain parent cell type or provisional cell type identifiers and names, respectively. Cluster_size indicates the number of single-nucleus or cell transcriptomes that were assigned membership to the transcriptome data cluster. Marker_gene_evidence indicates the number of marker genes that are necessary and sufficient to define the transcriptome cell type data cluster with maximal classification accuracy based on the NS-Forest v2.1 algorithm (see Supplementary Tables 4-6). F-measure_evidence is the f-beta score of classification accuracy from the NS-Forest v2.1 algorithm using the marker genes listed. The selectively_expresses column lists the minimum set of marker genes necessary and sufficient to define the transcriptome cell type data cluster. The definition brings together features to form a data driven ontological representation for each cell type cluster. The pCL annotations are available at https://github.com/mkeshk2018/Provisional_Cell_Ontology and https://bioportal.bioontology.org/ontologies/PCL.

**Supplementary Table 2.** Cluster annotations for human, marmoset, and mouse in separate worksheets. Cluster_label column identifies the RNA-seq cluster within each species. Cluster_size column denotes the number of nuclei that reside within each cluster (cluster_label). Class column identifies which cell class each cluster belongs to. Subclass column identifies which cell subclass each cluster belongs to. Cross-species cluster column indicates the cross-species consensus cluster taxonomy. DNAm_cluster_label column identifies the transcriptomic cluster (cluster_label) that is aligned to DNAm-determined clusters. ATAC_cluster label column identifies the transcriptomic cluster (cluster_label) that is aligned to ATAC-determined clusters.

**Supplementary Table 3.** Application of Allen Institute nomenclature schema to mouse, marmoset, and human M1 taxonomies. The “taxonomy_ids” tab lists ids and descriptions for the 11 taxonomies included and which tab those taxonomies are shown on. The “preferred_aliases” tab shows a list of preferred aliases for linking between taxonomies, as well as descriptions for these. The next five tabs show nomenclatures for each of the taxonomies and have the following column headers: “tree_order” is the order shown in the tree (if any); “cell_set_alias”, “cell_set_label”, and “cell_set_accession” are unique identifiers, as described in the Allen Institute nomenclature page (https://portal.brain-map.org/explore/classes/nomenclature), with “cell_set_alias” including the names used in this manuscript; “cell_set_preferred_alias” indicates which clusters correspond to the “preferred_alias”es from the previous tab, if any; “cell_set_alias_integrated” shows linkages between single species transcriptomics taxonomies and the integrated taxonomy; “cell_set_labels_CS191213#“ columns indicate linkages between cell sets in the transcriptomics and other modalities within a single species; “cell_set_descriptor” shows the type of cell set (or level of ontology); and “taxonomy_id” links to the “taxonomy_id” tab. Finally, the “Cell class hierarchy” tab shows the ordered class, level2, and subclass hierarchy and associated colors used as cell sets in previous tabs.

**Supplementary Table 4.** NS-Forest v2.1 was used to determine cell type cluster marker genes for all annotated levels of the human primary motor cortex cell type taxonomy defined by RNA-seq (Cv3). “clusterName” corresponds to the annotation label, either a cell type cluster name or a parent cell type class in the taxonomy. “markerCount” gives the optimal number of marker genes in the set that best discriminates the label. The “f-measure” column gives the f-beta score for classification using the set of markers. The next four columns “True Negative”, “False Positive”, “False Negative”, “True Positive” give the confusion matrix for the label given the set of markers. Finally, “Marker 1-5” lists the gene symbols corresponding to the optimal set of markers.

**Supplementary Table 5.** NS-Forest v2.1 was used to determine cell type cluster marker genes for all annotated levels of the mouse primary motor cortex cell type taxonomy defined by RNA-seq (Cv3). “clusterName” corresponds to the annotation label, either a cell type cluster name or a parent cell type class in the taxonomy. “markerCount” gives the optimal number of marker genes in the set that best discriminates the label. The “f-measure” column gives the f-beta score for classification using the set of markers. The next four columns “True Negative”, “False Positive”, “False Negative”, “True Positive” give the confusion matrix for the label given the set of markers. Finally, “Marker 1-5” lists the gene symbols corresponding to the optimal set of markers.

**Supplementary Table 6.** NS-Forest v2.1 was used to determine cell type cluster marker genes for all annotated levels of the marmoset primary motor cortex cell type taxonomy defined by RNA-seq (Cv3). “clusterName” corresponds to the annotation label, either a cell type cluster name or a parent cell type class in the taxonomy. “markerCount” gives the optimal number of marker genes in the set that best discriminates the label. The “f-measure” column gives the f-beta score for classification using the set of markers. The next four columns “True Negative”, “False Positive”, “False Negative”, “True Positive” give the confusion matrix for the label given the set of markers. Finally, “Marker 1-5” lists the gene symbols corresponding to the optimal set of markers.

**Supplementary Table 7.** DEGs determined by ROC test between each GABAergic neuron subclass and all other GABAergic nuclei within each species. Columns are labeled myAUC, which contains AUC scores > 0.7; avg_diff, which contains difference in expression between target subclass and all other GABAergic neurons; power; pct.1, which indicates the percent of nuclei that express the gene in the target cluster; pct.2, which indicates the percent of non-target nuclei that express the gene; cluster, which denotes the target cluster; gene, indicating the gene that was identified as DE; and species, which indicates the species the test was performed in.

**Supplementary Table 8.** List of DEGs (from Supplementary Table 7) that is sorted according to the order the genes appear within the heatmap.

**Supplementary Table 9.** Supervised MetaNeighbor results, within- and across-species. Each row corresponds to a unique entry for a given gene set and a given cell class, either Glutamatergic or GABAergic. The first five columns provide information about the gene sets, namely their provenance (SynGO or HGNC); numerical IDs; descriptive labels; manual classifications for plotting and interpretation; and finally the number of genes included in the analysis (after subsetting to genes with 1-1 orthologs across all three species). The sixth column indicates cell class. The remaining columns contain MetaNeighbor AUROCs for various analyses: within_species_meanROC (column 7) provides the mean of within-mouse (column 8), within-marmoset (column 9) and within-human (column 10) performance. For each species, tests were run with random 3-fold cross-validation, and the average across folds is reported. Columns 11 and 12 contain results from cross-species analyses, detailed in the methods. Results are sorted by their AUROC across primates (column 12).

**Supplementary Table 10.** DEGs determined by ROC test between each glutamatergic neuron subclass and all other glutamatergic nuclei within each species. Columns are labeled myAUC, which contains AUC scores > 0.7; avg_diff, which contains difference in expression between target subclass and all other glutamatergic neurons; power; pct.1, which indicates the percent of nuclei that express the gene in the target cluster; pct.2, which indicates the percent of non-target nuclei that express the gene; cluster, which denotes the target cluster; gene, indicating the gene that was identified as DE; and species, which indicates the species the test was performed in.

**Supplementary Table 11.** List of DEGs (from Supplementary Table 10) that is sorted according to the order the genes appear within the heatmap.

**Supplementary Table 12.** Average expression of isoforms in human and mouse subclasses and estimates of isoform genic proportions (P) based on the ratio of isoform to gene expression. Isoforms were included if they had at least moderate expression (TPM > 10) and P > 0.2 in either human or mouse and at least moderate gene expression (TPM > 10) in both species.

**Supplementary Table 13.** SNARE-Seq2 metadata, cluster annotations and quality statistics. Tab 14a indicates SNARE-Seq2 experiment level metadata (experiment name, library, patient, species, purification, age, sex) and mapping statistics for RNA (mean UMI detected, mean genes detected) and AC (mean fraction of reads in promoters or FRiP, mean uniquely mapped fragments grouped by 5000 base pair chromosomal bins, mean unique fragment counts per final peak locations, total number of final nuclei). Tab 14b indicates the SNARE-Seq2 local RNA clusters for human M1 generated using Pagoda2 (local cluster, annotated cluster name, broad cell type and abbreviation, k value used for Pagoda2 clustering, broad cell type markers, level 1 and level2 classes and associated markers, unique cluster markers). Tabs 14c-d indicates SNARE-Seq2 consensus or harmonized RNA and AC-Level cluster annotations for human and marmoset M1, respectively, including annotated cluster name, cluster order, associated subclass and class, and the number of datasets making up the clusters. Tabs 14e-f lists all metadata outlined in tabs 14a-d for all SNARE-Seq2 cell barcodes from human and marmoset M1 samples, respectively.

**Supplementary Table 14.** SNARE-Seq2 differentially accessible regions for human and marmoset M1. Tabs 15a and 15b show SNARE-Seq2 differentially accessible regions (DARs, q value < 0.001, log-fold change > 1) identified by AC-Level clusters (15a) or subclass level (15b) for human M1, indicating for each chromosomal location the p value (hypergeometric test), q value (Benjamini-Hochberg adjusted p value), log-fold change and associated cluster or subclass. Tab 15b shows subclass DARs (q value < 0.001, log-fold change > 1) for marmoset subclasses as in tab 15b. Tab 15d shows a summarization of human and marmoset DARs detected by matched subclasses, indicating actual number of DARs detected (tabs 15b and 15c) and the values normalized to cluster size and total number of DARs detected per species.

**Supplementary Table 15.** Cis-co-accessible sites, TF motif enrichments and differential TFBS activities for human and marmoset M1. Tab 16a (human M1) and 16b (marmoset M1) show cis-coaccessible network (CCAN) sites for subclass distinct markers genes (Wilcoxon Rank Sum test, adjusted P < 0.05, average log-fold change > 0.5). pct.1 indicates the percent of nuclei that express the gene in the target cluster, pct.2 indicates the percent of non-target nuclei that express the gene. For each cluster and marker gene, corresponding motif enrichment values (hypergeometric test) for gene-associated CCAN sites are shown (“observed” indicates number of features containing the motif, “background” indicates the total number of features from a random selection of 40000 features that contain the motif), and the motif associated differential chromVAR activity values identified using logistic regression. The full list of chromVAR differentially active TFBS activities are also provided. Tab 16c summarizes the number of CCAN-associated marker genes, associated TFBSs enriched and or active by subclass for both human and marmoset M1. Tabs 16d and 16e show cis-co-accessible sites, TFBS enrichments and differential activities by AC-level clusters for human and marmoset M1, respectively, similar to that provided in tabs 16a and 16b. Tab 16f shows chromVAR differentially active TFBS activities by consensus or harmonized cluster using logistic regression. Tabs 16g, 16h, and 16i show cis-co-accessible sites, TF motif enrichments and differential TFBS activities for human, marmoset and mouse M1 ChCs compared against BCs.

**Supplementary Table 16.** snmC-seq2 metadata. The table shows experiment level metadata, including species, sample name, gender, purification information, experiment nuclei numbers and pass-QC nuclei numbers.

**Supplementary Table 17.** Subclass TFBS enrichment results. TFBS enrichment analysis was done with AME ^78^ using JASPAR2020 motifs. Within a species, hypo-methylated DMRs in each subclass were tested against hypo-methylated DMRs of all the other subclasses (background). DMRs and 250bp around regions were used in the analysis. This table includes p-values and effect sizes (log2(TP/FP)) of the analysis results.

**Supplementary Table 18.** Subclass TFBS enrichment at TF cluster level. TFs in SI Tab 18 were grouped using clusters defined in Ref ^42^. The table lists the most significant p-values and the largest effect size of each TF cluster group.

**Supplementary Table 19.** DEGs determined by ROC test between chandelier cells and basket cells within each species. Columns are labeled as species, with true/false values indicating if a gene was enriched in chandelier cells for that species.

**Supplementary Table 20.** DEGs determined by ROC test between L5 ET subclass and L5 IT subclass within each species. Columns are labeled as species, with values of 1 indicating a gene was enriched in the L5 ET subclass for that species. A value of 0 indicates that the gene was not enriched in the L5 ET subclass for that species.

**Supplementary Table 21.** Genes with expression enrichment in L5 ET versus L5 IT that decreases with evolutionary distance from human (human > macaque > marmoset > mouse). Columns are labeled by species, and values indicate the log-fold change between L5 ET and L5 IT for that species. Genes were included if they had a minimum log-fold change equal to 0.5 in human.

---

**Extended Data Figure 1.**
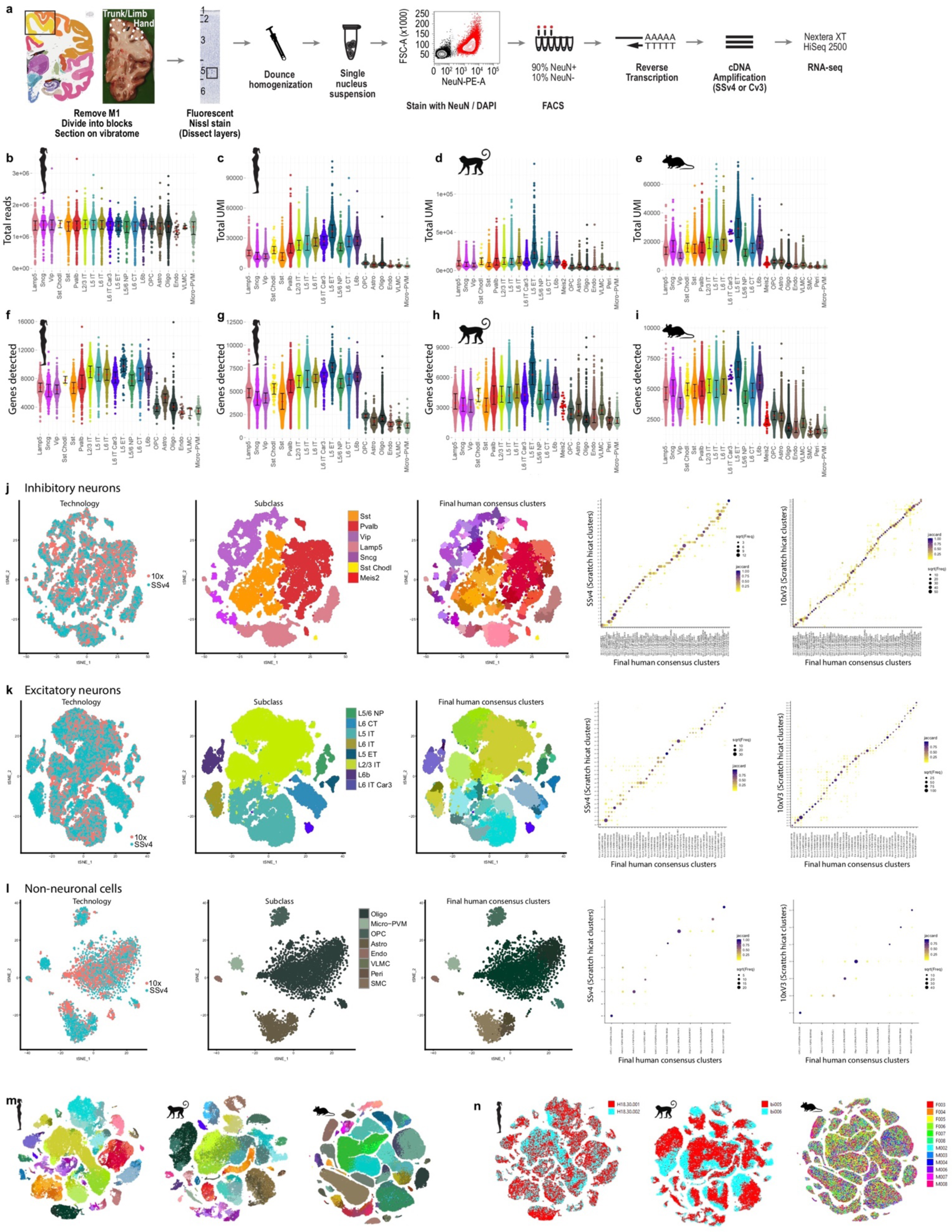
RNA-seq quality metrics and integration of human datasets. **a**, Schematic of single-nucleus isolation from M1 of post-mortem human brain and profiling with RNA-seq. Box in the Nissl image highlights a cluster of Betz cells in L5. **b**, Using SSv4, > 1 million total reads were sequenced across all subclasses in human. **c-e**, Using Cv3, total unique molecular identifiers (UMI) varies between subclasses, and these differences are shared across species. **f-i**, Gene detection (expression > 0) is highest in human using SSv4 (**e**) and lowest for marmoset using Cv3 (**h**). Note that the average read depth used for SSv4 was approximately 20-fold greater than for Cv3 (target 60,000 reads per nucleus). **j-k**, tSNE projections of single nuclei based on expression of several thousand genes with variable gene expression and colored by cluster label (**j**) or donor (**k**). **l-n**, Integration of SSv4 and Cv3 RNA-seq datasets from human single nuclei isolated from GABAergic (**l**) and glutamatergic (**m**) neurons and non-neuronal cells (**n**). Left: UMAP visualizations colored by RNA-seq technology, cell subclass, and unsupervised consensus clusters. Right: Confusion matrices show membership of SSv4 and Cv3 nuclei within integrated consensus clusters.

**Extended Data Figure 2.**
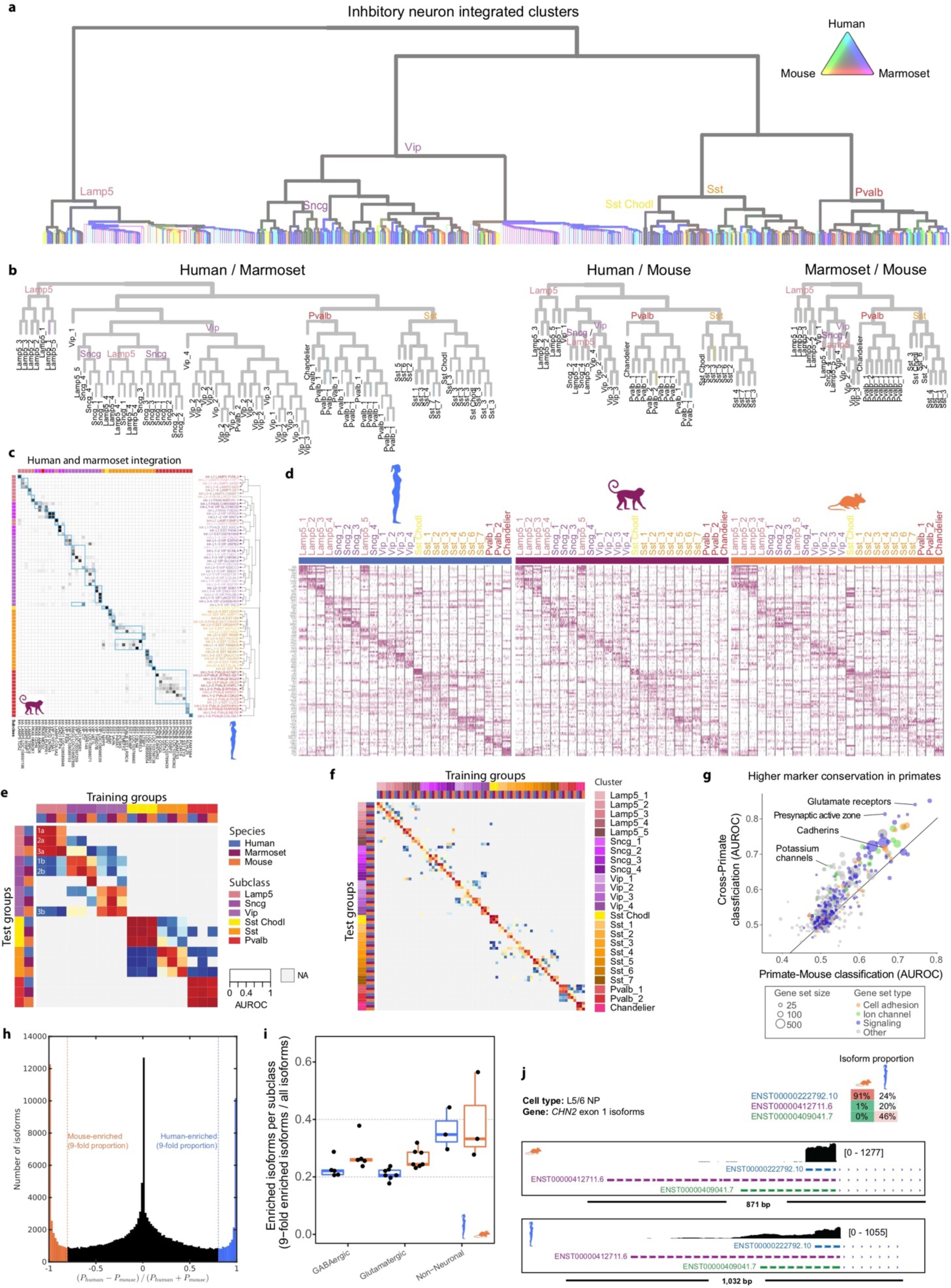
RNA-seq integration of GABAergic neurons across species. **a**, Dendrogram of GABAergic neuron clusters from unsupervised clustering of integrated RNA-seq data from human, marmoset and mouse. Edge thickness indicates the relative number of nuclei, and edge color indicates species mixing (grey is well mixed). Major branches are labeled by subclass. Dendrogram shown in **Figure 2f** is derived from this tree based on pruning species-specific branches. **b**, Dendrograms of pairwise species integrations from **Figure 2g** with leaves labeled by cross-species clusters and edges colored by species mixing. **c**, Cluster overlap heatmap from human-marmoset pairwise Seurat integration showing the proportion of within-species clusters that coalesce within integrated clusters. Columns and rows are ordered as in **Figure 2e** with cross-species consensus clusters indicated by blue boxes. Top and left color bars indicate subclasses of within-species clusters. **d**, Heatmaps showing scaled expression of the top 5 marker genes for each GABAergic cross-species cluster, and 5 marker genes for *Lamp5* and *Sst*. Initial genes were identified by performing a Wilcox test of every integrated cluster against every other GABAergic nuclei. Additional DEGs were identified for *Lamp5* and *Sst* cross-species clusters, by comparing one of the cross-species clusters to all other related nuclei (e.g. *Sst*_1 against all other *Sst*). **e-f**, Heatmap of 1-vs-best MetaNeighbor scores for GABAergic subclasses (**e**) and clusters (**f**). Each column shows the performance for a single training group across the three test datasets. AUROCs are computed between the two closest neighbors in the test dataset, where the closer neighbor will have the higher score, and all others are shown in gray (NA). For example, in **e** the first column contains results of training on human *Lamp5*, labeled with numbers to indicate test datasets, where 1 is human, 2 is marmoset and 3 is mouse, and letters to indicate closest (**a**) and second-closest (**b**) neighboring groups. Dark red 3×3 blocks along the diagonal indicate high transcriptomic similarity across all three species. **g**, Scatter plot of MetaNeighbor analysis showing the performance (AUROC) of gene sets to classify GABAergic neuron consensus types by training with human or marmoset data and testing with the other species (Cross-Primate, y-axis) or training with primate data and testing with mouse (Primate-Mouse, x-axis). Gene set size and type are indicated by point size and color, respectively. **h**, Histogram of the relative difference in isoform genic proportion (P) between human and mouse for all subclass comparisons. All moderately to highly expressed isoforms were included (gene TPM > 10 in both species; isoform TPM > 10 and proportion > 0.2 in either species). Vertical lines indicate >9-fold change in mouse or human. **i**, Proportion of all isoforms in **h** that switch between species (FDR P < 0.05; >9-fold change in P) summarized by subclass and grouped by cell class. **j**, Comparison between species of isoform genic proportions for the top three most common isoforms of Chimerin 2 (*CHN2*) expressed in the L5/6 NP subclass. Genome browser tracks of RNA-seq (SSv4) reads in human and mouse at the *CHN2* locus.

**Extended Data Figure 3.**
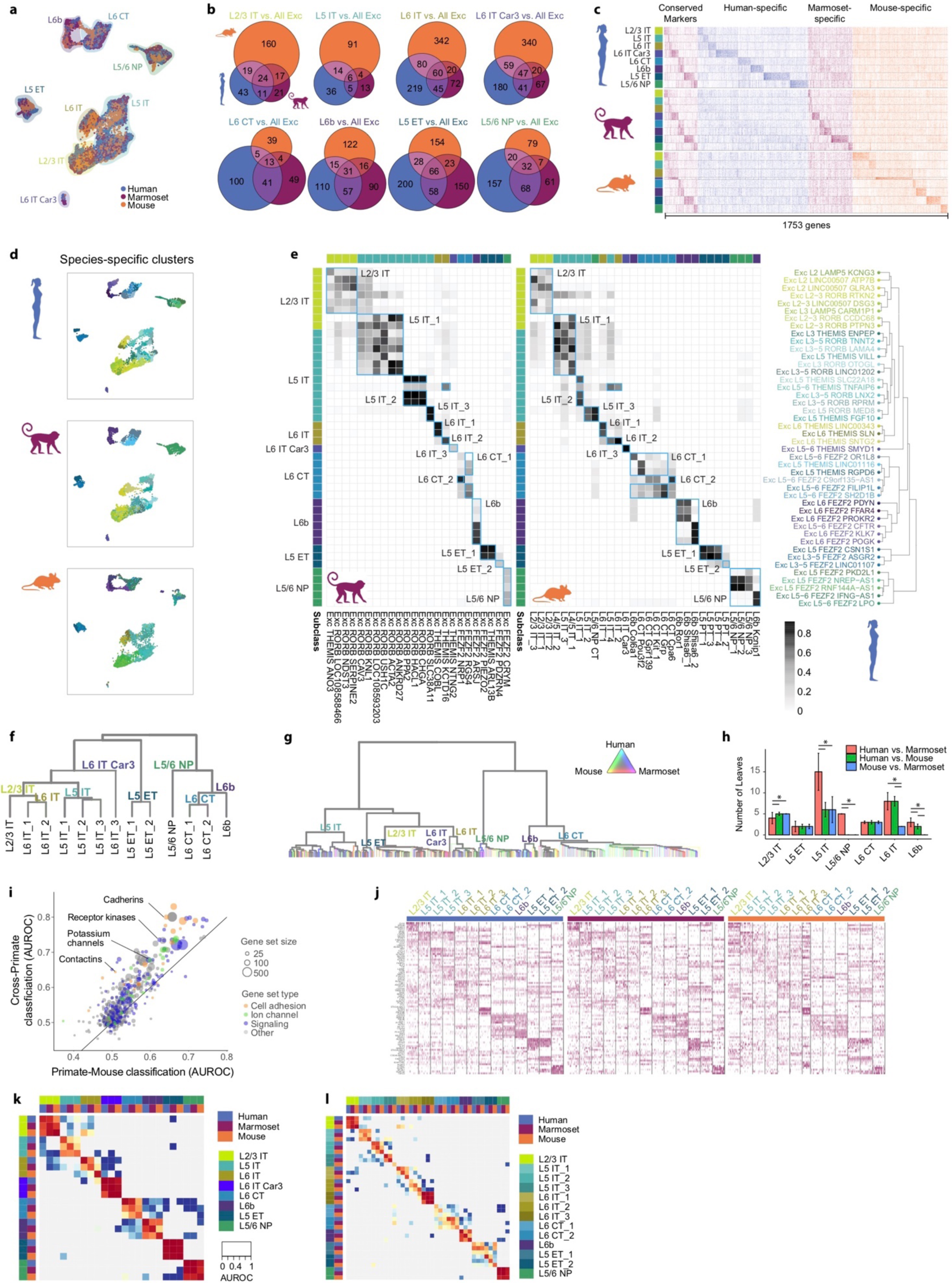
Glutamatergic neuron cell type homology across species. **a**, UMAP visualization of integrated snRNA-seq data from human, marmoset, and mouse glutamatergic neurons. Highlighted colors indicate subclass. **b**, Venn diagrams indicating number of shared DEGs across species by subclass. DEGs determined by ROC test of subclass against all other glutamatergic subclasses within a species. **c**, Heatmap of all DEGs from **b** ordered by subclass and species enrichment. Heatmap shows expression scaled by column for up to 50 randomly sampled nuclei from each subclass for each species. **d**, UMAP visualization of integrated snRNA-seq data with projected nuclei split by species. Colors indicate different within-species clusters. **e**, Cluster overlap heatmap showing the proportion of within-species clusters that coalesce with a given integrated cross-species cluster. Cross-species clusters are labelled and indicated by blue boxes with human-marmoset overlap shown to the left and human-mouse overlap shown to the right. Top and left axes indicate the subclass of a given within-species cluster by color. Bottom axis indicates marmoset (left) and mouse (right) within species clusters. Right axis shows the glutamatergic branch of the human dendrogram from **Figure 1c**. **f**, Dendrogram of glutamatergic neuron cross-species clusters. **g**, Unpruned dendrogram of glutamatergic neuron clusters from unsupervised clustering of integrated RNA-seq data. Edge thickness indicates the relative number of nuclei, and edge color indicates species mixing. Major branches are labeled by subclass. **h**, Bar plots quantifying the number of well-mixed clusters from unsupervised clustering of pairwise species integrations. Significant differences (adjusted P < 0.05, Tukey’s HSD test) between species are indicated for each subclass. **i**, Scatter plot of MetaNeighbor analysis showing the performance (AUROC) of gene sets to classify glutamatergic neuron consensus types by training with human or marmoset data and testing with the other species (Cross-Primate, y-axis) or training with primate data and testing with mouse (Primate-Mouse, x-axis). Gene set size and type are indicated by point size and color, respectively. **j**, Heatmaps showing scaled expression of marker genes for each glutamatergic cross-species cluster. The top 5 marker genes for each cross-species cluster are shown, with an additional 5 genes for L5 ET, L5 IT, and L6 IT. Initial genes were identified by performing a Wilcox test of every integrated cluster against every other glutamatergic nuclei. Additional DEGs were identified for L5 ET, L5 IT, and L6 IT cross-species clusters, by comparing one of the cross-species clusters to all other related nuclei (e.g. L5 IT_1 against all other L5 IT). **k**, **l**, Heatmap of 1-vs-best MetaNeighbor scores for glutamatergic subclasses (**k**) and clusters (**l**). Results are displayed as in **Extended Data** **Fig. 2e,f**.

**Extended Data Figure 4.**
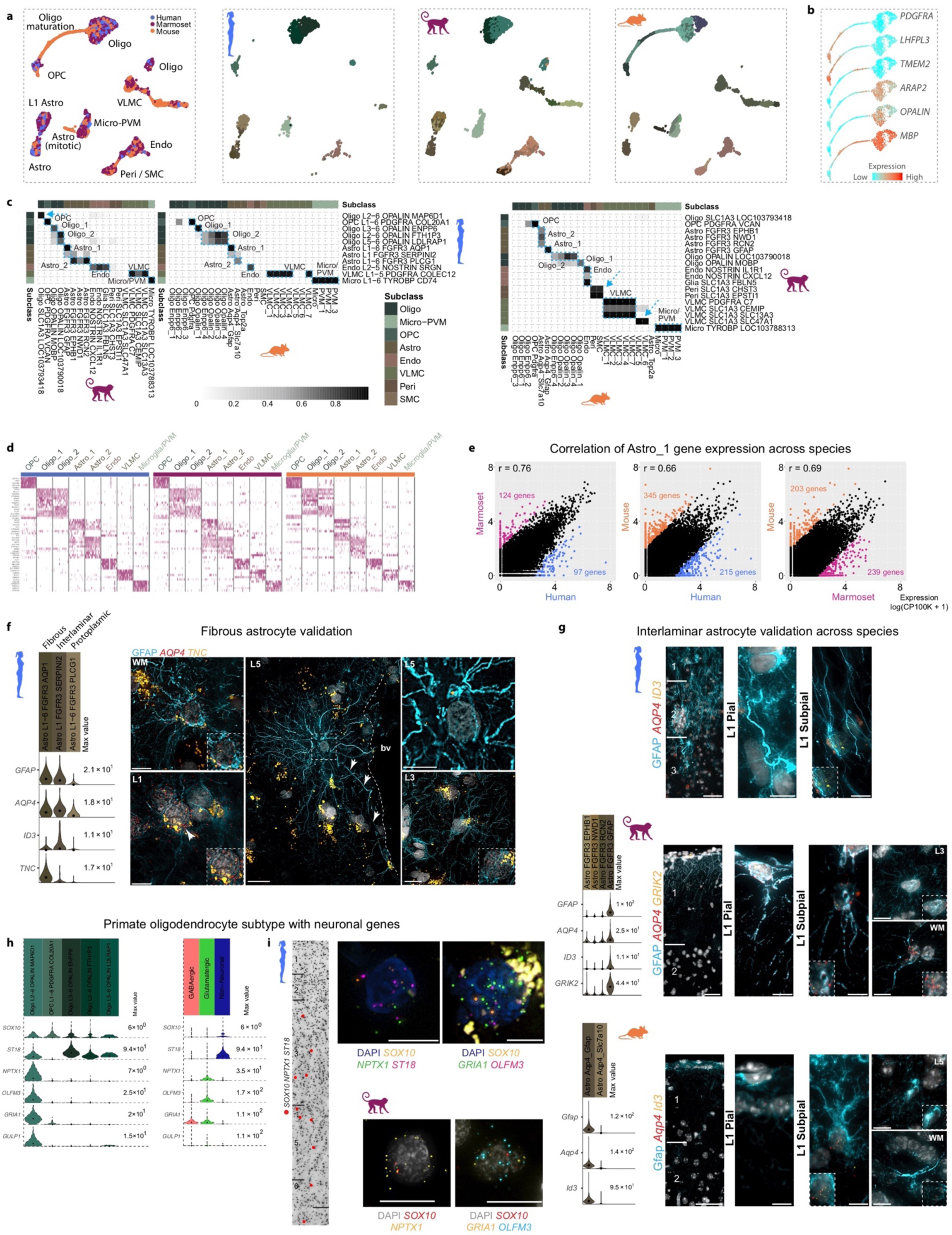
Non-neuronal cell type homology across species. **a**, UMAP plots of integrated RNA-seq data for non-neuronal nuclei, colored by species and within-species clusters. Note that some cell types are present in only one or two species. **b**, UMAP of mouse oligodendrocyte precursors and mature cells showing expression levels of marker genes for different stages of cell maturation. **c**, Heatmaps of the proportion of nuclei in each species-specific cluster that overlap in the integrated RNA-seq analysis. Blue boxes define homologous cell types that can be resolved across all three species. Arrows highlight clusters that overlap between two species and are not detected in the third species, due to differences in sampling depth of non-neuronal cells, relative abundances of cell types between species, or evolutionary divergence. **d**, Conserved marker genes for homologous cell types across species. **e**, Pairwise comparisons between species of log-transformed gene expression of the Astro_1 type. Colored points correspond to significantly differentially expressed (DE) genes (FDR < 0.01, log-fold change > 2). r, Spearman correlation. **f**, Fibrous astrocyte in situ validation. Violin plots of marker genes of human astrocyte clusters that correspond to fibrous, interlaminar, and protoplasmic types based on in situ labeling of types. Left ISH: Fibrous astrocytes located in the white matter (WM, top) and a subset of L1 (bottom) astrocytes express the Astro L1-6 *FGFR3 AQP1* marker gene *TNC*. Middle ISH: Image of putative varicose projection astrocyte located in cortical L5 adjacent to a blood vessel (bv) and extending long GFAP-labeled processes (white arrows) does not express the marker gene *TNC*. The white dashed box indicates the area shown at higher magnification in the top right panel. Likewise, the L3 protoplasmic astrocyte shown in the bottom right panel does not express *TNC*. **g**, Combined GFAP immunohistochemistry and RNAscope FISH for markers of L1 astrocytes in human, mouse, and marmoset. In human (top), pial and subpial interlaminar astrocytes are labeled with *AQP4* and *ID3* and extend long processes from L1 down to L3. In marmoset (middle), both pial and subpial L1 astrocytes express *AQP4* and *GRIK2* and extend GFAP-labeled processes through L1 that terminate before reaching L2. An image of a marmoset protoplasmic astrocyte located in L3 shows that this astrocyte type does not express the marker gene *GRIK2*. A subset of marmoset fibrous astrocytes located in the white matter (WM) express *GRIK2*, suggesting that fibrous and L1 astrocytes have a shared gene expression signature as shown in human ^2^. L1 astrocytes in mouse (bottom) consist of pial and subpial types that differ morphologically but are characterized by their expression of the genes *Aqp4* and *Id3*. Pial astrocytes in mouse extend short Gfap-labeled processes that terminate within L1 whereas mouse subpial astrocytes appear to extend processes predominantly toward the pial surface. Protoplasmic astrocytes (example shown in L5) do not express *Id3*, whereas fibrous astrocytes in mouse share expression of *Id3* with L1 astrocyte types. Inset images outlined with white dashed boxes illustrate cells in each of the accompanying images at higher magnification to show RNAscope spots for each gene labeled. Scale bars, 20 µm. **h**, Violin plots of marker genes of oligodendrocyte lineage clusters in human. Transcripts detected in the Oligo L2−6 *OPALIN MAP6D1* cluster include genes expressed almost exclusively in neuronal cells. Scale bars, 20 µm. **i**, Left: Inverted DAPI image showing a column of cortex labeled with markers of the human Oligo L2-6 *OPALIN MAP6D1* type. Red dots show cells triple labeled with *SOX10*, *NPTX1*, and *ST18*. Top right: Examples of cells labeled with marker gene combinations specific for the human Oligo L2-6 *OPALIN MAP6D1* type. Bottom right: Example of a marmoset cell labeled with the marker genes *OLIG2* and *NRXN3*. Scale bars, 20 µm.

**Extended Data Figure 5.**
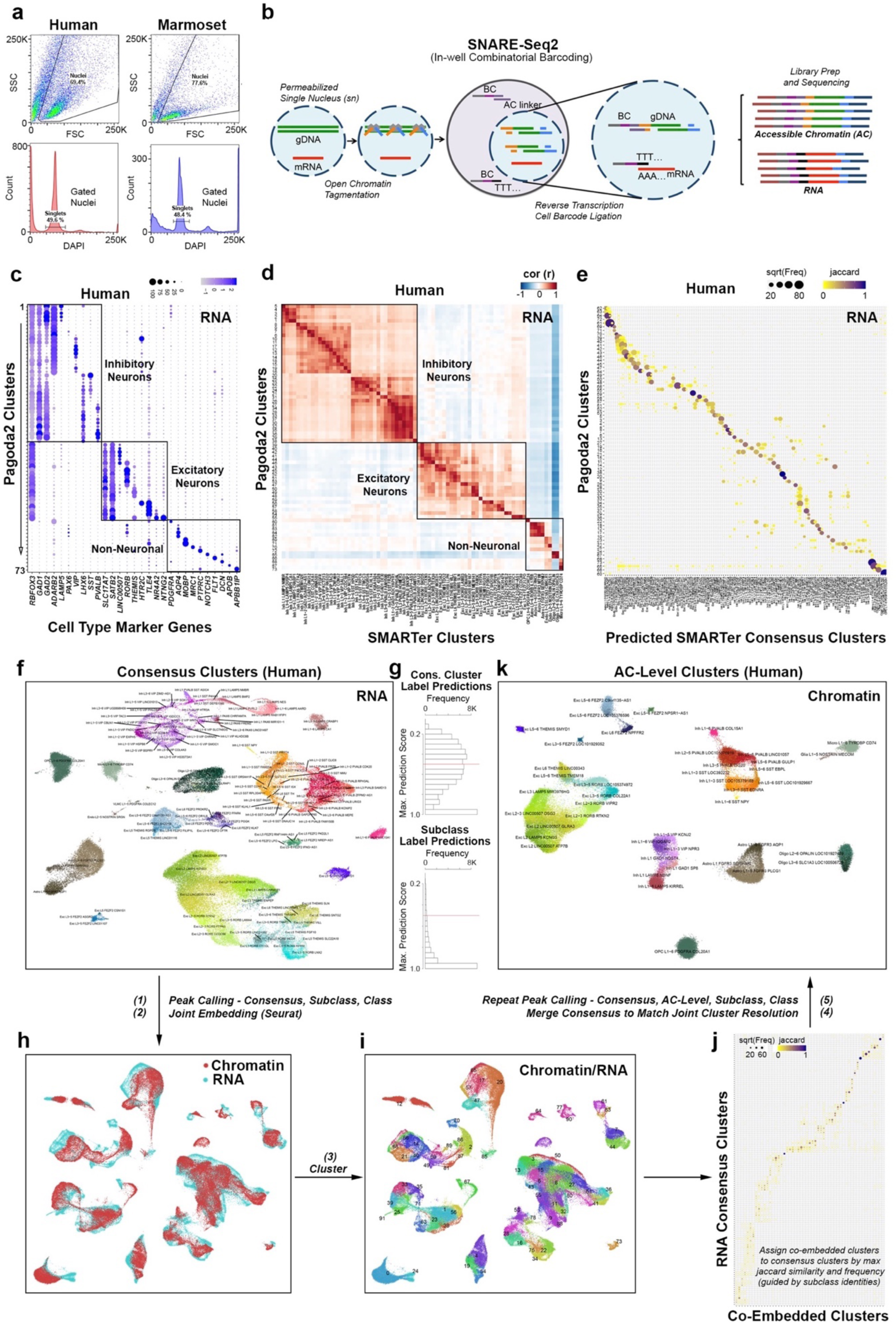
SNARE-seq2 transcriptomic profiling resolves M1 cell types. **a-b**, FACS gating parameters used for sorting human and marmoset single nuclei (**a**) that were used for SNARE-seq2 as outlined in (**b**), to generate both RNA and accessible chromatin (AC) libraries having the same cell barcodes. **c**, Dot plot showing averaged marker gene expression values (log scale) and proportion expressed for clusters identified in a preliminary analysis of SNARE-seq2 RNA using Pagoda2. **d**, Correlation heatmap of averaged scaled gene expression values for Pagoda2 clusters against SSv4 clusters from the same M1 region. **e**, Jaccard similarity plot for cell barcodes grouped according to Pagoda2 clustering compared against the predicted SSv4 consensus clustering. **f-k**, Overview of AC-level cluster assignment using RNA-defined clusters indicating the five main steps of the process. **f**, Consensus clusters visualized by UMAP on RNA expression data and that were used to independently call peaks from AC data. **g**, Histograms showing maximum prediction scores for consensus cluster (top) and subclass (bottom) labels from RNA data to corresponding accessibility data (cicero gene activities). **h**, Consensus cluster peaks, as well as those identified from subclass and class level barcode groupings, were combined and the corresponding peak by cell barcode matrix was used to predict gene activity scores using Cicero for integrative RNA/AC analyses. UMAP shows joint embedding of RNA and imputed AC expression values using Seurat/Signac. **i**, UMAP showing clusters identified from the joint embedding (**h**). **j**, Jaccard similarity plot comparing cell barcodes either grouped according to RNA consensus clustering or joint RNA/AC clustering (**i**). RNA consensus clusters were merged to best match the cluster resolution achieved from co-embedded clusters. Chromatin peak counts generated from peak calling independently on consensus, AC-level, subclass, and class barcode groupings were used to generate a final peak by cell barcode matrix. **k**, Final AC-level clusters visualized using UMAP.

**Extended Data Figure 6.**
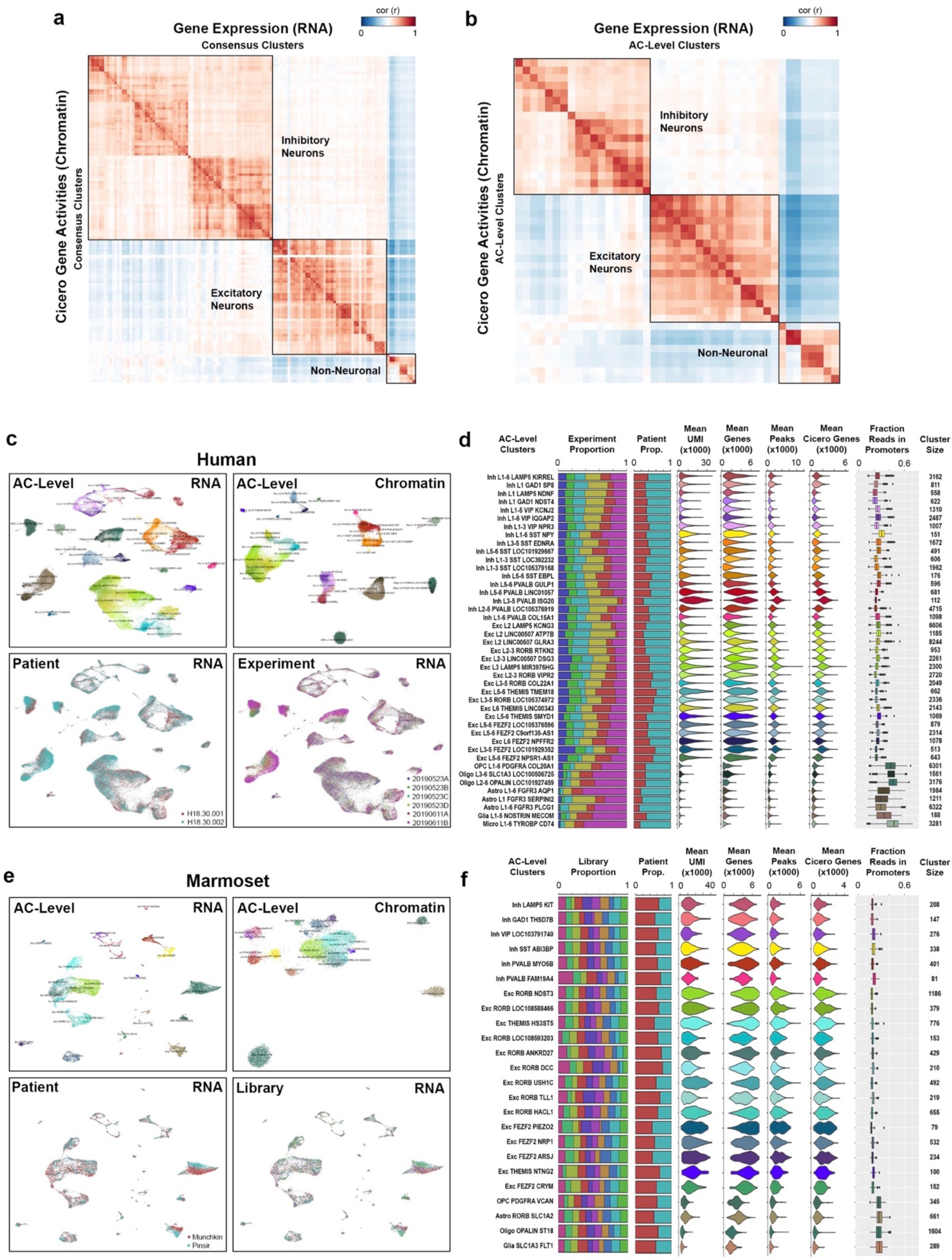
SNARE-Seq2 quality statistics. **a-b**, Correlation heatmaps of average scaled gene expression values against average scaled Cicero gene activity values for consensus clusters (**a**) and AC-level clusters (**b**). **c**, UMAP plots showing human AC-level clusters for both RNA and chromatin data, as well as the corresponding patient and experiment identities for the RNA embeddings. **d**, Bar, violin and box plots for human AC-level clusters showing proportion contributed by each experiment or patient, mean UMI and genes detected from the RNA data, the mean peaks and cicero active genes detected from AC data, the fraction of reads found in promoters for AC data, and the number of nuclei making up each of the clusters. **e**, UMAP plots showing marmoset AC-level clusters for both RNA and chromatin data, as well as the corresponding patient and library identities for the RNA embeddings. **f**, Bar, violin and box plots for marmoset AC-level clusters showing proportion contributed by each library or patient, mean UMI and genes detected from the RNA data, the mean peaks and cicero active genes detected from AC data, the fraction of reads found in promoters for AC data, and the number of nuclei making up each of the clusters.

**Extended Data Figure 7.**
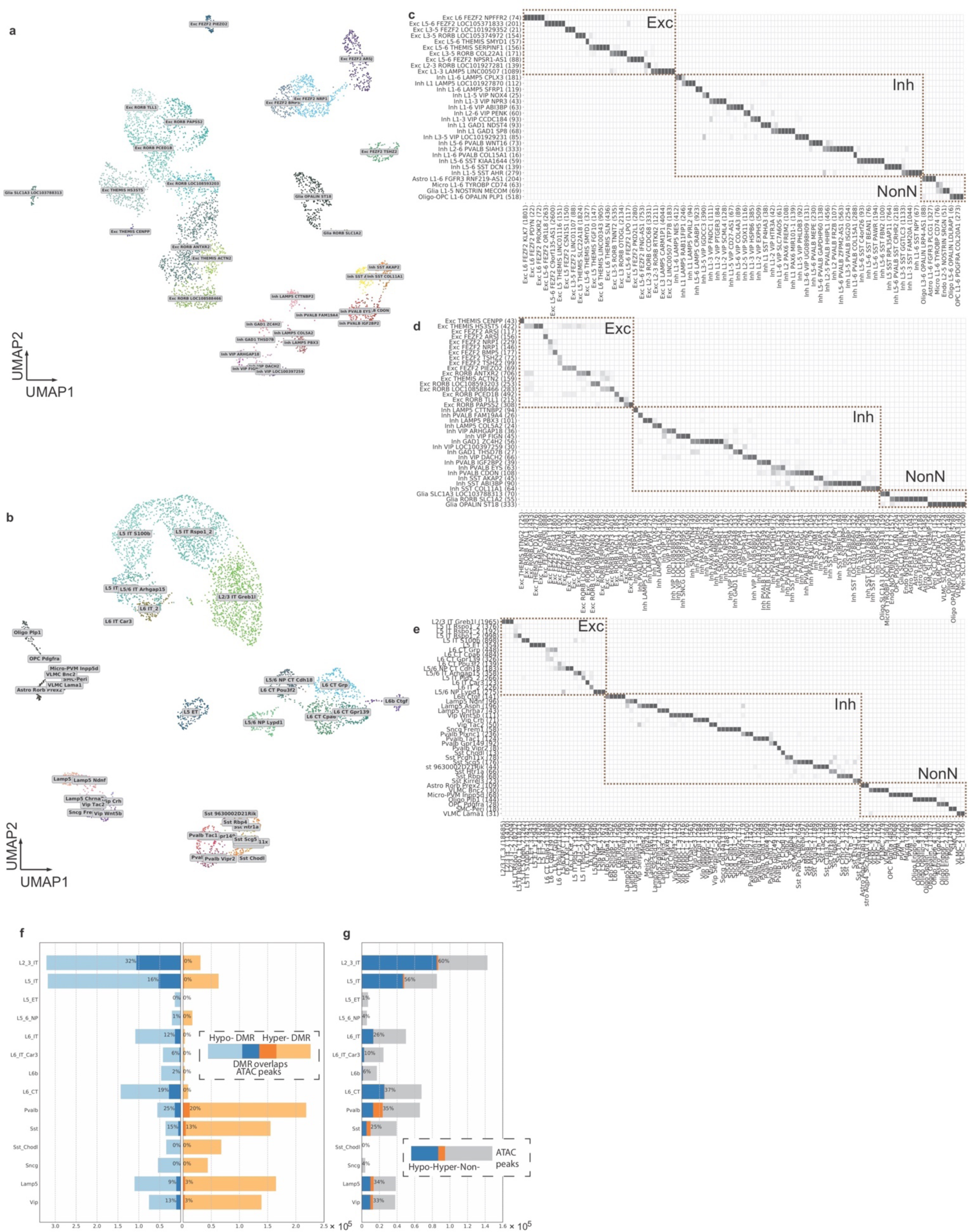
DNA-methylation cell type and integration with RNA-seq data. **a-b**, UMAP visualization of marmoset M1 and mouse MOp DNA methylation (snmC-seq2) data and cell clusters. **c-e**, Mapping between DNAm-seq and RNA-seq clusters from human (**c**), marmoset (**d**), and mouse (**e**). Number of nuclei in each cluster are listed in parentheses. **f**, Numbers of hypo- and hyper-methylated DMRs and overlap with chromatin accessible peaks in each subclass of human. **g**, Numbers of chromatin accessible peaks and overlap with DMRs in each subclass of human.

**Extended Data Figure 8.**
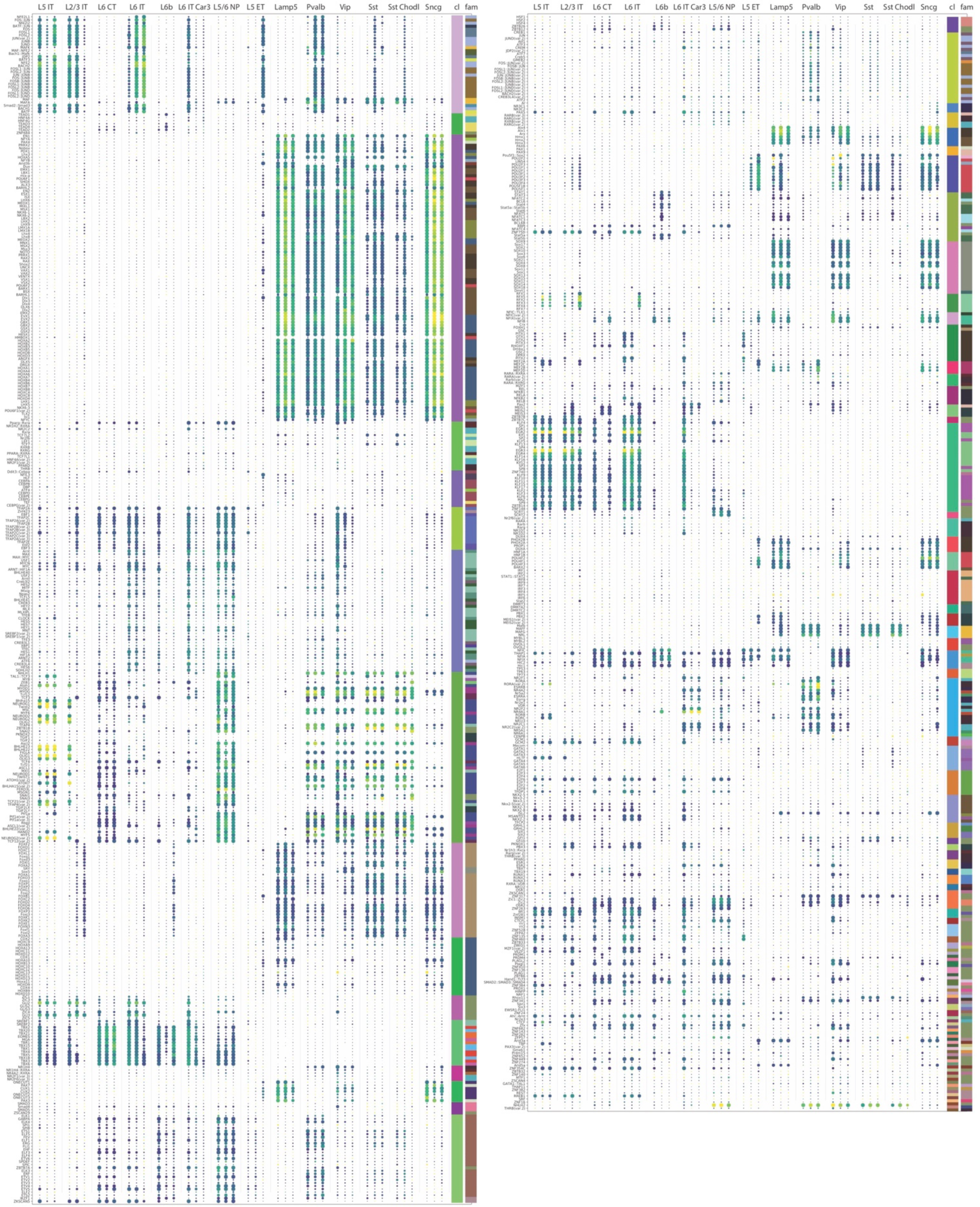
TFBS enrichment analysis on hypo-methylated DMRs at subclass level show conservativity of gene regulation across species. Motif enrichment analysis of TFBS were conducted using JASPAR’s non-redundant core vertebrata TF motifs for neuronal subclasses in each species. Each subclass tri-column shows the results of human, marmoset and mouse, respectively from left to right. The size of a dot denotes the p-value of the corresponding motif, while the color denotes the fold change. The rightmost two columns show TF clusters (cl) identified from motif profiles and TF family (fam) identified from TF structures.

**Extended Data Figure 9.**
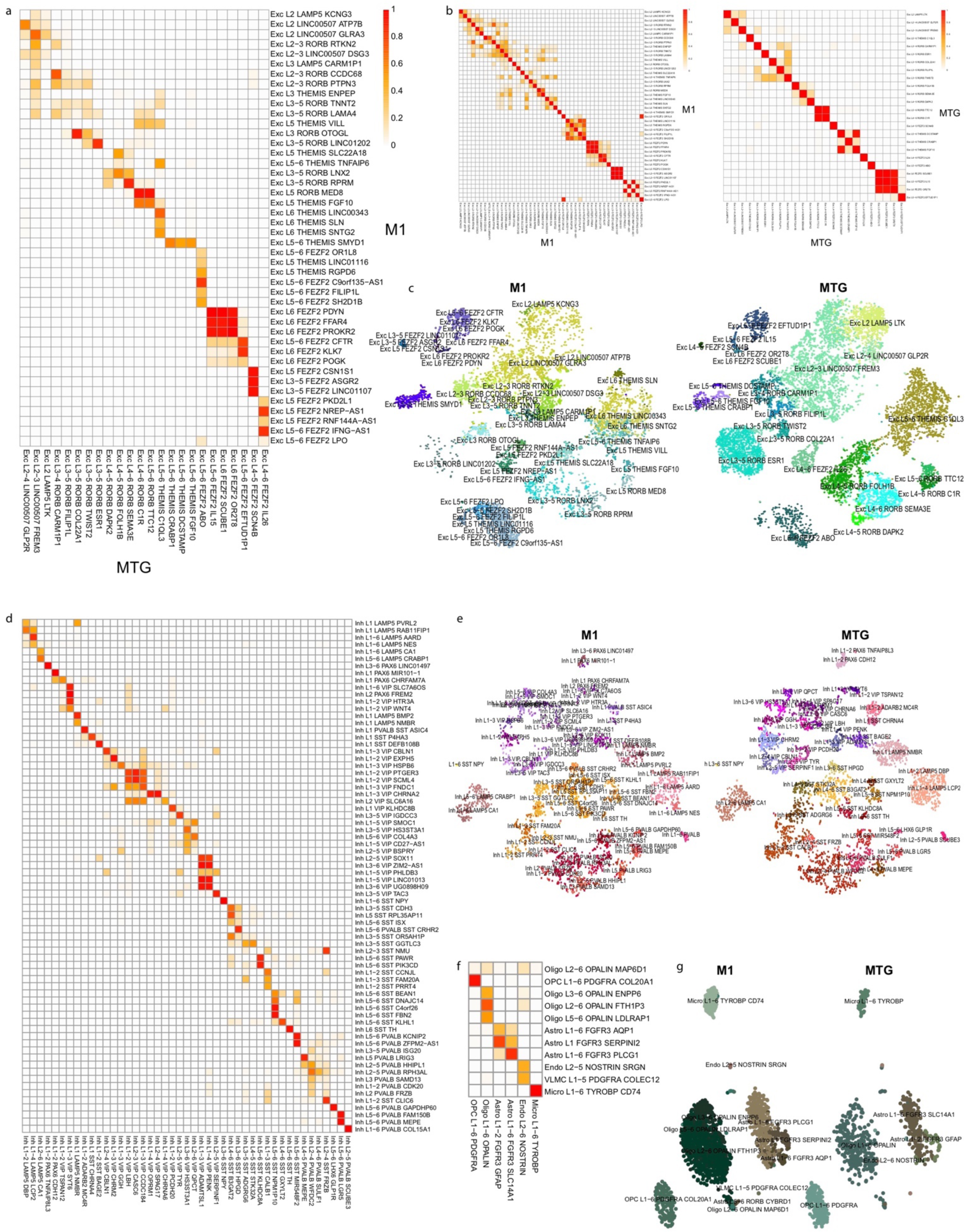
Cell type homologies between human cortical areas based on RNA-seq integration. **a**, Heatmap of glutamatergic neuron cluster overlap between M1 and MTG. **b**, Heatmaps of glutamatergic neuron cluster overlap for M1 and MTG test datasets. Clusters were split in half and two datasets were integrated using the same analysis pipeline as the M1 and MTG integration. Most clusters mapped correctly (along the diagonal) with some loss in resolution between closely related clusters (red blocks). **c**, tSNE plots of integrated glutamatergic neurons labeled with M1 and MTG clusters. **d-g**, Cluster overlap heatmaps and tSNE plots of integrations of GABAergic neurons (**d**, **e**) and non-neuronal cells (**f**, **g**), as described for glutamatergic neurons.

**Extended Data Figure 10.**
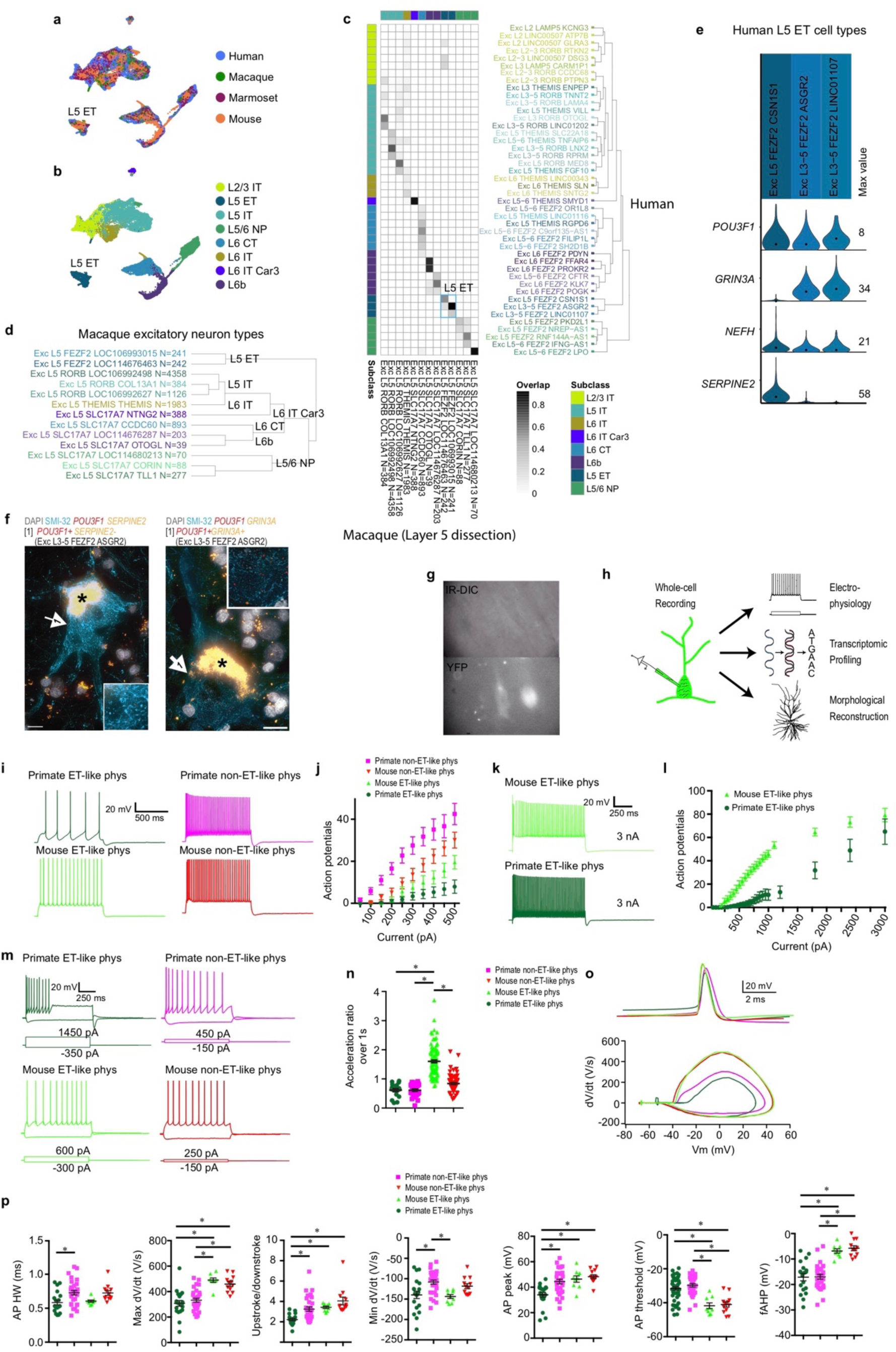
Cross-species alignment of glutamatergic neurons and differences in L5 neuron spike trains and single spike properties. **a, b**, UMAP visualizations of cross-species integration of snRNA-seq data for glutamatergic neurons isolated from human, macaque (L5 dissection only), marmoset, and mouse. Colors indicate species (**a**) or cell subclass (**b**). **c**, Cluster overlap heatmap showing the proportion of nuclei from within-species clusters that are mixed within the same integrated clusters. Human clusters (rows) are ordered by the dendrogram reproduced from **Figure 1c**. Macaque clusters (columns) are ordered to align with human clusters. Color bars at top and left indicate subclasses of within-species clusters. Blue box denotes the L5 ET subclass. **d**, Dendrogram showing all macaque clusters from L5 dissection with subclasses denoted to the right. **e**, Violin plot showing expression of marker genes for human L5 ET neuron subtypes. **f**, Two examples of ISH labeled, SMI-32 IF stained Betz cells in L5 of human M1 that correspond to the L5 ET cluster Exc L3-5 *FEZF2 ASGR2*. Insets show higher magnification of ISH-labeled transcripts in corresponding cells. Scale bars, 20 µm. Asterisks mark lipofuscin. **g**, Example IR-DIC (top) and fluorescent (bottom) images obtained from a macaque organotypic slice culture. Note the inability to visualize the fluorescently labeled neurons in IR-DIC because of dense myelination. **h**, patch-seq involves the collection of morphological, physiological and transcriptomic data from the same neuron. Following electrophysiological recording and cell filling with biocytin via whole cell patch clamp, the contents of the cell are aspirated and processed for RNA-sequencing. This permits a transcriptomic cell type to be pinned on the physiologically-probed neuron. **i**, Example voltage responses to a 1 s, 500 pA step current injection. **j.** Action potentials as a function of current injection amplitude. Primate ET neurons display shallowest action potential-current injection relationship, perhaps partially because of their exceptionally low input resistance. **k**, Voltage responses to a 1 s, 3 nA step current injection. **l**, Action potentials as a function of current injection for a subset of experiments in which current injection amplitude was increased incrementally to 3 nA. While both mouse and primate ET neurons could sustain high firing rates, primate neurons required 3 nA of current over 1s to reach similar average firing rates as mouse ET neurons. **m**, Example voltage responses to 1 s depolarizing step current injections. The amplitude of the current injection was adjusted to produce ∼10 spikes. Also shown are voltage responses to a hyperpolarizing current injection. **n**, The firing rate of primate ET and IT neurons decreased during the 1 s step current injection, whereas, the firing rate of mouse ET neurons increased. Acceleration ratio=2^nd^/last interspike interval. **o**, Example single action potentials (above) and phase plane plots (below). **p**, Various action potential features are plotted as a function of cell type. Notably, action potentials in primate ET neurons were reminiscent of fast spiking interneurons in that they were shorter and more symmetrical compared with action potentials in other neuron types/species. Intriguingly, K^+^ channel subunits K_v_3.1 and K_v_3.2 that are implicated in fast spiking physiology^91^ are encoded by highly expressed genes (*KCNC1* and *KCNC2*) in primate ET neurons (Fig. 7c) * p < 0.05, Bonferroni corrected t-test.

**Extended Data Table 1.** Summary of human tissue donors. RIN, RNA integrity number. Data type: SMART-Seqv4 (SSv4), 10x Genomics Chromium Single Cell 3’ Kit v3 (Cv3), Single-Nucleus Chromatin Accessibility and mRNA Expression sequencing (SNARE-seq2), Single-nucleus methylcytosine sequencing (snmC-seq2).

| Specimen ID | Age | Sex | Race | Cause of Death | PMI (hr) | Tissue RIN | Hemisphere Sampled | Data Type |
| --- | --- | --- | --- | --- | --- | --- | --- | --- |
| H200.1023 | 43 | F | Iranian descent | Mitral valve prolapse | 18.5 | 7.4 ± 0.7 | L | SSv4 |
| H200.1025 | 50 | M | Caucasian | Cardiovascular | 24.5 | 7.6 ± 1.0 | L | SSv4 |
| H200.1030 | 54 | M | Caucasian | Cardiovascular | 25 | 7.7 ± 0.8 | L | SSv4 |
| H18.30.001 | 60 | F | Unknown | Car accident | 18 | 7.9 ± 2.5 | R | SSv4, Cv3, SNARE-seq2, sn-methylome |
| H18.30.002 | 50 | M | Unknown | Cardiovascular | 10 | 8.2 ± 0.4 | R | SSv4, Cv3, SNARE-seq2, snmC-seq2 |

**Extended Data Table 2.** Summary of marmoset specimens. Data type: 10x Genomics Chromium Single Cell 3’ Kit v3 (Cv3). ACD Bio multiplex fluorescent in situ hybridization (FISH).

| Specimen ID | Age (years) | Sex | Data Type |
| --- | --- | --- | --- |
| bi005 | 2.3 | M | Cv3 |
| bi006 | 3.1 | F | Cv3 |
| bi003 | 1.9 | M | FISH |

